# *k*-mer-based GWAS enhances the discovery of causal variants and candidate genes in soybean

**DOI:** 10.1101/2023.03.28.534607

**Authors:** Marc-André Lemay, Maxime de Ronne, Richard Bélanger, François Belzile

## Abstract

Genome-wide association studies (GWAS) are powerful statistical methods that detect associations between genotype and phenotype at genome scale. Despite their power, GWAS frequently fail to pinpoint the causal variant or the gene controlling a trait at a given locus in crop species. Assessing genetic variants beyond single-nucleotide polymorphisms (SNPs) could alleviate this problem, for example by including structural variants (SVs). In this study, we tested the potential of SV-and *k*-mer-based GWAS in soybean by applying these methods to 13 traits. We also performed conventional GWAS analysis based on SNPs and small indels for comparison. We assessed the performance of each GWAS approach based on results at loci for which the causal genes or variants were known from previous genetic studies. We found that *k*-mer-based GWAS was the most versatile approach and the best at pinpointing causal variants or candidate genes based on the most significantly associated *k*-mers. Moreover, *k*-mer-based analyses identified promising candidate genes for loci related to pod color, pubescence form, and resistance to the oomycete *Phytophthora sojae*. In our dataset, SV-based GWAS did not add value compared to *k*-mer-based GWAS and may not be worth the time and computational resources required to genotype SVs at population scale. Despite promising results, significant challenges remain regarding the downstream analysis of *k*-mer-based GWAS. Notably, better methods are needed to associate significant *k*-mers with sequence variation. Together, our results suggest that coupling *k*-mer-and SNP/indel-based GWAS is a powerful approach for discovering candidate genes in crop species.

## Introduction

Genome-wide association studies (GWAS) are analytical approaches that detect statistical associations between phenotypic observations for a trait of interest and the genotypes of variants distributed throughout the genome (Tam et al., 2019). This approach has been used for over 15 years (Visscher et al., 2012) and has enabled significant advances in our understanding of the genetic architecture of traits involved in human health (e.g. The Wellcome Trust Case Control Consortium, 2007) and crop (Tibbs Cortes et al., 2021) and animal (Ma et al., 2019) production, among others.

Despite their power and convenience for finding significantly associated loci, GWAS analyses frequently fail to pinpoint the gene(s) associated with a trait and the causal variant(s) involved (Tam et al., 2019). One possible reason for this is that many GWAS analyses do not include the causal variant itself in their variant dataset (Tibbs Cortes et al., 2021). This could be either because the analysis is restricted to a particular type of variant, such as single-nucleotide polymorphisms (SNPs), or because the set of markers/variants genotyped in the association panel is limited by the technology being used (e.g. Bandillo et al., 2015; Sonah et al., 2015). Despite this limitation, variants located near the gene controlling the trait often do appear statistically associated with the phenotype through linkage disequilibrium (LD) with the causal variant (Korte and Farlow, 2013). A typical workflow then involves the identification of candidate genes within the haplotype blocks defined by the statistically associated markers and follow-up functional analyses to confirm that a given gene controls the trait (e.g. Wang et al., 2018; Liu et al., 2020a).

The inclusion of structural variants (SVs) as genotype data in GWAS may partly overcome this issue by providing a set of variants with a potentially high functional impact. SVs include any variant involving a difference of at least 50 nucleotides, such as deletions, insertions, or inversions (Ho et al., 2020). Such variants are known to have large phenotypic impacts by disrupting coding or regulatory sequences (Marroni et al., 2014), and have been identified as causal variants in traits such as resistance to soybean cyst nematode (Cook et al., 2012), aluminum tolerance in wheat (Maron et al., 2013), and branching in maize (Studer et al., 2011). SVs have already been used in GWAS with promising results (e.g. Zhang et al., 2015; Akakpo et al., 2020; Domínguez et al., 2020; Liu et al., 2020b), but their use in GWAS is the exception rather than the norm.

Despite their phenotypic impact, SVs are still largely understudied compared to SNPs because discovering and genotyping them with accuracy is difficult. Comprehensive detection of SVs requires a combination of several methods that often need to be tailored to the study species and data available (Alkan et al., 2011; Ho et al., 2020). Even then, assessments of the performance of SV discovery and genotyping generally reveal subpar sensitivity and precision (e.g. Cameron et al., 2019; Chaisson et al., 2019; Kosugi et al., 2019). Moreover, despite recent improvements in computational approaches for genotyping SVs (e.g. Sirén et al., 2021; Ebler et al., 2022), such approaches remain relatively inaccurate compared to SNP genotyping workflows.

Alternative approaches that allow one to assess a large spectrum of variants in GWAS without genotyping SVs at population scale could be useful. Voichek and Weigel (2020) developed a *k*-mer-based GWAS approach that may represent such an alternative method. This approach relies on the presence or absence of *k*-mers observed in sequence reads in place of variant genotypes for use in GWAS. Once a presence/absence table of *k*-mers has been generated for the population under study, this table can be used in association analyses similarly to SNP or SV genotype calls. One advantage of using *k*-mers is that they can act as a molecular signature for any type of variant as long as these variants result in presence/absence of *k*-mers of a given length. Another advantage is that *k*-mers are not tied to a specific genomic location and can thus be used to query associations between genotype and phenotype at genomic locations that are not included in a reference genome. A few studies have applied this *k*-mer-based GWAS approach (e.g. Tripodi et al., 2021; Colque-Little et al., 2021) and other similar approaches have been developed (Rahman et al., 2018; He et al., 2021), but similarly to the use of SVs in GWAS, such approaches have not been widely adopted yet.

In this study, we assessed the potential of SV-and *k*-mer-based GWAS in a major crop, soybean (*Glycine max*). We also conducted conventional GWAS based on SNPs and indels. In order to test these methods, we analyzed ten qualitative traits (Bandillo et al., 2017) and two quantitative traits (Bandillo et al., 2015) that have been previously studied using a SNP array developed for soybean (Song et al., 2013). In addition, we analyzed a quantitative trait (horizontal resistance to *Phytophthora sojae*) that has been previously studied using whole-genome sequencing (WGS) data (de Ronne et al., 2022). Since the underlying genes and causal variants are already known for several of these traits, they provide a good test case for the performance of GWAS methods. The objectives of our study were to:

1. Assess the potential of SV-and *k*-mer-based GWAS approaches to pinpoint genes and causal variants associated with loci whose underlying genes or variants are already known.
2. Identify potential candidate genes and/or causal variants at loci whose underlying genes or candidate variants are not yet known.
3. Develop a set of computational tools for the downstream analysis of significantly associated *k*-mers once these have been identified.

## Results and discussion

We analyzed a total of 13 traits in a population of 363 resequenced *G. max* accessions using three GWAS approaches that differed in the genotypes used. The first approach used SNP and indel genotypes called by Platypus. Despite both SNPs and indels being used, we will refer to this approach as SNP-based for simplicity. Our SV-based workflow used various sources of data for SV discovery and a single program (Paragraph) for genotyping SVs at population scale from Illumina WGS data. Finally, the third GWAS approach relied on the presence/absence of *k*-mers in the WGS data as genotypes. Because of the large number of *k*-mers analyzed, we only report results for those that were considered significantly associated with traits of interest.

Since two or three different GWAS analyses were conducted for some of the traits, a total of 22 GWAS analyses were performed using each of the three approaches. Given the large volume of data that this represents, we focus here on the most noteworthy results. Readers interested in results at loci that are not discussed here are referred to Additional file 1. In particular, Table S1 (Additional file 1) provides a summary of the *p*-values obtained by each approach at each of the loci considered in this study.

### Loci with known genes or causal variants

We first wanted to assess the performance of the various GWAS approaches by analyzing the results at loci for which the genes have already been cloned. These loci provide interesting test cases because the expected results (causal genes or variants) are known. Table 1 shows a summary of the results discussed below.

**Table 1:** Performance of three GWAS approaches at detecting causal genes or variants at cloned loci.

| Trait | Locus | Gene | Causal variant | SNPs<sup>a</sup> | SVs<sup>b</sup> | *k*-mers |
| --- | --- | --- | --- | --- | --- | --- |
| Flower color | *W1* | Glyma.13g072100 | 65-bp deletion (Zabala and Vodkin, 2007) | ** | *** | *** |
| Pubescence color | *T* | Glyma.06g202300 | 1-bp deletion (Zabala and Vodkin, 2003) | ** | * | ** |
| Pubescence color | *Td* | Glyma.03g258700 | SNP (Yan et al., 2020) | * | - | *** |
| Seed coat color | *I* | CHS gene cluster | 10.9-kb inversion/duplication (Tuteja and Vodkin, 2008) | ** | ** | ** |
| Seed coat color | *G* | Glyma.01g198500 | SNP (Wang et al., 2018) | ** | ** | *** |
| Seed coat color<sup>c</sup> | *T* | Glyma.06g202300 | 1-bp deletion (Zabala and Vodkin, 2003) | - | - | - |
| Seed coat color<sup>c</sup> | *R* | Glyma.09g235100 | SNPs and indels (Gillman et al., 2011) | - | - | - |
| Stem termination type | *Dt1* | Glyma.19g194300 | SNPs (Liu et al., 2010) | ** | * | ** |
| Stem termination type<sup>c</sup> | *Dt2* | Glyma.18g273600 | SNPs (Ping et al., 2014) | - | - | - |
| Stem termination type<sup>c</sup> | *E3* | Glyma.19g224200 | Various types (Tardivel et al., 2019) | - | - | - |
| Hilum color<sup>c</sup> | *T* | Glyma.06g202300 | 1-bp deletion (Zabala and Vodkin, 2003) | ** | ** | ** |
| Hilum color<sup>c</sup> | *I* | CHS gene cluster | 10.9-kb inversion/duplication (Tuteja and Vodkin, 2008) | * | * | ** |
| Hilum color | *R* | Glyma.09g235100 | SNPs and indels (Gillman et al., 2011) | * | - | ** |
| Hilum color<sup>c</sup> | *W1* | Glyma.13g072100 | 65-bp deletion (Zabala and Vodkin, 2007) | - | - | - |
| Pubescence density | *Ps* | Glyma.12g187200 | 25.6-kb CNV (Liu et al., 2020a) | ** | ** | *** |
| Pubescence density<sup>c</sup> | *Pd1* | Glyma.01g240100 | SNP (Liu et al., 2020a) | - | - | - |
| Pubescence density<sup>c</sup> | *P1* | Glyma.09g278000 | SNP (Liu et al., 2020a) | - | - | - |
| Seed coat luster | *B* | Duplicated HPS genes | CNV (several kb) (Gijzen et al., 2006) | ** | ** | ** |
| Seed coat luster<sup>c</sup> | *I* | CHS gene cluster | 10.9-kb inversion/duplication (Tuteja and Vodkin, 2008) | - | - | - |
| Maturity group<sup>c</sup> | *E1* | Glyma.06g207800 | Various types (Tardivel et al., 2019) | - | - | - |
| Maturity group<sup>c</sup> | *E2* | Glyma.10g221500 | Various types (Tardivel et al., 2019) | - | - | - |
| Maturity group<sup>c</sup> | *E3* | Glyma.19g224200 | Various types (Tardivel et al., 2019) | - | - | - |
| Maturity group<sup>c</sup> | *E4* | Glyma.20g090000 | Various types (Tardivel et al., 2019) | - | - | - |
| Oil<sup>c</sup> | *oilGm20* | Glyma.20g085100 | 321-bp deletion (Fliege et al., 2022) | - | - | - |
| Oil<sup>c</sup> | *oilGm15* | Glyma.15g049200 | SNPs and indels (Zhang et al., 2020) | - | - | - |
| Protein<sup>c</sup> | *proGm20* | Glyma.20g085100 | 321-bp deletion (Fliege et al., 2022) | - | - | - |
| Protein<sup>c</sup> | *proGm15* | Glyma.15g049200 | SNPs and indels (Zhang et al., 2020) | - | - | - |

<sup>a</sup> SNPs and indels genotyped by Platypus<sup>b</sup> SVs genotyped by Paragraph<sup>c</sup> Locus not discussed in main text. See Additional file 1 for detailed results.
- The approach detected no signal overlapping the gene.
\* The signal found at this locus overlaps the gene.
\*\* The region defined by the top 5% (SVs) or 1% (SNPs and *k*-mers) variants overlaps the gene.
\*\*\* The most significantly associated variant is the causal variant.

#### Flower color – *W1* locus

Zabala and Vodkin (2007) identified the flavonoid 3’5’-hydroxylase (F3’5’H) gene Glyma.13g-072100 as associated with the *W1* locus for flower color. In Williams82 and other accessions with white flowers, a 65-bp insertion into the third exon of the gene results in a premature stop codon that renders the F3’5’H enzyme non-functional and prevents pigmentation of the flower. All GWAS approaches detected a signal overlapping the gene at this locus (Additional file 1: Figures S1 and S2). The most significant SV and *k*-mers corresponded to the known causal variant (Figure 1a-b and Additional file 1: S3c-d). In particular, the assembly of the reads containing the most significant *k*-mers and the alignment of the resulting haplotypes showed that these *k*-mers directly tagged the causal variant (Figure 1c-e). As the catalogue of variants analyzed by Platypus contains only SNPs and small indels, the causal variant could not be captured by this approach. Still, SNPs and indels located within the gene sequence were among the most significantly associated ones (Additional file 1: Figure S3b).

**Figure 1:**
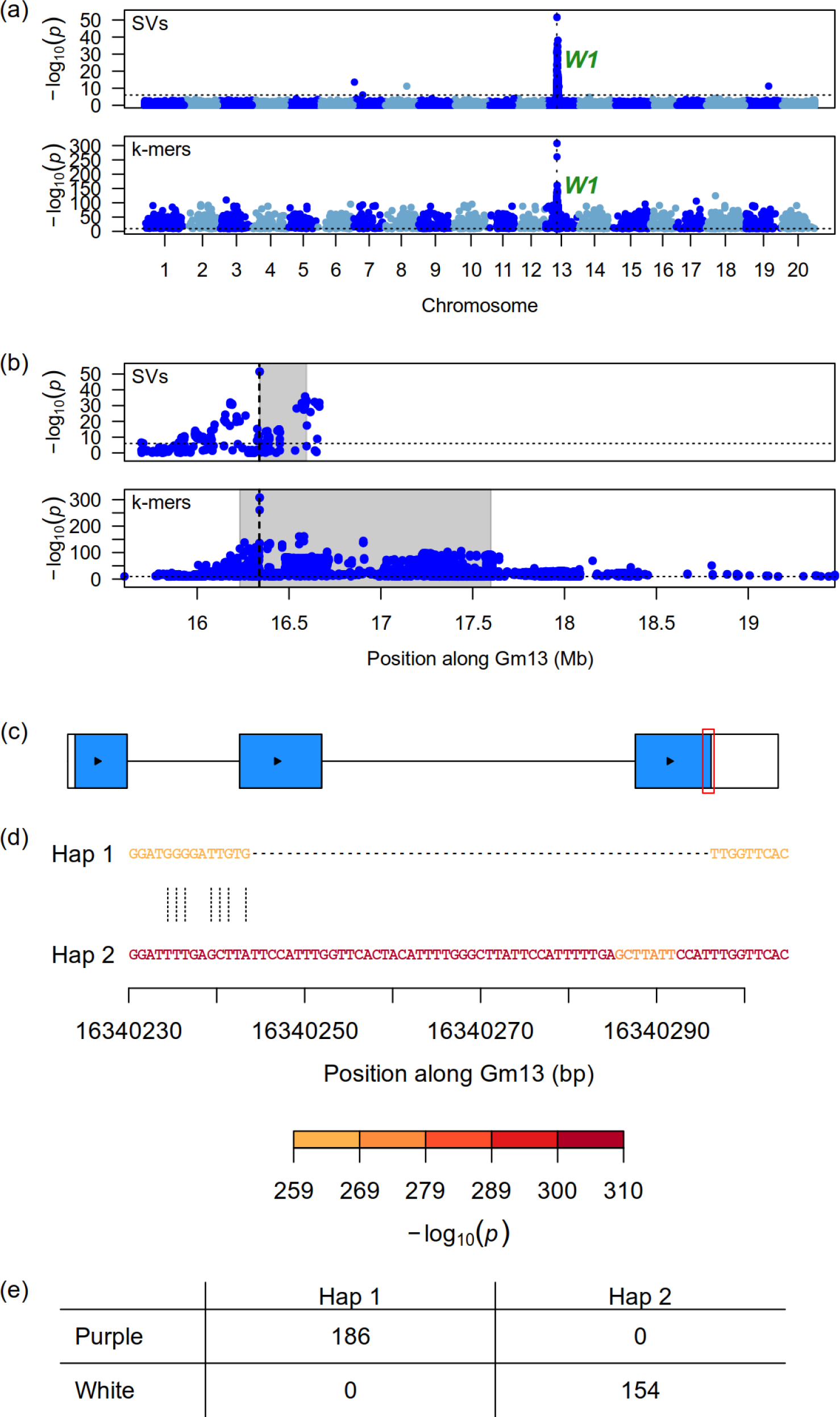
Results of SV-and *k*-mer-based GWAS at the *W1* locus for flower color. (a) Genome-wide Manhattan plots. Horizontal dotted lines indicate the significance threshold whereas vertical dotted lines indicate the location of the *W1* locus. (b) Zoomed-in Manhattan plots at the *W1* locus. The vertical dotted lines indicate the location of the Glyma.13g072100 gene associated with this locus. The shaded gray rectangles show the regions delimited by the top 5% (SVs) and 1% (*k*-mers) associations. (c) Gene model of Glyma.13g072100. Rectangles represent exons, colored rectangles represent coding sequences, and arrows indicate the direction of transcription. The red rectangle highlights the region shown in panel (d). (d) Pairwise alignment of the two haplotype sequences in the population at the *W1* locus. Nucleotides are colored according to the *p*-value of the most significant *k*-mer overlapping them. Vertical dotted lines between haplotypes indicate sequence differences, whereas dashes indicate gaps in the alignment. (e) Contingency table of the phenotypes and haplotypes found in the population at the *W1* locus.

The *k*-mer GWAS analysis also identified presumably spurious significant *k*-mers scattered throughout the genome (Additional file 1: Figure S1c). An analysis of the pairwise LD between a subset of those *k*-mers showed that most *k*-mers formed a single LD block linked to the *W1* locus (Additional file 1: Figure S4). A small LD block occurred on chromosome Gm09, however it was in moderate LD with the one on chromosome Gm13 and therefore probably does not represent a separate locus.

We should note that when we first analyzed the results of the GWAS on flower color, we observed that two accessions bearing the haplotype typically associated with purple flowers were noted as having white flowers. Upon further inspection, we found that the most likely cause of this discrepancy was a mismatch between the genotype data and the identity of the accession for which the phenotypic data was obtained. Indeed, when we compared SNP calls made directly from the WGS data to those made using the SoySNP50K array, we observed that these two accessions had low concordance rates of only 62% and 70%. To ensure that such mismatches would not affect our GWAS analyses, we computed the concordance between genotype calls made from WGS data and the SoySNP50K array for all samples (see the Methods) and excluded 24 accessions for which the concordance rate was below 90%.

#### Pubescence color – *T* locus

Zabala and Vodkin (2003) identified Glyma.06g202300 as the gene associated with the *T* locus for pubescence color and found a 1-bp deletion as a putative causal variant at this locus. For all three GWAS approaches, we found a signal overlapping this gene (Additional file 1: Figure S5). In particular, the regions defined by the 1% most associated variants identified by the SNP-and *k*-mer-based approaches included Glyma.06g202300 (Additional file 1: Figure S6). While the most significant *k*-mers did occur within the sequence of this gene, they did not correspond to the documented causal variant (Additional file 1: Figure S7d). Still, the fourth and fifth most strongly associated *k*-mers corresponded to this causal variant (Additional file 1: Figures S7d and S8). We observed that not all accessions with gray pubescence harbored the deleted nucleotide (Additional file 1: Figure S8c), which could indicate either that other causal variants are resulting in a non-functional F3’H in this population, or that the causal variant lies elsewhere. As for SNP-based GWAS, the most significant variant within the sequence of Glyma.06g202300 was the causal variant, however it was not the most significantly associated variant overall (Additional file 1: Figures S6b and S7b). SV-based GWAS did not detect the causal variant at this locus, as expected from its size.

#### Pubescence color – *Td* locus

We conducted a separate GWAS analysis contrasting only accessions with tawny and light tawny pubescence because the *Td* locus is known to control these differences in color. Accordingly, we detected signals in this region using all three approaches, but only SNP-and *k*-mer-based GWAS identified a signal that overlapped Glyma.03g258700 (Figure 2a-b, and Additional file 1: Figures S9 and S10). While the most significant *k*-mer identified the causal variant documented at this locus (Yan et al., 2020), it was the fourth most significantly as-sociated variant for the SNP-based approach (Figure 2b-d, Additional file 1: Figure S11). Interestingly, some of the accessions bearing the haplotype associated with tawny pubescence displayed light tawny pubescence, suggesting that there may be more than one causal variant at this locus (Figure 2e).

**Figure 2:**
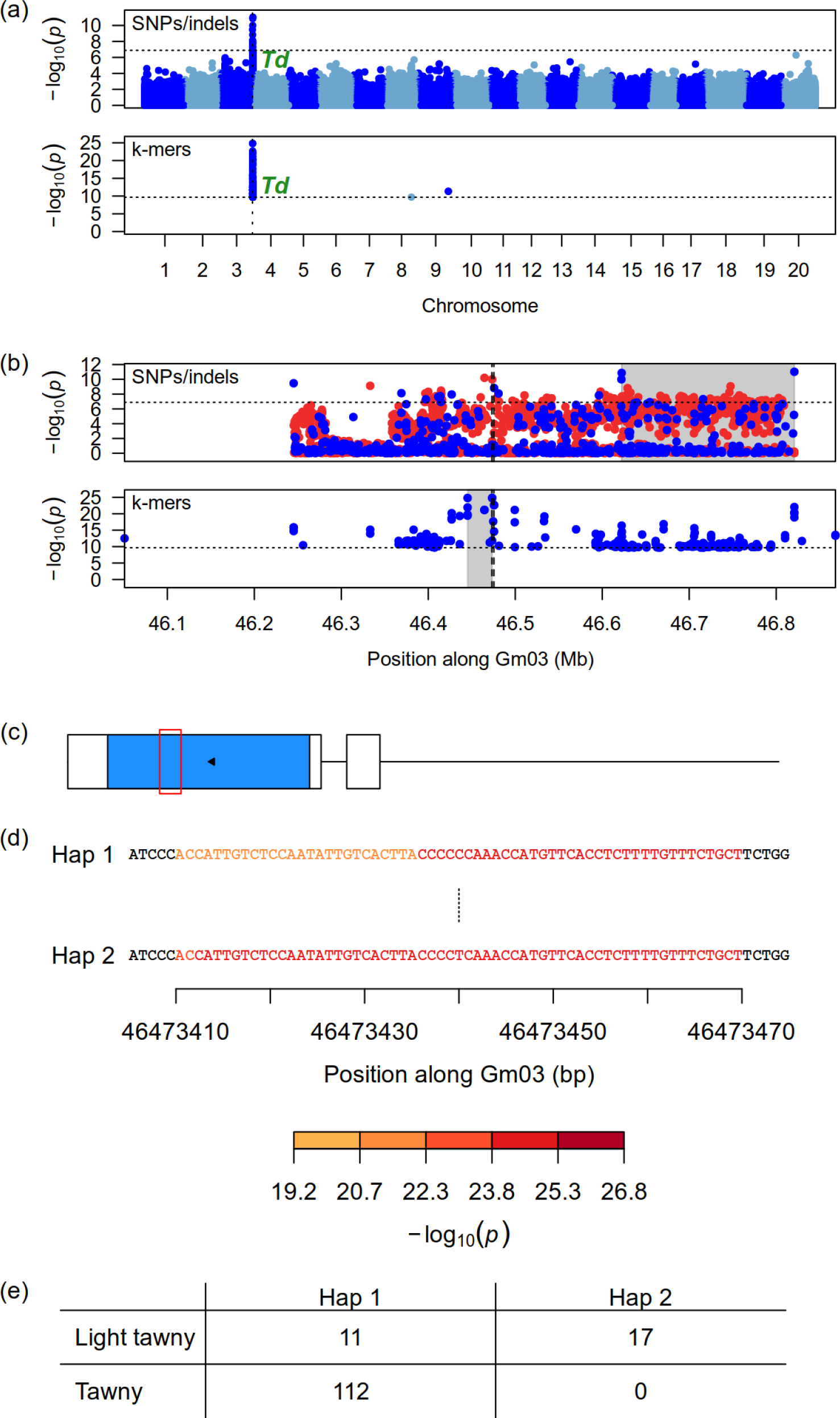
Results of SNP-and *k*-mer-based GWAS at the *Td* locus for pubescence color. (a) Genome-wide Manhattan plots. Horizontal dotted lines indicate the significance threshold whereas vertical dotted lines indicate the location of the *Td* locus. (b) Zoomed-in Manhattan plots at the *Td* locus. The vertical dotted lines indicate the location of the Glyma.03g258700 gene associated with this locus. The shaded gray rectangles show the regions delimited by the top 1% associations. In the case of SNPs, blue points denote markers used in the original analysis, whereas red points denote markers that had originally been pruned but whose *p*-values were computed after signal discovery. (c) Gene model of Glyma.03g258700. Rectangles represent exons, colored rectangles represent coding sequences, and the arrow indicates the direction of transcription. The red rectangle highlights the region shown in panel (d). (d) Pairwise alignment of the two haplotype sequences in the population at the *Td* locus. Nucleotides are colored according to the *p*-value of the most significant *k*-mer overlapping them. The vertical dotted line between haplotypes indicates the location of the causal SNP at this locus. (e) Contingency table of the phenotypes and haplotypes found in the population at the *Td* locus.

#### Seed coat color - *I* locus

The causal variant at the *I* locus for seed coat color is a complex SV that takes the form of a 10.91-kb inverted duplication of three chalcone synthase (CHS) genes (CHS1, CHS3 and CHS4; Tuteja and Vodkin, 2008). The presence of this inverted duplication results in the silencing of all CHS genes in the seed coat specifically and thus in the absence of seed coat pigmentation (Tuteja et al., 2009). While we did find signals overlapping this causal variant using all three GWAS approaches (Additional file 1: Figures S12 and S13), none of the approaches identified it. This failure to detect the causal variant is likely due to its complexity, which makes it difficult to detect and genotype it. While it is likely that at least some of the significantly associated *k*-mers actually are derived from this causal variant, the lack of systematic methods for linking *k*-mers to sequence variation hindered our ability to do so.

#### Seed coat color - *G* locus

We conducted a second GWAS analysis contrasting accessions with yellow and green seed coats in order to target the *G* locus. Accordingly, this analysis found strong signals overlapping the *G* locus using all three approaches (Additional file 1: Figure S14). Wang et al. (2018) identified Glyma.01g198500 as the gene associated with this locus and found an A>G SNP affecting transcript splicing as the causal variant. Consistently with their results, we identified the two most significant *k*-mers and the second most significant marker from the SNP-based analysis as corresponding to that causal variant (Figure 3 and Additional file 1: Figures S15 and S16). Unsurprisingly, SV-based GWAS did not identify the causal variant given its type.

**Figure 3:**
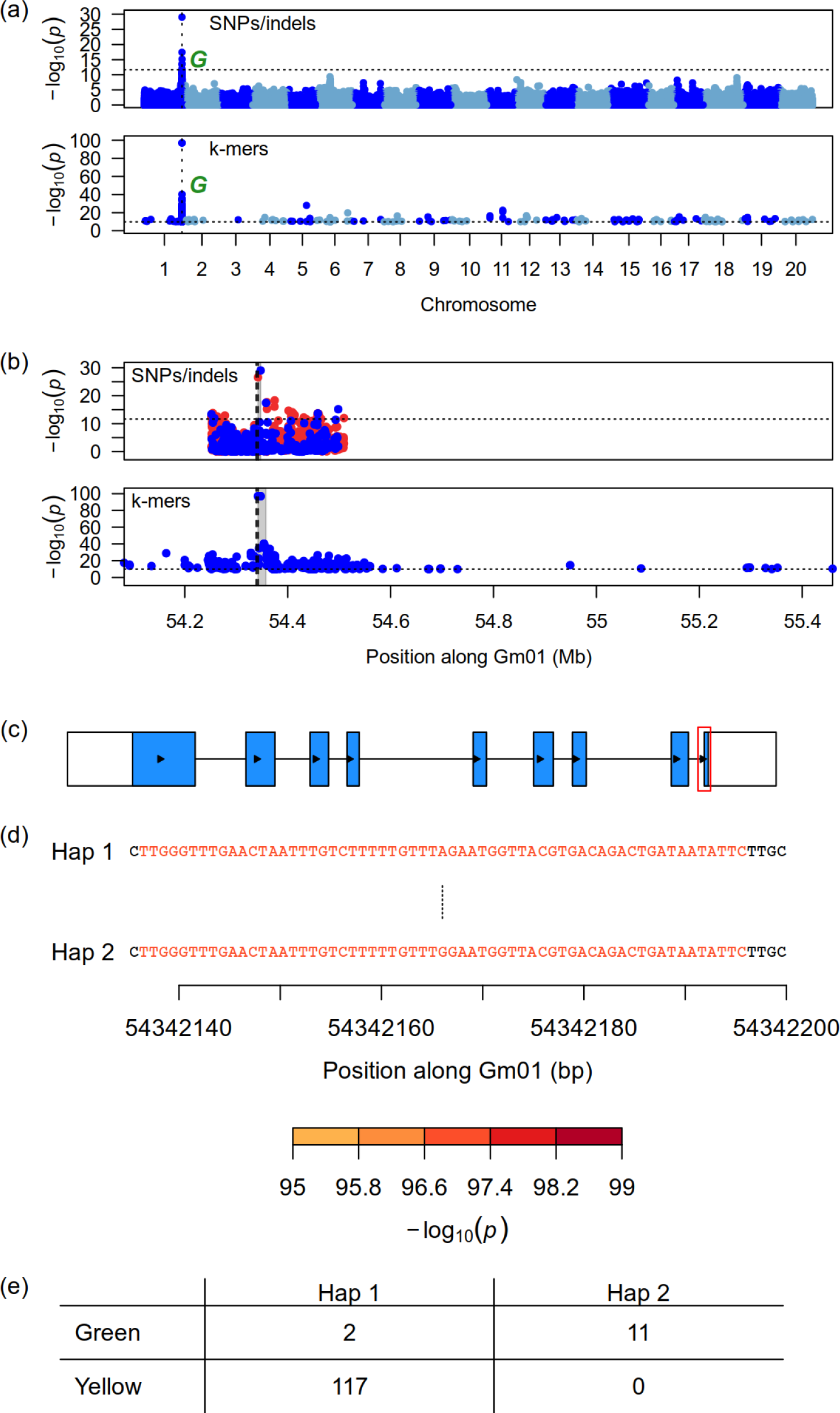
Results of SNP-and *k*-mer-based GWAS at the *G* locus for seed coat color. (a) Genome-wide Manhattan plots. Horizontal dotted lines indicate the significance threshold whereas vertical dotted lines indicate the location of the *G* locus. (b) Zoomed-in Manhattan plots at the *G* locus. The vertical dotted lines indicate the location of the Glyma.01g198500 gene associated with this locus. The shaded gray rectangles show the regions delimited by the top 1% associations. In the case of SNPs, blue points denote markers used in the original analysis, whereas red points denote markers that had originally been pruned but whose *p*-values were computed after signal discovery. (c) Gene model of Glyma.01g198500. Rectangles represent exons, colored rectangles represent coding sequences, and the arrows indicate the direction of transcription. The red rectangle highlights the region shown in panel (d). (d) Pairwise alignment of the two haplotype sequences in the population at the *G* locus. Nucleotides are colored according to the *p*-value of the most significant *k*-mer overlapping them. The vertical dotted line between haplotypes indicates the location of the causal SNP at this locus. (e) Contingency table of the phenotypes and haplotypes found in the population at the *G* locus.

#### Stem termination type - *Dt1* locus

While all GWAS approaches found signals near the Glyma.19g194300 gene associated with the *Dt1* locus for stem termination type, the SNP-and *k*-mer-based analyses performed best since the region defined by their most highly associated markers (top 1%) included the causal gene (Additional file 1: Figures S17 and S18). We did identify one of the SNPs suggested as putative causal variants for this locus by Tian et al. (2010) and Liu et al. (2010), however this SNP was only the 135^th^ most associated marker found by Platypus and the 12^th^ most associated *k*-mer (Additional file 1: Figures S19 and S20). Our results are consistent with previous findings in suggesting that there is more than one causal variant at the *Dt1* locus (Liu et al., 2010; Tian et al., 2010). Indeed, a few cultivars bearing the allele associated with *dt1* showed indeterminate or semi-determinate phenotypes and some cultivars showed a determinate phenotype without bearing the allele associated with *dt1* (Additional file 1: Figure S20c). The existence of several causal variants might explain why our analyses did not identify a causal variant more clearly.

#### Hilum color – *R* locus

We conducted a GWAS analysis that only compared accessions with black and brown hila in an attempt to detect stronger signals at the *R* locus. We accordingly detected a signal overlapping this locus using all approaches, but only SNP-and *k*-mer-based GWAS detected signals overlapping the Glyma.09g235100 gene associated with *R* (Additional file 1: Figures S21 and S22). Gillman et al. (2011) identified this R2R3 MYB transcription factor as the molecular basis for the *R* locus and documented four different loss-of-function alleles linked to brown hilum color. Although our analyses did detect strong signals in the vicinity of this gene, the most associated markers were not located in the body of the gene (Additional file 1: Figure S22). Interestingly, the most significantly associated *k*-mers detected by our analysis mapped 2 kb upstream of another gene (Glyma.09g234900) annotated as a MYB transcription factor and putatively involved in the regulation of anthocyanin biosynthesis. This gene as well as Glyma.09g235100 and two others are part of a tandem array of four R2R3 MYB genes considered as candidates by Gillman et al. (2011), and of which Glyma.09g235100 was the only one expressed in the seed coat. Given the convincing evidence provided by Gillman et al. (2011), it is most likely that Glyma.09g235100 is indeed the gene associated with the *R* locus and that our failure to identify the most significantly associated *k*-mers as causal variants is due to the coexistence of several variants that can cause loss of function at the *R* locus. Still, we observed two causal variants previously documented by Gillman et al. (2011) in our dataset (Additional file 1: Figures S23 and S24).

#### Pubescence density – *Ps* locus

We conducted a GWAS analysis comparing accessions with normal and semi-sparse pubescence in order to detect loci associated with pubescence density. This analysis detected signals at the *Ps* locus using all three approaches (Figure 4a-b and Additional file 1: Figures S25, S26 and S27). The causal variant at this recently cloned locus is a copy number variant (CNV) overlapping the Glyma.12g187200 gene, with higher copy number resulting in reduced pubescence (Liu et al., 2020a). We identified a region of *∼*42 kb that varied in copy number across the population and was associated with the trait (Figure 4c), and therefore possibly corresponded to the 25.6-kb region identified as the causal CNV by Liu et al. (2020a). We found that reads containing the most significantly associated *k*-mer mapped to either of the two ends of the CNV (Figure 4d). This suggests that the most associated *k*-mer is indeed linked to the CNV that is causal at this locus, while *k*-mers within the CNV region may represent differences in sequence between the copies.

**Figure 4:**
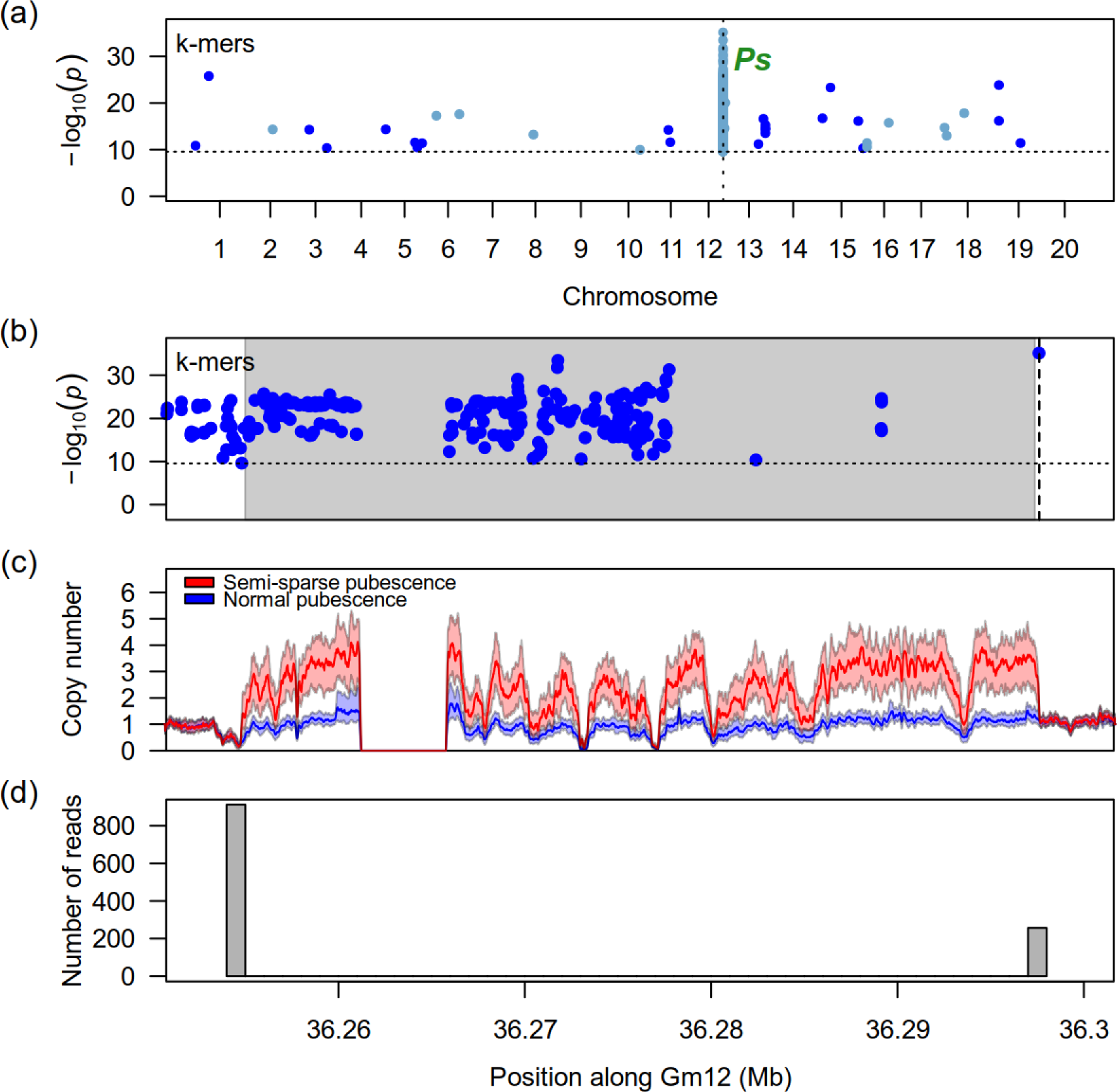
Results of *k*-mer-based GWAS at the *Ps* locus for pubescence density. (a) Genome-wide Manhattan plot showing the location of significant *k*-mers. The horizontal dotted line indicates the significance threshold whereas the vertical dotted line indicates the location of the *Ps* locus. (b) Zoomed-in Manhattan plot at the *Ps* locus. The vertical dotted line indicates the location of the most significant *k*-mer. The shaded gray area shows the location of the causal CNV. (c) Estimated copy number (sequencing depth at position divided by accession average sequencing depth) of accessions with normal and semi-sparse pubescence at the *Ps* locus. Solid lines represent the median copy number across 73 (semi-sparse pubescence) and 180 accessions (normal pubescence) computed using a 100-bp sliding window. Shaded areas outline the first and third quartiles. The start and end of the CNV region can be clearly inferred from the divergence between the copy number of accessions with contrasting pubescence. (d) Histogram showing the location of 1,169 mapped reads containing the most significant *k*-mer in accessions with semi-sparse pubescence. Reads containing this *k*-mer map to either end of the CNV region, suggesting that they are associated with this variant. Panels (b), (c) and (d) share the same x-axis.

The difference in size between the CNV reported by Liu et al. (2020a) and our results may be due to the presence of some unresolved sequence (a stretch of 4,635 N nucleotides) over that interval in the reference sequence that we used. Because our SV filtering process removed any alternate allele containing N nucleotides, we reasoned that the CNV may have been found by our pipeline but filtered out later on. Indeed, we found a 42.8-kb duplication corresponding to the one described above among the genotype calls made by smoove. An association analysis of that single variant using GAPIT found a *p*-value of 1.9e-23, which would have made it the most significant SV. In this particular case, the failure of the SV-based GWAS analysis to find identify the causal variant was due to our filtering parameters and not to limitations of the SV genotype calls per se.

#### Seed coat luster – *B* locus

We conducted a GWAS analysis comparing dull and shiny seed coat phenotypes for the seed coat luster trait. Seed coat luster has been shown by Gijzen et al. (1999) to be largely caused by the deposition of a hydrophobic protein (HPS) at the seed surface. Copy number variation of this gene at the *B* locus was later shown to explain variation in luster, although sequence variation in the HPS sequence may also play a role (Gijzen et al., 2006). Using a sequencing depth-based analysis, we identified a 31-kb region spanning positions 9,386,109-9,417,431 on chromosome Gm15 that exhibited variation in copy number across accessions. This CNV overlapped four genes annotated as containing HPS domains and therefore likely represented the causal variant (Additional file 1: Figures S28 and S29). We found signals overlapping this CNV using all approaches, and the most significantly associated *k*-mer was notably located within the CNV region (Additional file 1: Figures S30 and S28). However, we were not able to directly link any of the variants or *k*-mers identified to the causal CNV.

### Analysis of loci with unknown causal genes

In addition to analyzing loci for which the genes are already known, we were able to suggest candidate genes at loci for which the underlying genes are not known yet.

#### Pod color – *L1* and *L2* loci

Both the *L1* and the *L2* loci for pod color have yet to be cloned. While He et al. (2015) proposed the gene Glyma.19g101700 as likely corresponding to *L1* based on fine mapping and expression analysis, our results as well as those reported by Bandillo et al. (2017) do not support this gene as a candidate due to its location *∼*3 Mb away from GWAS signals. Our analyses based on *k*-mers and SNPs found signals overlapping 14 and 6 genes, respectively (Additional file 1: Figures S31 and S32). Of these, a MATE transporter (Glyma.19g120300) appears as a prime candidate as the most significant *k*-mer was observed within its sequence and it is highly expressed in pods according to data hosted on SoyBase (Grant et al., 2010; Severin et al., 2010). Moreover, the role of some MATE proteins in transporting flavonoids has been demonstrated (Chen et al., 2015).

The signals found by SNP-and *k*-mer-based analyses at the *L2* locus overlap 24 and 50 genes, respectively (Additional file 1: Figures S31 and S33). Despite this somewhat high number, we noticed that the closest gene to the most significant *k*-mer (Glyma.03g005800) is also a MATE transporter, similarly to our top candidate for *L1*. This gene may therefore represent an interesting candidate at this locus for future studies.

#### Pubescence form – *Pa1* and *Pa2* loci

The two known loci for pubescence form in soybean, *Pa1* and *Pa2*, have also not been cloned yet. Nevertheless, Gilbert (2017) has suggested Glyma.12g213900 as a candidate for *Pa1* based on data published by Bandillo et al. (2017). This gene encodes a MYB transcription factor annotated as playing a role in trichome branching. Indeed, the most significant signals found by all three approaches were located near this gene (Figure 5a-b, and Additional file 1: Figures S34 and S35). The analyses based on SNPs and *k*-mers identified two highly significant non-synonymous SNPs in Glyma.12g213900 that were associated with appressed pubescence (Figure 5c-d and Additional file 1: Figures S36 and S37). The alternative alleles for these two SNPs co-occurred together in our dataset and may represent causal variants at the *Pa1* locus. The signals observed at the *Pa2* locus were much weaker and the signal observed from the *k*-mer GWAS overlapped a total of 20 genes (Additional file 1: Figures S34 and S38), none of which appeared obviously linked to pubescence form.

**Figure 5:**
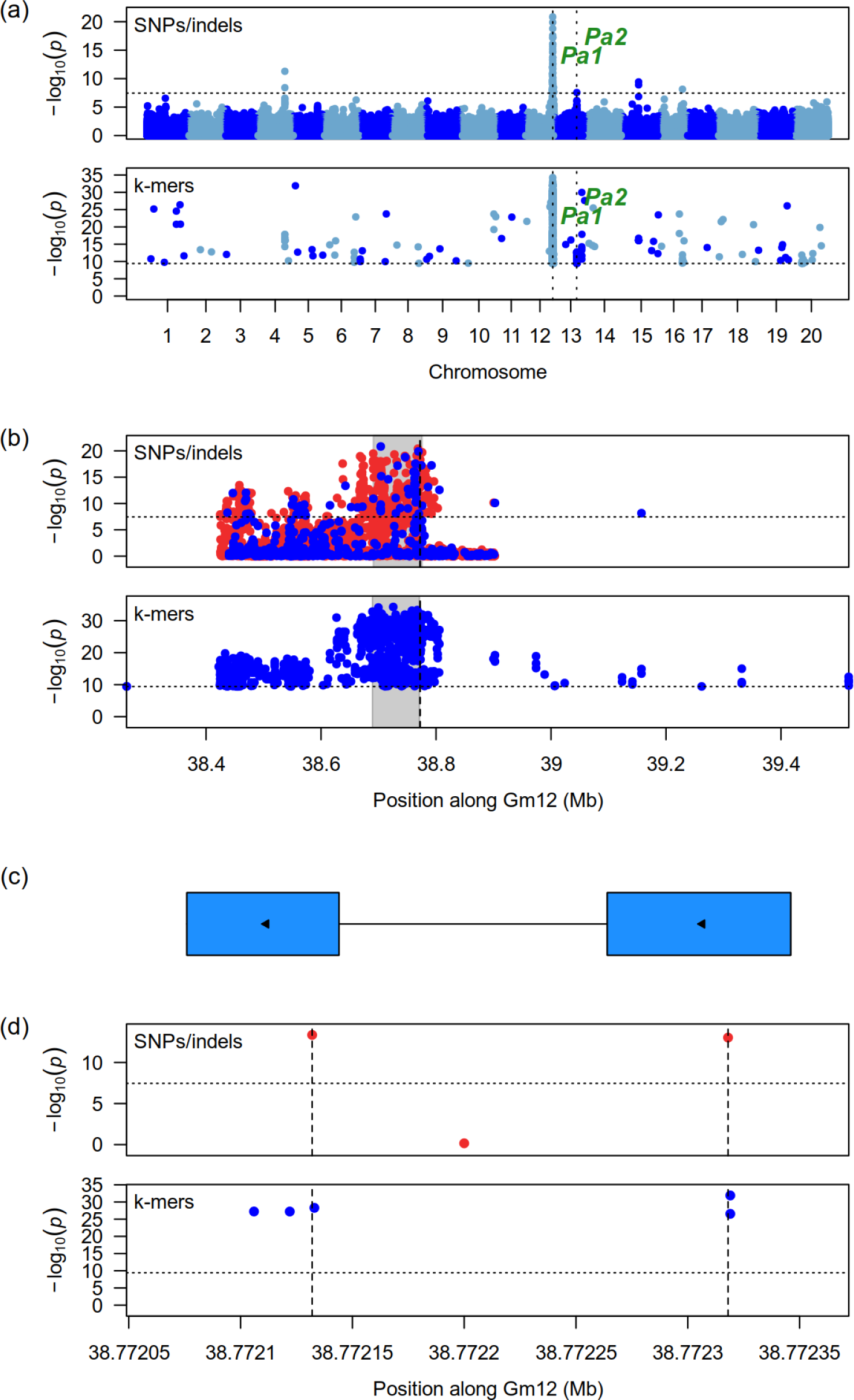
Results of SNP-and *k*-mer-based GWAS at the *Pa1* locus for pubescence form. (a) Genome-wide Manhattan plots. Horizontal dotted lines indicate the significance threshold whereas vertical dotted lines indicate the location of the *Pa1* and *Pa2* loci. (b) Zoomed-in Manhattan plots at the *Pa1* locus. The vertical dotted lines indicate the location of the Glyma.12g213900 gene suggested as a candidate for this locus. The shaded gray rectangles show the regions delimited by the top 1% associations. In the case of SNPs, blue points denote markers used in the original analysis, whereas red points denote markers that had originally been pruned but whose *p*-values were computed after signal discovery. (c) Gene model of Glyma.12g213900. Rectangles represent exons, colored rectangles represent coding sequences, and the arrows indicate the direction of transcription. (d) Manhattan plots at the location of the Glyma.12g213900 gene. Vertical lines indicate the location of two non-synonymous SNPs associated with appressed pubescence, which may represent causal variants at this locus. Panels (c) and (d) share the same x-axis.

A second GWAS analysis focused only on appressed and semi-appressed phenotypes with the objective of enhancing the signal at the *Pa2* locus. However, this analysis only detected weak signals at the *Pa1* locus (Additional file 1: Figure S39) and its results were therefore not analyzed further.

#### Resistance to *P. sojae*

We used corrected dry weight (CDW) as described by de Ronne et al. (2022) as a measure of horizontal resistance to *P. sojae*. We found a strong signal overlapping the genomic region reported by these authors on Gm15 using all three GWAS approaches (Additional file 1: Figure S40). de Ronne et al. (2022) proposed Glyma.15g217100 as a candidate gene for explaining phenotypic variation at that locus based on a combination of functional annotation and contrasting gene expression in resistant and susceptible lines four days after inoculation. Although the signals found by our analyses did overlap Glyma.15g217100, we found that this gene was located at the distal end of the signal in a region with markers or *k*-mers showing a weaker association (Additional file 1: Figure S41). The large size of the observed signal means that the gene associated with the phenotype could be located almost anywhere in this region. However, we identified a particularly interesting region defined by the top 1% associations in the SNP-based analysis. This region overlaps only three annotated genes and also contains the most significant *k*-mer (Additional file 1: Figure S41). One of the genes in this region is a calcium ion binding protein with an EF-hand domain (Glyma.15g217700). Given the important role played by calcium signaling in defense responses (Zhang et al., 2014), we suggest Glyma.15g217700 as an interesting candidate gene for explaining resistance to *P. sojae* in this population.

#### Novel loci for stem termination and pod color

In addition to verifying known loci, we also observed potentially novel loci in the *k*-mer GWAS analysis of stem termination and pod color. When comparing accessions with indeterminate and semi-determinate stem termination types, we observed previously undocumented signals on chromosomes Gm11, Gm16 and Gm18 in addition to the known *Dt1* locus (Figure 6a). Analysis of the pairwise LD between significant *k*-mers found no LD between these signals, suggesting that they may represent *bona fide* loci associated with stem termination type (Figure 6b). Similarly, a *k*-mer GWAS analysis comparing accessions with black and brown pods found a previously undocumented signal on chromosome Gm15 (Additional file 1: Figure S42). *k*-mers at this new locus showed no LD with *k*-mers at the known *L1* locus and may therefore represent a distinct locus (Additional file 1: Figure S43).

**Figure 6:**
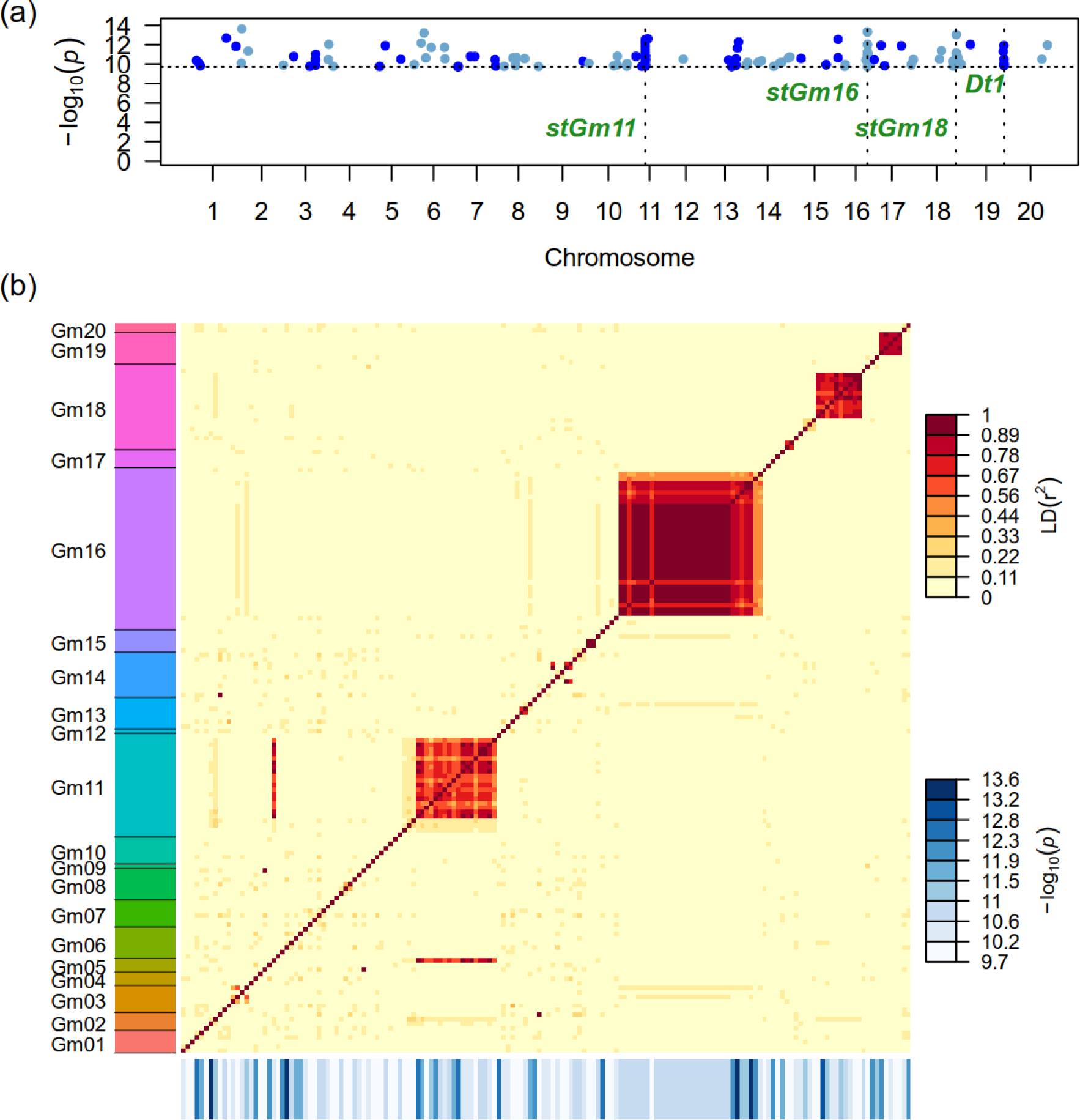
Detection of novel signals for stem termination type using a *k*-mer-based GWAS analysis contrasting accessions with indeterminate and semi-determinate stems. (a) Genome-wide Manhattan plot showing the location of significant *k*-mers. The horizontal dotted line indicates the significance threshold whereas vertical dotted lines indicate the location of the *Dt1* classical locus as well as three new loci of interest (*stGm11*, *stGm16*, and *stGm18*). (b) Pairwise LD between the 162 significant *k*-mers identified. *k*-mers are sorted along the axes according to their putative position on the reference assembly version 4 of Williams82. *k*-mers are represented in the same order along the x-and y-axis. The blue rectangles below the x-axis represent the-log_10_(*p*) of each *k*-mer.

The observed signal overlaps a total of 27 genes (Additional file 1: Figure S44), but none of those are annotated as having an obvious role in controlling pod color.

### Potential of *k*-mer-based GWAS

One of the most striking results from the analyses presented above is the impressive performance of *k*-mer-based GWAS compared to SNP-and SV-based GWAS. The GWAS analyses using *k*-mers identified the causal variant as the most significantly associated variant at four loci with various types of causal variants: *W1* for flower color, *Td* for pubescence color, *G* for seed coat color, and *Ps* for pubescence density (Table 1). In comparison, this result was achieved by SV-based GWAS at the *W1* locus only, and at none of the loci for SNP-based GWAS. The *k*-mer approach also systematically performed at least as well as the other two approaches in identifying signals overlapping known variants or genes (Table 1). Moreover, whenever *k*-mer-based GWAS did detect a signal, the region defined by the 1% most significant *k*-mers overlapped the known causal variant or gene.

Even in cases where the most significantly associated *k*-mers did not pinpoint known causal variants, they were often located near those variants. For example, the most significantly associated *k*-mers mapping to the *T* locus for pubescence color were located in intronic sequences of the gene known to be associated with this locus. Similarly, the most significantly associated *k*-mers at the *B* locus for seed coat luster mapped within a genomic interval known to vary in copy number at that locus. In addition, the most significant *k*-mers at the *R* locus mapped just upstream of a homolog of the gene associated with the locus. Even though the *k*-mer analysis did not identify the causal variants directly in those cases, it still provided a very strong starting point for candidate gene identification.

As would be expected of any GWAS analysis, the success of the *k*-mer approach depends on the quality of the phenotypic data and the complexity of the genetic architecture of the trait. It is clear that the loci for which the *k*-mer approach was most successful (*W1*, *Td*, *G* and *Ps*) have a simple genetic architecture and phenotypes that can be easily determined. For other traits, challenges in determining accurate phenotypes may result in a decreased ability to identify causal variants or genes. For example, Bandillo et al. (2017) noted that observed phenotypes for stem termination type in the GRIN database may not correspond to the expected genotypes at classical loci, and that the intermediate phenotype for seed coat luster is not always consistent across different environments. For qualitative traits, the numerical scale used to recode phenotypes for use in GWAS is somewhat arbitrary and may influence the final results. In the case of quantitative traits, the usual limitations to the precision and stochastic variation of phenotypic values will necessarily make the interpretation more difficult than for simple qualitative traits. Notwithstanding those limitations, *k*-mer-based GWAS clearly performed better than any of the other approaches tested here given the same phenotypic data. This is partly due to the fact that the *k*-mer approach encompasses all variant types. Moreover, it is likely that the simpler workflow that results from using *k*-mer presence/absence avoids some of the errors introduced during variant calling and genotyping, and therefore yields more accurate results.

### Limitations and challenges of *k*-mer-based GWAS

While we have have shown that using *k*-mers for GWAS in soybean performed well compared to other GWAS methods, this approach does present some unique challenges and limitations. One obvious limitation is that using the presence/absence of *k*-mers may not be appropriate in species where heterozygosity is common (i.e., most species other than inbred crops) as the *k*-mers associated with the presence of both alleles will be observed at heterozygous loci. Moreover, although the presence/absence of *k*-mers can detect variation independently of its type, some types of variation may be more difficult to detect than others. For example, copy number variation at a locus may not be detected from presence/absence alone if all copies are identical and occur in tandem, such that no *k*-mer can differentiate between two copies and more. For similar reasons, *k*-mers may also fail to capture variation occurring in repeated sequences. Alternative models based on *k*-mer counts instead of presence/absence have been developed (Rahman et al., 2018; He et al., 2021) which may solve issues like these and broaden the scope of variants that can be identified from *k*-mers.

The complexity of analyzing the results of *k*-mer-based GWAS is another limitation of this approach. Although the method developed by Voichek and Weigel (2020) is in itself relatively simple and computationally inexpensive, downstream analysis and interpretation of the output is challenging. Indeed, there are no state-of-the-art methods for identifying the putative genomic coordinates of a *k*-mer, grouping significant *k*-mers together into coherent signals, and linking them to biologically meaningful sequence variation. Going forward, mapping the *k*-mers or reads to a graph-based genome (e.g. using the vg toolkit; Sirén et al., 2021) may represent an improvement to using a single linear genome. Grouping significant *k*-mers by LD instead of by genomic position, as was done for some of the analyses in this paper, could prove useful in determining how many loci control a trait. In this study, we used an assembly-based approach similar to what others have previously done (e.g. Voichek and Weigel, 2020; Rahman et al., 2018). This approach allowed us to link *k*-mers to their underlying variation for simple cases involving SNPs and indels, but we were unable to link significantly associated *k*-mers to more challenging causal variants such as those explaining the *I* or *B* loci. In this context, developing approaches to systematically link significant *k*-mers to the sequence variation that underlies their presence is much needed. Furthermore, finding ways to limit the number of spurious associations found for some traits, as was the case for flower color and pubescence color in our study, is also needed in order to obtain robust results.

In this work, we provide a set of tools enabling the downstream analysis of *k*-mer-based GWAS. Most importantly, we developed the katcher program for retrieving all reads containing any of a set of significant *k*-mers in a computationally efficient manner. Previous studies usually mapped *k*-mers to a reference genome to identify their reference-based coordinates (e.g. Voichek and Weigel, 2020; He et al., 2021; Rahman et al., 2018; Tripodi et al., 2021). However, while this approach may work for *k*-mers that show little divergence from the reference, it might fail to identify the genomic location of *k*-mers that diverge significantly or are not found at all in the reference. Moreover, it does not take into account the sequence context of the read where the *k*-mer was found, which may provide valuable information for mapping purposes. Our approach avoids these pitfalls by instead systematically retrieving reads containing significant *k*-mers, including those that did not map at all to the reference assembly. While we were able to position the most significant *k*-mers on the reference sequence in our analyses, it may not be the case for highly variable species or if the causal variation at a locus results from a large novel insertion.

### Use of SV genotypes for GWAS

In addition to *k*-mers, we also assessed the use of SVs in GWAS analysis of soybean. Although we did find the causal variant as the most significantly associated SV at the *W1* locus, results were otherwise rather inconclusive. Indeed, most of the other known or putative causal variants were SNPs or indels, and therefore were not represented in the SV datasets. At most loci where SVs are known to be the underlying causal variant (e.g. *I*, *B*, *Ps*), the analyses based on SVs did not find the causal variants, most likely due to the difficulty of calling and genotyping such variants (Kosugi et al., 2019; Lemay et al., 2022). In the case of the *Ps* locus, this limitation was due to a filtering step in our pipeline rather than to an inherent inability to properly call and genotype the causal variant.

Given the current performance of SV genotyping programs, it is not clear whether using SVs for GWAS provides any advantage over using *k*-mer-or SNP-based GWAS, or a combination of both. Indeed, in the single case where the SV-based analyses identified the causal variant, the *k*-mer-based GWAS identified it as well. Moreover, SNP-based GWAS provided better guidance in delimiting the signals found at a given locus because of the much higher density of markers compared to SVs. The usefulness of SV-based GWAS compared to SNP-based GWAS will largely depend on whether SVs can represent variation that is not otherwise in LD with SNPs. In tomato, Domínguez et al. (2020) found that most transposable element insertions associated with phenotypic variation in agronomic traits were not tagged by SNPs. On the other hand, in humans, Maretty et al. (2017) found that the majority of SVs were in high LD with SNPs. It is therefore likely that how well SVs can be tagged by SNPs will depend on the particular type of SV and on the species. A more efficient approach than SV-based GWAS may be to conduct *k*-mer-based GWAS and only then link known SVs to significant *k*-mers. Alternatively, *k*-mer-based GWAS can be conducted specifically for *k*-mers that are associated with SVs (Jayakodi et al., 2020). Previous studies have obtained noteworthy results by using SVs for GWAS analyses (e.g. Zhang et al., 2015; Akakpo et al., 2020; Domínguez et al., 2020), however it would be worthwhile to see whether these results could be replicated using *k*-mer-based GWAS.

## Conclusion

In conclusion, we used SV-based and *k*-mer-based GWAS to study ten qualitative traits and three quantitative traits in soybean and compared the results to conventional SNP-based GWAS. While *k*-mer-based GWAS proved to be a powerful approach in pinpointing the causal variation or genes associated with known loci, it is unclear whether conducting SV-based GWAS is worth the investment. In addition, we used the results from the *k*-mer-and SNP-based GWAS to suggest candidate genes for a few classical loci that have yet to be cloned. Based on our results, we believe that an optimal workflow may involve conducting SNP-based and *k*-mer-based GWAS in parallel to identify significant signals and candidate genes, potentially in combination with SV datasets. Given the large number of samples for which WGS data is now available in several species, applying *k*-mer-based GWAS to leverage already existing sequencing and phenotypic data appears feasible and promising. As part of our work, we developed several computational tools that should help other researchers with the downstream analysis of *k*-mer-based GWAS. However, much work remains to be done in developing state-of-the-art methods for the downstream analysis of significant *k*-mers. In particular, better approaches are needed for linking *k*-mers to sequence variation and moving from presence/absence-based methods to count-based methods.

## Methods

### Sample selection and processing of sequencing data

We selected 389 inbred *G. max* accessions based on the availability of Illumina WGS data and phenotypic data for resistance to the oomycete *P. sojae* (de Ronne et al., 2022). We identified 741 SRA runs corresponding to Illumina paired-end sequencing data for those 389 accessions (data from Zhou et al., 2015; Valliyodan et al., 2016; Fang et al., 2017; Bayer et al., 2022) and extracted paired-end reads using the fastq-dump command v. 2.9.6 (SRA toolkit development team, https://github.com/ncbi/sra-tools) with the option-split-3. Reads that were not paired in the raw sequencing data were not used for downstream analyses. The runs downloaded from the SRA and associated metadata are listed in Additional file 2.

We first filtered the sequencing data for quality and the presence of sequencing adapters using bbduk from BBtools v. 38.25 (Bushnell, https://sourceforge.net/projects/bbmap/). Reads were then mapped to the reference assembly version 4 of *G. max* cultivar Williams82 (Valliyodan et al., 2019) concatenated with chloroplast and mitochondrion sequences obtained from SoyBase (Grant et al., 2010). We used bwa-mem v. 0.7.17-r1188 (Li and Durbin, 2009) with default parameters to map the reads in paired-end mode. Reads that were left unpaired following adapter-and quality-trimming were aligned separately in single-end mode with bwa-mem and merged with mapped paired-end reads using samtools merge v. 1.12 (Li et al., 2009). We then added read groups identifying individual SRA runs to the BAM files and merged the reads belonging to the same accession using bamaddrg (Garrison, https://github.com/ekg/bamaddrg). We used the resulting BAM files for all downstream analyses requiring aligned reads. Metadata on mapping depth of aligned data can be found in Additional file 3.

### Discovery and genotyping of SVs

Our SV analysis pipeline consisted of separate SV discovery and genotyping steps. The *discovery* step identified a set of candidate SVs using various methods, whereas the *genotyping* step determined the genotype of the candidate SVs for all accessions in the population from the mapped Illumina reads. We used three different approaches in the SV discovery step to generate a set of candidate variants:

- We used Illumina WGS data from the 389 accessions mentioned above to call SVs in the population following a pipeline outlined by Lemay et al. (2022). In brief, we combined information from the SV calling programs AsmVar (Liu et al., 2015), Manta (Chen et al., 2016), smoove (Pedersen and Quinlan, 2019) and SvABA (Wala et al., 2018) to generate a set of candidate SVs from the mapped Illumina data.
- We used SVs discovered by Lemay et al. (2022) from Oxford Nanopore sequencing data of 17 Canadian soybean cultivars.
- We called SVs from 26 high-quality genome assemblies published by Liu et al. (2020b), as well as that of the cultivar Zhonghuang 13 (ZH13), *G. soja* accession W05, and *G. max* cultivar Lee. We used methods similar to those of Liu et al. (2020b) to call SVs from the comparison of these assemblies to that of Williams82.

The SVs called using each of these approaches were merged using SVmerge (Wong et al., 2010) and the resulting set of candidate SVs was used for genotyping. We used Paragraph (Chen et al., 2019) to genotype the candidate SVs from the mapped Illumina reads for all accessions. We filtered the genotyped SVs by setting the genotype calls made from less than two reads to missing and by removing variants with a minor allele frequency < 0.02 or a proportion of missing data > 0.5. The resulting dataset of 186,306 genotyped SVs was used for GWAS. Detailed methods regarding the SV discovery and genotyping steps can be found in Additional file 1.

### Discovery and genotyping of SNPs and indels

We used Platypus v. 0.8.1.1 (Rimmer et al., 2014) to call SNPs and indels from the mapped reads of the 389 accessions. The 21.1 M SNPs and indels from the raw output of Platypus were filtered to keep only those with:

1. FILTER field set to PASS
2. minor allele frequency (MAF) *≥* 0.02
3. fraction of missing genotypes *≤* 0.5
4. heterozygosity rate *≤* 0.2

Moreover, we pruned the dataset using the-indep-pairwise option of PLINK v. 1.90b5.3 (Purcell et al., 2007) with a window of 1,000 markers, a step of 100 markers and an *r*^2^ LD threshold of 0.9. This pruning step was implemented to reduce the computational requirements of the GWAS analysis by removing co-segregating markers. Markers located on unanchored scaffolds were also removed. The resulting dataset was used for GWAS and comprised 773,060 SNPs and 151,570 indels.

### Computing the presence/absence table of *k*-mers

We used the approach outlined by Voichek and Weigel (2020) for *k*-mer-based GWAS and their recommendations on the associated GitHub page (v. 0.2-beta, https://github.com/voichek/kmersGWAS) to generate a presence/absence *k*-mer table for use in GWAS. The first step of the approach involved counting all *k*-mers of length 31 (31-mers) present in the trimmed (FASTQ) reads for each accession using KMC3 v. 3.2.1 (Kokot et al., 2017). *k*-mers were counted twice, the first time using their canonized form (the first of either the observed *k*-mer or its reverse complement in lexicographical order) and the second time using the observed *k*-mers themselves (non-canonized form). The two sets of *k*-mers were then combined and filtered by keeping only those seen in at least 5 accessions, and observed in both canonized and non-canonized form in at least 20% of the accessions in which they are found. We then generated a table indicating the presence or absence of each *k*-mer in all accessions. This table was used to compute a kinship table for use in GWAS and was also used directly as input genotypes for the GWAS analyses. We used *k*-mers with a MAF *≥* 0.02 for both the computation of kinship and GWAS.

### Phenotypic data

We analyzed ten qualitative traits previously studied by Bandillo et al. (2017): flower color, pubescence color, seed coat color, stem termination type, hilum color, pod color, pubescence form, pubescence density, seed coat luster, and maturity group. We obtained phenotypic data for these traits from the GRIN database (https://npgsweb.ars-grin.gov/gringlobal/search) by querying the database using the PI identifiers of the 389 accessions used. Four out of the 389 accessions did not match the GRIN database and were therefore not used for the analysis of qualitative traits. The phenotypes of qualitative traits were recoded to numerical values for GWAS analysis following methods similar to those of Bandillo et al. (2017). For some traits, we conducted more than one GWAS analysis using targeted subsets of the observed phenotypes in order to focus on specific loci. The phenotypes and numerical values used for all analyses as well as the number of observations in the dataset that we used for GWAS are listed in Tables S2 to S11 (Additional file 1).

We analyzed three quantitative traits in addition to the ten qualitative traits mentioned above. Seed oil and protein content were retrieved from the GRIN database. In cases where more than one value was listed for a given accession, we computed the average of those values and used it for GWAS analyses. As a third quantitative trait, we used the horizontal resistance to the oomycete *P. sojae* (corrected dry weight, CDW) as recently assessed in a hydroponic assay (de Ronne et al., 2022). Our analysis included 340 accessions whose phenotypic data was already published by de Ronne et al. (2022) as well as 49 additional accessions for which sequencing data was not yet available when their study was conducted. Because of ambiguity regarding the identity of one of the accessions (HN019), this accession was dropped from all GWAS analyses. Similarly to seed oil and protein content, the CDW values for some accessions for which more than one observation was available were averaged.

Because of clear discrepancies between the observed haplotypes and the reported phenotypes for some simple traits such as flower color and seed coat color, we suspected possible errors in the identity of some of the accessions. To investigate this, we compared the SNP genotypes derived from the SoySNP50K array and our WGS data at over 32,000 SNPs, and identified 24 accessions with < 90% concordance between the two genotype datasets (see Additional file 1 for detailed methods). These samples were removed from the dataset in addition to another accession that exhibited an atypical GC content suggesting contamination. Following these filtering steps and the removal of HN019, 363 accessions remained for GWAS analyses. The concordance between WGS data and SoySNP50K genotypes is included in Additional file 3 and averaged 98.3% among retained lines (Additional file 1: Figure S45). The phenotypic data used in this study can be found in Additional file 4.

### GWAS analyses

We used the GAPIT3 R package v. 3.1.0 (Wang and Zhang, 2021) for conducting GWAS on the SNP/indel and SV datasets described above. SVs and indels required some additional preprocessing steps to make variant representation suitable for downstream analyses (Lemay and Malle, 2022). Briefly, we recoded all variants as SNPs prior to converting the VCF files to Hapmap diploid format using the TASSEL command-line tools (Bradbury et al., 2007); variant IDs enabled retrieval of the proper metadata for each variant after GWAS.

We ran GAPIT using a mixed linear model (MLM) with 9 principal components and the VanRaden algorithm for computing the kinship matrix. We used an MLM model for consistency with the statistical model used by the *k*-mer-based approach. We used a randomization approach to determine the 5% family-wise error-rate threshold for each GWAS analysis as described in Voichek and Weigel (2020). To compute this threshold, we permuted the phenotypic observations 100 times and computed a GWAS on these permuted phenotypes to obtain a distribution of top *p*-values under the null hypothesis. The fifth most significant of the top 100 *p*-values was used as the significance threshold for inference, i.e. markers with *p*-values lower than this threshold were considered significant.

We conducted GWAS based on *k*-mers using the method developed by Voichek and Weigel (2020). In brief, the analysis was conducted in two steps. The first step used an approximate model to identify 1 million potentially significant *k*-mers. These were then used in the second step as input to an exact model implemented in GEMMA (Zhou and Stephens, 2012), resulting in a *p*-value for each *k*-mer. We used the kinship computed from the *k*-mers to correct for relatedness between accessions. The program automatically outputs a list of *k*-mers that passed a 5% family-wise error-rate threshold determined by a randomization approach, which we considered significant for the purposes of downstream analyses.

### Analysis of significant *k*-mers

*k*-mers are not intrinsically associated with a particular genomic region and a tailored analysis was therefore required to associate them with genomic coordinates. To do so, we queried the mapped reads (in BAM format) of all accessions for matches to significant *k*-mers for a given trait. Because this is a computationally demanding operation (there were typically tens of thousands of significant *k*-mers per trait), we developed a C program called katcher (https://github.com/malemay/katcher) that uses the htslib library (Bonfield et al., 2021) for efficient reading and writing of BAM files. katcher and its associated utilities allow for efficient retrieval and annotation of mapped reads containing any *k*-mer from a set of interest.

Once we retrieved all the reads containing significant *k*-mers, we next needed to gather the mapping information contained in those reads for the purposes of identifying contiguous signals and comparing them to annotations. To simplify the amount of information to process in downstream analyses, we considered only the most significant *k*-mer contained in a given read and the position where this *k*-mer was most often observed in the dataset. Moreover, we limited the analysis to reads with a minimum mapping quality of 20 and to *k*-mers that were observed at least 10 times in at least one accession. The latter was done to remove spurious *k*-mers that may originate from sequencing errors. All downstream analyses (Manhattan plots, identification of signals, analysis of genes located near significant *k*-mers, etc.) used this processed set of significant *k*-mers.

### Downstream analyses of significant association signals

We used contiguous regions of significant variants, which we called “signals”, as a basis to compare our results with those obtained by previous studies and to query the genome for candidate genes. We defined the boundaries of those signals by grouping any significant markers or *k*-mers that occurred within 250 kb of one another. We did consider the length of deletions or *k*-mers in defining the distance between significant variants.

For the GWAS analyses computed from SNP/indel genotype calls, we retrieved all markers that had initially been pruned in a region ranging from 50 kb downstream to 50 kb upstream of significant signals and computed their *p*-values by running GAPIT using the same parameters as noted above. The *p*-values thus computed on markers that were initially left out were not used for initial signal discovery, but provided finer resolution when analyzing the results of the SNP-based GWAS analyses. Indeed, some markers that had not been included in the original GWAS may be the causal variant or simply yield more significant *p*-values than those initially used.

To help with the interpretation of results and the identification of candidate genes, we defined regions of highly significant associations within each signal by delimiting a range of coordinates containing the top 5% most significant associations (Paragraph) or 1% most significant associations (SNPs/indels and *k*-mers) found within the signal. In cases where such a fraction represented fewer than two markers or *k*-mers, the highly significant region was simply delimited by the two most significant associations.

Because previous studies (Bandillo et al., 2015, 2017) used assembly version 1 of Williams82, we converted the genomic coordinates of their signals to assembly version 4 using the same method that we used for converting SoySNP50K positions (see Additional file 1). For the analysis of qualitative traits, we only compared our results to signals at named classical loci and ignored previously reported minor signals with *p*-values that just barely made it above the significance threshold.

### Linking significant *k*-mers to variants

We used the reads retrieved by katcher and an assembly-based approach for identifying the variants associated with a given set of *k*-mers. In addition to the reads themselves, we also retrieved their paired reads even though they may not have matched a significant *k*-mer in the first place. These read pairs were used for *de novo* assembly with SPAdes v. 3.15.4 (Prjibelski et al., 2020) using the –careful parameter. The assemblies were done individually for each accession and then aligned to the Williams82 reference assembly using bwa-mem with default parameters. Aligned assembled sequences that overlapped loci of interest were realigned using multiple sequence alignment with the MAFFT program v. 7.475 (Katoh et al., 2002) for identifying and visualizing the haplotypes at these loci. The identification of variants supported by significant *k*-mers was based entirely on visual analysis of haplotype alignments. We only considered haplotypes occurring at least five times in the population in such analyses in order to leave out potential assembly errors. In a single case (*Dt1* locus), assembly at the locus succeeded for a very low number of accessions (26). In this case, we instead used consensus sequences from the reads aligned with bwa for comparing the haplotypes across accessions.

### Computation of LD between *k*-mers

In some cases, the analysis of *k*-mers revealed signals at previously undocumented loci or found a very large number of associations throughout the genome. To shed light on these situations, we computed the pairwise LD between *k*-mers in order to identify *k*-mers that co-segregated and likely corresponded to a single locus. Because of the sheer number of pairwise comparisons that needed to be made for some traits (> 20,000 significant *k*-mers kept even after the filtering steps mentioned above), we limited the number of *k*-mers used for LD calculation to 1,500. For traits where more than 1,500 *k*-mers were found, we computed LD on a subsample of these by selecting the 500 *k*-mers with the most significant *p*-values and sampling the remaining 1000 *k*-mers randomly. This random sampling was done with a probability inversely proportional to the number of *k*-mers matching a given chromosome or scaffold such that *k*-mers were sampled from the whole genome and not simply from the most significant locations.

### Software used

Unless otherwise noted, all analyses were conducted using R v. 4.2.0 (R Core Team, 2022) and Bioconductor v. 3.15 (Huber et al., 2015). We used Bioconductor packages Biostrings v. 2.64.0 (Pagès et al., 2022), GenomicRanges v. 1.48.0 (Lawrence et al., 2013), Rsamtools v. 2.12.0 (Morgan et al., 2022), rtracklayer v. 1.56.0 (Lawrence et al., 2009) and VariantAnnotation v. 1.42.1 (Obenchain et al., 2014). We gathered several functions used for the downstream analysis of GWAS results into a package called gwask which is available on GitHub (https://github.com/malemay/gwask).

## Additional files

**Additional file 1.pdf** Supplemental methods and results, including Tables S1 to S11 and Figures S1 to S60 (PDF 12.9 MB)

**Additional file 2.csv** Metadata on the sequencing runs retrieved from the Sequence Read Archive (SRA) (CSV 71 KB)

**Additional file 3.csv** Metadata on individual accessions following mapping and comparison to SoySNP50K data (CSV 36KB)

**Additional file 4.csv** Phenotypic dataset used for GWAS analyses (CSV 135 KB)

## Availability of data and materials

Some of the datasets generated as part of this work are available on figshare (Lemay et al., 2023b).

The Illumina sequencing data used for the analyses are listed in Additional file 2 and available through NCBI BioProjects PRJNA257011, PRJNA289660 and PRJNA639876.

The high-quality assemblies generated by Liu et al. (2020b) are available on the Genome Warehouse through Accession Number PRJCA002030.

The assemblies of ZH13, W05 and Lee are available on SoyBase (https://soybase.org/GlycineBlastPages/blast_descriptions.php).

The *I* locus contig assembled by Tuteja and Vodkin (2008) is available from NCBI (https://www.ncbi.nlm.nih.gov/nuccore/EF623854).

SoySNP50K genotype calls are available from SoyBase (https://soybase.org/snps/).

The SVs identified from Oxford Nanopore data by Lemay et al. (2022) are available on figshare (Lemay et al., 2021).

The following software used in this work is available on GitHub:

- Code used for the analyses: https://github.com/malemay/soybean_kmer_gwas
- katcher software for retrieving reads containing *k*-mers: https://github.com/malemay/ katcher
- gwask R package used for processing the output of GAPIT and *k*-mer GWAS analyses, and for plotting: https://github.com/malemay/gwask
- Software for computing the LD between *k*-mers based on presence/absence: https://github.com/malemay/kmers_ld
- Forked svmu version that was used for the analyses shown in this paper: https://github.com/malemay/svmu
- svmutools R package for converting the output of svmu to VCF format: https://github.com/malemay/svmutools

The code associated with the results presented in this manuscript is archived on figshare (Lemay et al., 2023a).

## Authors’ contributions

Conception and design of the study: MAL, MDR, RB, FB. Phenotypic data acquisition (resistance to *Phytophthora sojae*): MDR, RB. Data analysis: MAL. Data interpretation: MAL, FB. Software writing: MAL. Manuscript drafting: MAL, FB. All authors have revised the manuscript and approved its submission.

## Acknowledgements

We would like to thank Jonas A. Sibbesen for providing very helpful comments on the methodology used and on a previous version of this manuscript. We also thank Yoav Voichek for help regarding *k*-mer GWAS analysis as well as Brian Boyle and Martine Jean for valuable comments regarding the methodology. We are thankful to the Digital Research Alliance of Canada for using their high-performance computing servers.

