## Additional file 1 for "*k*-mer-based GWAS enhances the discovery of causal variants and candidate genes in soybean"

### Detailed methods

#### Discovery of structural variation from Illumina data

We used the Illumina whole-genome sequencing (WGS) data from the 389 selected *G. max* accessions for SV discovery using methods similar to those described by Lemay et al. (2022). Briefly, we used four different SV discovery programs or pipelines:

1. We used AsmVar (version of 2015-04-16, Liu et al., 2015) to call SVs from *de novo* assemblies generated with SOAPdenovo2 v. 2.04 (Luo et al., 2012) and aligned to the reference genome using LAST v. 1047 (Kielbasa et al., 2011).
2. We ran Manta v. 1.6.0 (Chen et al., 2016) in randomly selected batches: 77 batches of 5 samples and 1 batch of 4 samples. We used the candidate SVs identified from each of the batches for downstream analyses.
3. We used smoove v. 0.2.4 (Pedersen and Quinlan, 2019) to obtain a dataset of candidate SVs using the following sequence of commands: `smoove call`, `smoove merge`, `smoove genotype` and `smoove paste`. For three (3) of the 389 samples, we were unable to obtain candidate variants due to a reproducible error (segmentation fault) when running

`smoove call`.

4. We used SvABA v. 1.1.3 (Wala et al., 2018) to call SVs from each of the samples separately. We classified the SVs called as breakends by SvABA as either deletions, duplications or inversions, and converted them to explicit sequence using a custom R script. Variants called as indels by SvABA were used as is for downstream analyses.

We filtered the SVs called by each tool to remove those meeting any of the following conditions:

- smaller than 50 bp or larger than 500 kb in size
- located on unanchored scaffolds or organellar genomes
- classified as unresolved breakends (`SVTYPE=BND`)
- containing at least one N in the alternate allele sequence

Next, we converted all filtered variants to a sequence-explicit format using `bayesTyperTools convertAllele` (Sibbesen et al., 2018) and normalized them using bcftools `norm` v. 1.10.2-105 (Li et al., 2009).

A single VCF file was generated from the variants called using Illumina data by running SVmerge (Wong et al., 2010) in two steps. In the first step, we merged very similar variants among the ones called by the same tool using a sliding window of 5 bp. Next, we used the output of the first merging steps for all four tools and merged similar variants across the four datasets using a sliding window of 15 bp. This approach was used because a sliding window of 15 bp on the outputs of all tools without pre-merging resulted in a computationally intractable number of pairwise comparisons.

### Discovery of structural variation from Oxford Nanopore sequencing data

We used SVs discovered among 17 Canadian soybean cultivars sequenced using Oxford Nanopore Technologies (ONT) by Lemay et al. (2022) as an additional source of candidate SVs. Although this set of cultivars does not overlap the ones analyzed in this study, it is likely that many of the SVs occurring in Canadian germplasm also occur in the germplasm used for this study given that modern soybean cultivars are derived from a rather small set of progenitors. We merged the SVs detected in the long-read (ONT) data with the ones discovered in the short-read (Illumina) data using SVmerge and systematically favored SVs discovered by Illumina whenever SVs from the two sets were to be merged. Our rationale for favoring variants discovered from the Illumina data was that whenever both technologies discover the same variants, the breakpoints of SVs discovered from Illumina data are likely to be more precise given the higher basecalling accuracy.

### Discovery of structural variation from high-quality assemblies

In addition to Illumina data from 389 accessions, we also used high-quality genome assemblies of 29 *G. max* or *Glycine soja* accessions to call SVs. We retrieved 26 high-quality genomes assembled by Liu et al. (2020) as well as the genome of the cultivar Zhonghuang 13 (ZH13) from the Genome Warehouse repository (`https://ngdc.cncb.ac.cn/gwh/`). Furthermore, we retrieved the genomes of *G. soja* accession W05 and *G. max* cultivar Lee from SoyBase (Grant et al., 2010).

We used methods based on Liu et al. (2020) to call SVs from the genome assemblies. First, we used the nucmer program of the MUMmer suite of tools v. 4.0.0rc1 (Marçais et al., 2018) with option `-c 1000` to align each of the assemblies to the Williams82 assembly. Next, we used the delta-filter program of the MUMmer suite to keep only one-to-one alignments between any genome and the Williams82 reference using the `-1` option. The filtered alignments were used to call SVs with a version of the svmu program (Chakraborty et al., 2019) that we forked from the original repository (`https://github.com/malemay/svmu`) in order to improve execution time and memory usage.

We used a set of tools that we assembled in an R package called svmutools (`https://github.com/malemay/svmutools`) to convert the svmu output to VCF format. We limited ourselves to processing deletions and insertions because of the complexity of resolving other types of variants from whole-genome assemblies. Copy number variants and inversions called by svmu were therefore excluded from downstream analyses. The resulting set of variants was filtered to remove those smaller than 50 bp or larger than 500 kb and those with any N in either the reference or alternate allele sequence. Finally, we normalized the VCF files using `bcftools norm` and merged them using SVmerge to obtain a single file with variants discovered through comparison of high-quality genome assemblies.

### Genotyping structural variation using Paragraph

We used Paragraph v. 2.4a (Chen et al., 2019) to generate a set of SV genotypes from the Illumina data of the 389 accessions. The set of SVs used as input for Paragraph was generated using SVmerge by merging variants discovered from the Illumina and ONT data with those discovered from the high-quality assemblies and svmu. Variants in this candidate set were prepared for input to Paragraph by first removing those closer than 150 bp (the maximum read length in our dataset) to any chromosome end and then padding the variant representation using a script written by Hickey et al. (2020).

We ran Paragraph individually on each sample as recommended by the authors. We computed the average sequencing depth for each sample from the output of the `samtools coverage` command and used 20 times that value for the `-M` parameter of Paragraph. We ran Paragraph using the `multigrmpy` command and merged the resulting VCF files using `bcftools merge` in order to get a single genotype file for the whole population. The resulting genotype calls were filtered as described in the main text.

### Comparison of WGS and SoySNP50K genotypes

When we first analyzed the results from the GWAS analysis on all 389 samples, we found discrepancies between the haplotypes at some loci and the expected phenotype that should correspond to that haplotype for simple traits such as flower color and seed coat color. Upon further analysis, we found that a mismatch between the sequencing data and the identity of the accession was the most probable cause of these discrepancies. In order to identify the samples for which this could be the case, we compared the genotypes of SNPs obtained from the SoySNP50K array (Song et al., 2013) to those predicted from the Illumina WGS data. We did this for the 385 samples that had a match in the GRIN database. Briefly, we retrieved SoySNP50K genotype calls from SoyBase (Grant et al., 2010) and extracted all SNPs with a MAF $\geq 0.1$ in our population. We then translated the positions of these SNPs from assembly version 2 to assembly version 4 of Williams by finding exact matches of 41-bp sequences surrounding the SNP positions. The 32,852 SNPs for which unambiguous positions could be found on genome assembly version 4 were genotyped from the mapped WGS reads using `bcftools mpileup` and `bcftools call`. We then used `bcftools gtcheck` to compute the concordance between the WGS genotype calls and the SoySNP50K genotype dataset. We identified 24 samples with $< 90\%$ concordance between the two genotype datasets and removed them from the dataset. We also excluded an additional sample for which concordance was over 90% but poor raw sequencing data quality (%GC content) suggested contamination. The concordance between WGS data and SoySNP50K genotypes is included in Additional file 3 and shown in Figure S45.

#### Analysis of extremely low *p*-values

For one trait (flower color), the significance of some $k$-mers was so high that their $p$-values were numerically equal to 0. In order to compute $-\log_{10}(p)$ for visualization in Manhattan plots in these cases, we set the $p$-value to the smallest numerical value that could be represented using R on our machine (2.225074e-308). In this particular case, the values represented on the Manhattan plots represent an upper bound on the $p$-value and not an exact value.

### Supplemental results

This section describes results at additional loci that were not discussed in the main text. A summary of the signals found at all loci using all three GWAS methods is presented in Table S1.

#### Flower color – *L1* locus

In addition to the *W1* locus, Bandillo et al. (2017) reported a signal for flower color near the *L1* locus. None of the GWAS approaches that we tested found a signal for flower color near this locus (Figure S1). This result is unsurprising as W1 is the major contributor to flower color in soybean, whereas the *L1* locus (typically associated with pod color) was detected with a much less significant $p$-value (see Table S1) and only conjectured as associated with flower color by Bandillo et al. (2017).

#### Pubescence color – locus on chromosome Gm16

In addition to the *T* locus, the first GWAS analysis conducted on pubescence color revealed a second locus on chromosome Gm16 (Figure S5). Upon closer analysis, this locus appeared to be in strong LD with $k$-mers mapping to chromosome Gm01 and most importantly in moderate LD with the $T$ locus (Figure S46), which suggests that this does not represent an independent locus controlling pubescence color. This interpretation is also supported by the fact that the signal at that locus only spans about 10 kb on chromosome Gm16.

### Seed coat color – loci *T*, *O* and *R*

In addition to the $I$ and $G$ loci, the $k$-mer-based GWAS analysis of five different seed coat color phenotypes (first GWAS in Table S4) detected weak signals overlapping the $T$ and $O$ loci (Figure S12c). However, we interpreted those results as chance associations given the low $p$-value associated with the $T$ locus and the proximity of the $O$ locus to the $I$ locus. None of the three GWAS approaches detected a signal overlapping the $R$ locus.

### Stem termination type – loci *Dt2* and *E3*

We performed a GWAS analysis comparing only semi-determinate and indeterminate stem termination phenotypes in an attempt to detect signals at the *Dt2* locus (Table S5). However, this was unsuccessful using all three GWAS approaches (Figure S47). Similarly, we failed to detect signals at the *E3* locus, which is more commonly associated with maturity but was mentioned as associated with stem termination type by Bandillo et al. (2017).

### Hilum color – *T* locus

The $T$ locus discussed in the main text for pubescence color is also a determining locus for hilum color. We detected signals overlapping the corresponding gene using all approaches (Figures S48, S49). The regions defined by the top 5% (SVs) and top 1% (SNPs and $k$-mers) variants included the Glyma.06g202300 gene associated with this locus. Similarly to what we observed for the analysis of the $T$ locus based on pubescence color, we did identify the documented causal variant at this locus using the Platypus and $k$-mer analyses, although these variants were not the most significantly associated overall (Figures S50, S51).

### Hilum color – *I* locus

Similarly, the *I* locus discussed in the main text for its effect on seed coat color also controls hilum color and we detected signals overlapping the associated inverted duplication using all approaches (Figures S48, S52). However, as was the case for the effect of *I* on seed coat color, we were not able to link any of the variants identified to the causal variation given its highly complex nature.

### Hilum color – loci *W1* and *O*

None of the three GWAS analyses that we conducted on hilum color found signals at the *W1* and *O* loci studied by Bandillo et al. (2017) (Figures S21 S48, S53).

### Pubescence density – loci *Pd1* and *P1*

Our GWAS analysis on pubescence density did not detect any signals at the *Pd1* and *P1* (Figure S25). Our failure to detect these loci is likely due to our small population size compared to Bandillo et al. (2017) and to the smaller contribution of those loci to pubescence density in our population.

### Seed coat luster – loci *Hps*, *B?* and *I*

We performed three different GWAS analyses on seed coat luster (Table S10). Since all three analyses yielded similar results (Figure S54, S55 and S30), we chose to focus on the GWAS contrasting the dull and shiny phenotypes as it yielded the most significant signals. In addition, it also provided results that were most consistent with the analysis previously done by Bandillo et al. (2017).

The analysis detected signals at the *B* locus as the sole classical locus using all three methods (see main text and Figure S30). All methods also detected signals overlapping the *Hps* locus as defined by Bandillo et al. (2017), however results shown by Gijzen et al. (2003) and our results suggest that *Hps* and *B* are the same locus.

Apparent signals on chromosomes Gm20 and Gm09 (which may correspond to the B? locus reported by Bandillo et al. (2017)) were also detected, but analysis of the LD patterns between $k$-mers suggested that these signals are most likely in LD with the *B* locus (Figure S56).

Bandillo et al. (2017) mentioned the *I* locus as possibly associated with seed coat luster, however we did not find evidence for such an association in our dataset.

### Pubescence form – additional loci

The first GWAS analysis on pubescence form found signals on chromosomes Gm04 and Gm15 with SNPs and $k$-mers (Figure S34). However, upon further analysis of LD patterns, they appeared to be linked to the *Pa1* locus (Figure S57).

### Maturity group

We did not detect any signals associated with maturity using our dataset (Figure S58). The use of maturity group as a proxy for time to maturity instead of the number of days, combined with the limited phenotypic variation of this trait in this collection of accessions (Table S11), likely explains our failure to detect any signals.

### Seed oil and protein content

We conducted a GWAS analysis on seed oil content using data obtained from the GRIN database. While several seed oil content quantitative trait loci (QTL) have been reported due to high interest in this trait for soybean breeding (Chaudhary et al., 2015), we focused on two major QTL on chromosomes Gm15 and Gm20. These loci have often been reported in *G. max* and were notably observed by Bandillo et al. (2015) on a collection of USDA lines using SoySNP50K chip genotypes and GRIN phenotypic data. We identified a signal located on chromosome Gm15 using the $k$-mer approach only (Figure S59). Although this signal did not overlap with the coordinates found by Bandillo et al. (2017), it was located only 2 kb away from it and 30 kb away from Glyma.15g049200, the sugar transporter gene identified by Zhang et al. (2020) as associated with that locus. Therefore, the significantly associated $k$-mers were not linked to the causal variation at that locus.

Similarly to seed oil content, we conducted a GWAS on seed protein content using data obtained from the GRIN database. As is generally the case in soybean given the negative correlation between seed oil and protein content, the two major QTL considered above and reported by Bandillo et al. (2015) for seed oil content are also major QTL for seed protein content. However, none of the approaches detected a signal at this locus (Figure S60).

Our failure to detect signals overlapping the known genes at these two loci for oil and protein is probably due to the fact that the oil and protein content data that we used for GWAS were not obtained from orthogonal trials, which was a clear limitation to this GWAS given our relatively small sample size.

### Supplemental tables

**Table S1:** Description of loci associated with the traits studied and -$\log_{10}(p)$ of significant signals detected in our study using three different genotype datasets (SNPs/indels, SVs, $k$-mers).

| Trait | Locus | Chromosome | -$\log_{10}(p)$[a] | Gene[b] | SNPs/indels[c] | SVs | $k$-mers | Study |
|---|---|---|---|---|---|---|---|---|
| Flower color | *W1* | Gm13 | 169.8 | Glyma.13g072100 | 42.2 | 51.6 | 307.7[d] | Bandillo et al. (2017) |
| Flower color | *L1* | Gm19 | 14.8 | - | - | - | - | Bandillo et al. (2017) |
| Pubescence color | *T* | Gm06 | 298.2 | Glyma.06g202300 | 29.1 | 26.6 | 59.5 | Bandillo et al. (2017) |
| Pubescence color | *Td* | Gm03 | 95.8 | Glyma.03g258700 | 11 | 9.8 | 24.8 | Bandillo et al. (2017) |
| Seed coat color | *G* | Gm01 | 274.1 | Glyma.01g198500 | 29 | 16.1 | 97 | Bandillo et al. (2017) |
| Seed coat color | *I* | Gm08 | 37.5 | CHS gene cluster | 24.8 | 25.7 | 56.8 | Bandillo et al. (2017) |
| Seed coat color | *T* | Gm06 | 11.1 | Glyma.06g202300 | - | - | 14.4 | Bandillo et al. (2017) |
| Seed coat color | *O* | Gm08 | 8.6 | - | - | - | 11.1 | Bandillo et al. (2017) |
| Seed coat color | *R* | Gm09 | 5.8 | Glyma.09g235100 | - | - | - | Bandillo et al. (2017) |
| Stem termination type | *Dt1* | Gm19 | 238.8 | Glyma.19g194300 | 19.3 | 20.1 | 29.7 | Bandillo et al. (2017) |
| Stem termination type | *Dt2* | Gm18 | 17.3 | Glyma.18g273600 | - | - | - | Bandillo et al. (2017) |
| Stem termination type | *stGm11* | Gm11 | - | - | - | - | 12.5 | This study |
| Stem termination type | *stGm16* | Gm16 | - | - | - | - | 13.3 | This study |
| Stem termination type | *stGm18* | Gm18 | - | - | - | - | 13 | This study |
| Stem termination type | *E3* | Gm19 | 6.2 | Glyma.19g224200 | - | - | - | Bandillo et al. (2017) |
| Hilum color | *T* | Gm06 | 100.3 | Glyma.06g202300 | 13.1 | 10.5 | 17.6 | Bandillo et al. (2017) |
| Hilum color | *I* | Gm08 | 96.6 | CHS gene cluster | 17.5 | 17.1 | 26.7 | Bandillo et al. (2017) |
| Hilum color | *R* | Gm09 | 80.6 | Glyma.09g235100 | 12.2 | 11.8 | 26.6 | Bandillo et al. (2017) |
| Hilum color | *O* | Gm08 | 35 | - | - | - | - | Bandillo et al. (2017) |
| Hilum color | *W1* | Gm13 | 8.6 | Glyma.13g072100 | - | - | - | Bandillo et al. (2017) |
| Pod color | *L2* | Gm03 | 147.9 | - | 10.2 | 8.6 | 14.6 | Bandillo et al. (2017) |
| Pod color | *L1* | Gm19 | 127.5 | - | 11.9 | 11 | 17.2 | Bandillo et al. (2017) |
| Pod color | *pdcGm15* | Gm15 | - | - | - | - | 13.5 | This study |
| Pubescence form | *Pa1* | Gm12 | 251.3 | Glyma.12g213900? | 20.8 | 19.9 | 34.3 | Bandillo et al. (2017) |
| Pubescence form | *Pa2* | Gm13 | 26.9 | - | 7.6 | - | 12.8 | Bandillo et al. (2017) |

Continued on next page

**Table S1:** Loci associated with the traits studied *(continued)*

| Trait | Locus | Chromosome | -log10(p) | Gene | SNPs/indels | SVs | k-mers | Study |
|---|---|---|---|---|---|---|---|---|
| Pubescence density | *Ps* | Gm12 | 224.3 | Glyma.12g187200 | 17.8 | 16.4 | 35.1 | Bandillo et al. (2017) |
| Pubescence density | *P1* | Gm09 | 21.4 | Glyma.09g278000 | - | - | - | Bandillo et al. (2017) |
| Pubescence density | *Pd1* | Gm01 | 8.1 | Glyma.01g240100 | - | - | - | Bandillo et al. (2017) |
| Seed coat luster | *B* | Gm15 | 110.2 | Duplicated HPS genes | 15.6 | 12.4 | 26.2 | Bandillo et al. (2017) |
| Seed coat luster | *B?* | Gm09 | 34.2 | - | - | - | - | Bandillo et al. (2017) |
| Seed coat luster | *I* | Gm08 | 12.8 | CHS gene cluster | - | - | - | Bandillo et al. (2017) |
| Seed coat luster | *B1* | Gm13 | 7.7 | - | - | - | - | Bandillo et al. (2017) |
| Maturity group | *E2* | Gm10 | 49.7 | Glyma.10g221500 | - | - | - | Bandillo et al. (2017) |
| Maturity group | *E1* | Gm06 | 24 | Glyma.06g207800 | - | - | - | Bandillo et al. (2017) |
| Maturity group | *E3* | Gm19 | 21.5 | Glyma.19g224200 | - | - | - | Bandillo et al. (2017) |
| Maturity group | *E4* | Gm20 | 6.4 | Glyma.20g090000 | - | - | - | Bandillo et al. (2017) |
| Resistance to *P. sojae* | *cdwGm15* | Gm15 | 12.1 | Glyma.15g217100? | 10.7 | 9.1 | 13.7 | de Ronne et al. (2022) |
| Oil | *oilGm15* | Gm15 | 26.8 | Glyma.15g049200 | - | - | - | Bandillo et al. (2015) |
| Oil | *oilGm20* | Gm20 | 15.9 | Glyma.20g085100 | - | - | - | Bandillo et al. (2015) |
| Protein | *proGm20* | Gm20 | 32.3 | Glyma.20g085100 | - | - | - | Bandillo et al. (2015) |
| Protein | *proGm15* | Gm15 | 19.2 | Glyma.15g049200 | - | - | - | Bandillo et al. (2015) |

[a] Most significant -$\log_{10}(p)$ previously reported at this locus by the study in the Study column

[b] Gene associated with the locus. A question mark following the identifier of the gene indicates that this is simply a candidate that has yet to be confirmed. A dash indicates that no candidate has been suggested yet. We have only included candidate genes that have been suggested in previous studies.

[c] Most significant -$\log_{10}(p)$ reported from SNPs and indels at this locus. A dash indicates that no signal was detected from SNPs and indels at this locus. The same applies for other methods (SVs, $k$-mers) in their respective columns.

[d] The $p$-value at the W1 locus using the $k$-mers method was so small that it was numerically equal to zero. This value was therefore set to the -$\log_{10}$ of the smallest value that could be represented in R using our machine.

Table S2: Phenotype frequency and numerical coding of phenotypic data used for the GWAS analysis of flower color. "NA" indicates a missing value in the source data. Dashes indicate that this phenotype was not used in the GWAS analysis.

| Phenotype | Frequency | GWAS |
|---|---|---|
| Dark purple | 2 | - |
| Light purple | 2 | - |
| Purple | 191 | 1 |
| White | 154 | 2 |
| NA | 14 | - |

Table S3: Phenotype frequency and numerical coding of phenotypic data used in two GWAS analyses of pubescence color. "NA" indicates a missing value in the source data. Dashes indicate that this phenotype was not used in the GWAS analysis.

| Phenotype | Frequency | GWAS #1 | GWAS # 2 |
|---|---|---|---|
| Curly or glabrous | 2 | - | - |
| Gray | 157 | 1 | - |
| Light tawny | 37 | 3 | 2 |
| Near gray | 5 | - | - |
| Tawny | 148 | 2 | 1 |
| NA | 14 | - | - |

Table S4: Phenotype frequency and numerical coding of phenotypic data used in two GWAS analyses of seed coat color. "NA" indicates a missing value in the source data. Dashes indicate that this phenotype was not used in the GWAS analysis.

| Phenotype | Frequency | GWAS #1 | GWAS # 2 |
|---|---|---|---|
| Black | 56 | 5 | - |
| Brown | 17 | 3 | - |
| Gray | 4 | - | - |
| Grayish green | 1 | - | - |
| Green | 17 | 2 | 2 |
| Greenish brown | 9 | - | - |
| Light green | 4 | - | - |
| Reddish brown | 8 | 4 | - |
| Yellow | 244 | 1 | 1 |
| NA | 3 | - | - |

Table S5: Phenotype frequency and numerical coding of phenotypic data used in two GWAS analyses of stem termination type. "NA" indicates a missing value in the source data. Dashes indicate that this phenotype was not used in the GWAS analysis.

| Phenotype | Frequency | GWAS #1 | GWAS # 2 |
|---|---|---|---|
| Determinate | 89 | 1 | - |
| Indeterminate | 232 | 3 | 2 |
| Semi-determinate | 28 | 2 | 1 |
| NA | 14 | - | - |

Table S6: Phenotype frequency and numerical coding of phenotypic data used in three GWAS analyses of hilum color. "NA" indicates a missing value in the source data. Dashes indicate that this phenotype was not used in the GWAS analysis.

| Phenotype | Frequency | GWAS #1 | GWAS # 2 | GWAS # 3 |
|---|---|---|---|---|
| Buff | 100 | 3 | - | - |
| Black | 99 | 7 | 2 | - |
| Black hilum with brown outer ring | 3 | - | - | - |
| Brown | 61 | 4 | 1 | 1 |
| Brown with black | 7 | - | - | - |
| Brown with black/black | 1 | - | - | - |
| Dark buff | 1 | - | - | - |
| Dark gray | 1 | - | - | - |
| Dark imperfect black | 1 | - | - | - |
| Gray | 5 | 2 | - | - |
| Green | 2 | - | - | - |
| Greenish brown | 2 | - | - | - |
| Imperfect black | 21 | 6 | - | - |
| Imperfect gray | 1 | - | - | - |
| Light buff | 7 | - | - | - |
| Reddish buff | 1 | - | - | - |
| Reddish brown | 15 | 5 | - | 2 |
| Tan | 6 | - | - | - |
| Yellow | 25 | 1 | - | - |
| Yellow/Buff | 1 | - | - | - |
| NA | 3 | - | - | - |

Table S7: Phenotype frequency and numerical coding of phenotypic data used in two GWAS analyses of pod color. "NA" indicates a missing value in the source data. Dashes indicate that this phenotype was not used in the GWAS analysis.

| Phenotype | Frequency | GWAS #1 | GWAS # 2 |
|---|---|---|---|
| Black | 40 | 4 | 2 |
| Brown | 219 | 2 | 1 |
| Dark brown | 11 | 3 | - |
| Heterogeneous | 1 | - | - |
| Light brown | 4 | - | - |
| Tan | 85 | 1 | - |
| NA | 3 | - | - |

Table S8: Phenotype frequency and numerical coding of phenotypic data used in two GWAS analyses of pubescence form. "NA" indicates a missing value in the source data. Dashes indicate that this phenotype was not used in the GWAS analysis.

| Phenotype | Frequency | GWAS #1 | GWAS # 2 |
|---|---|---|---|
| Appressed | 38 | 1 | 1 |
| Curly | 2 | - | - |
| Erect | 259 | 3 | - |
| Semi-appressed | 51 | 2 | 2 |
| NA | 13 | - | - |

Table S9: Phenotype frequency and numerical coding of phenotypic data used for the GWAS analysis of pubescence density. “NA” indicates a missing value in the source data. Dashes indicate that this phenotype was not used in the GWAS analysis.

| Phenotype | Frequency | GWAS |
|---|---|---|
| Dense | 2 | - |
| Glabrous | 1 | - |
| Normal | 280 | 1 |
| Sparse | 2 | - |
| Semi-sparse | 73 | 2 |
| Semi-sparse/normal | 1 | - |
| NA | 4 | - |

Table S10: Phenotype frequency and numerical coding of phenotypic data used in three GWAS analyses of seed coat luster. "NA" indicates a missing value in the source data. Dashes indicate that this phenotype was not used in the GWAS analysis.

| Phenotype | Frequency | GWAS #1 | GWAS # 2 | GWAS # 3 |
|---|---|---|---|---|
| Bloom | 10 | 1 | 1 | - |
| Dull | 67 | 2 | 2 | 1 |
| Intermediate | 177 | 3 | - | - |
| Intermediate/Bloom | 1 | - | - | - |
| Light bloom | 9 | 4 | 3 | - |
| Shiny | 84 | 5 | 4 | 2 |
| Shiny/intermediate | 1 | - | - | - |
| NA | 14 | - | - | - |

Table S11: Phenotype frequency and numerical coding of phenotypic data used for the GWAS analysis of maturity group. “NA” indicates a missing value in the source data. Dashes indicate that this phenotype was not used in the GWAS analysis.

| Phenotype | Frequency | GWAS |
|---|---|---|
| II | 13 | 1 |
| III | 120 | 2 |
| III/IV | 1 | - |
| IV | 153 | 3 |
| V | 62 | 4 |
| NA | 14 | - |

### Supplemental figures

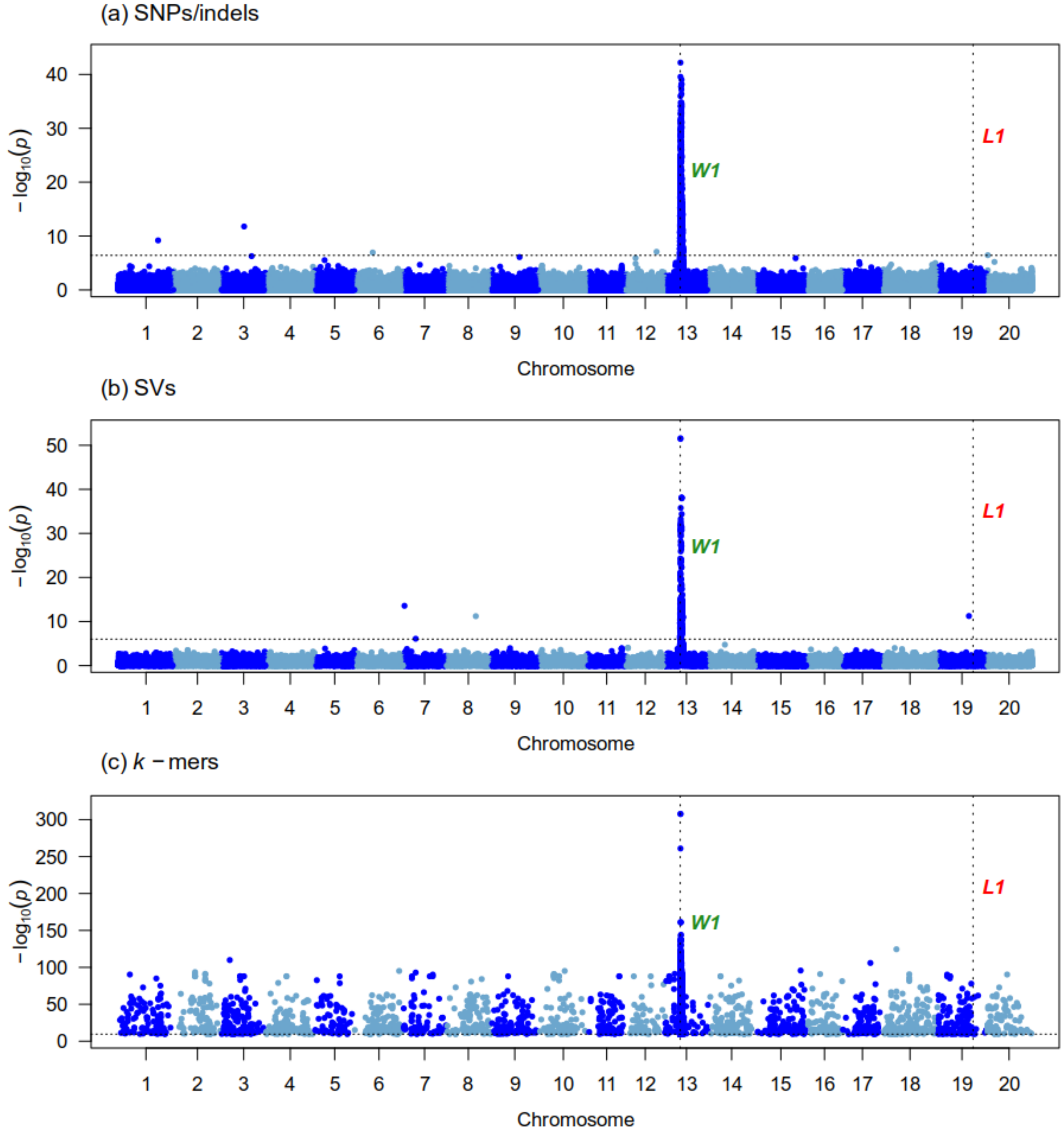


Figure S1: Manhattan plots generated from the GWAS analysis of flower color on 345 samples using three genotype datasets : (a) Platypus (SNPs and indels), (b) Paragraph (SVs), (c) $k$-mers presence/absence. The x-axis shows the position along the reference assembly version 4 of Williams82. Each point shows the $-\log_{10}(p)$ associated with a particular marker or $k$-mer. Horizontal dotted lines indicate the 5% family-wise error-rate significance threshold determined from a randomization approach. Vertical dotted lines indicate the position of signals associated with the trait. Documented loci are colored according to whether they were found (green) or not (red) by a particular approach. The "Gm" prefix has been left out of chromosome names for simplicity.

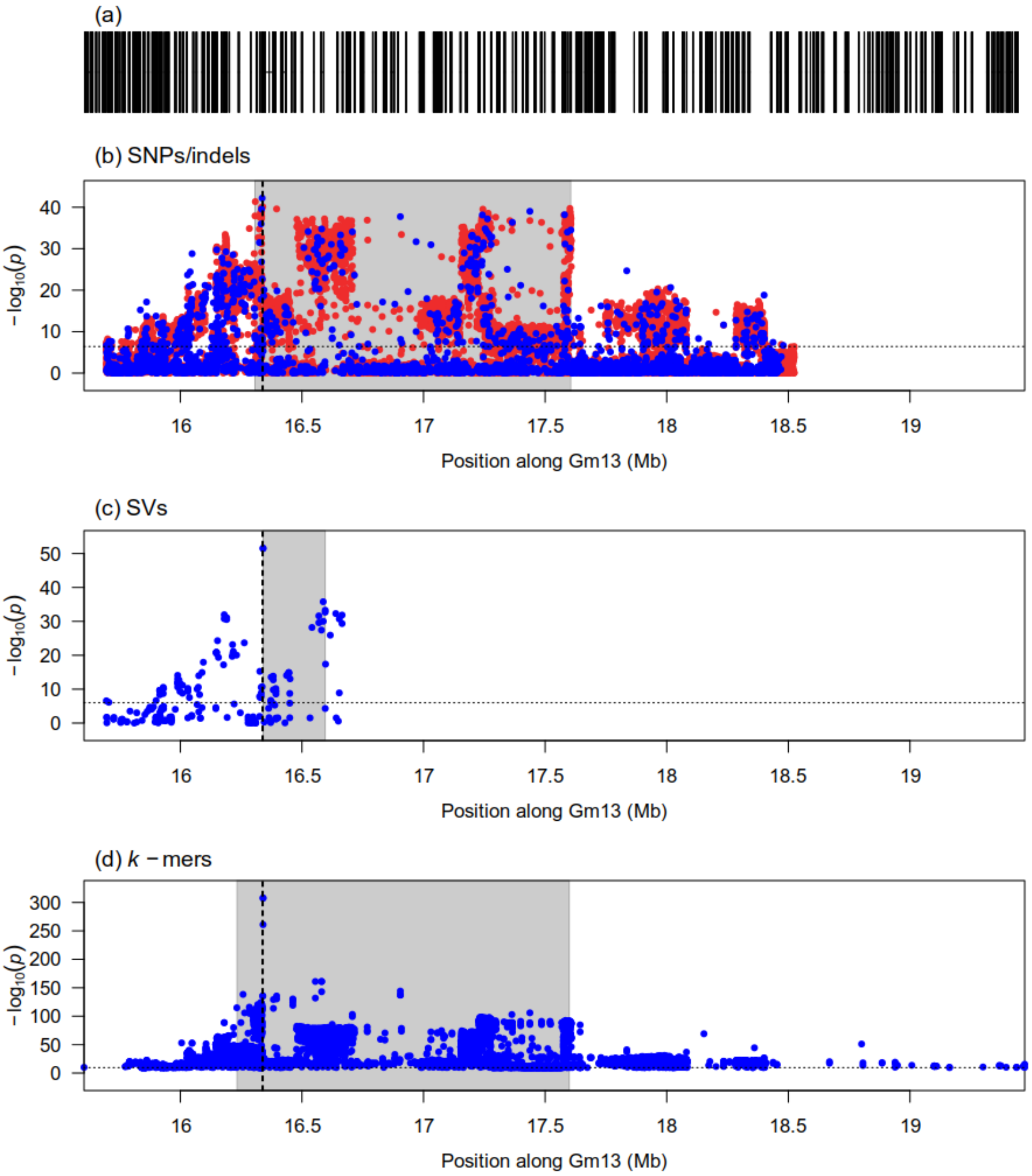


Figure S2: Zoomed-in Manhattan plots of signals detected by the GWAS analysis of flower color at the *W1* locus using three genotype datasets: (b) Platypus (SNPs and indels), (c) Paragraph (SVs), (d) *k*-mers presence/absence. Panel (a) shows gene models over the genomic interval. Horizontal dotted lines indicate the 5% family-wise error-rate significance threshold determined from a randomization approach. Vertical dotted lines indicate the location of the Glyma.13g072100 gene associated with the locus. Gray shaded rectangles indicate the region delimited by the top 5% (SVs) or top 1% (SNPs/indels and *k*-mers) associations in the signal region. In the case of SNPs/indels, blue points denote markers used in the original analysis, whereas red points denote markers that had originally been pruned but whose *p*-values were computed after signal discovery.

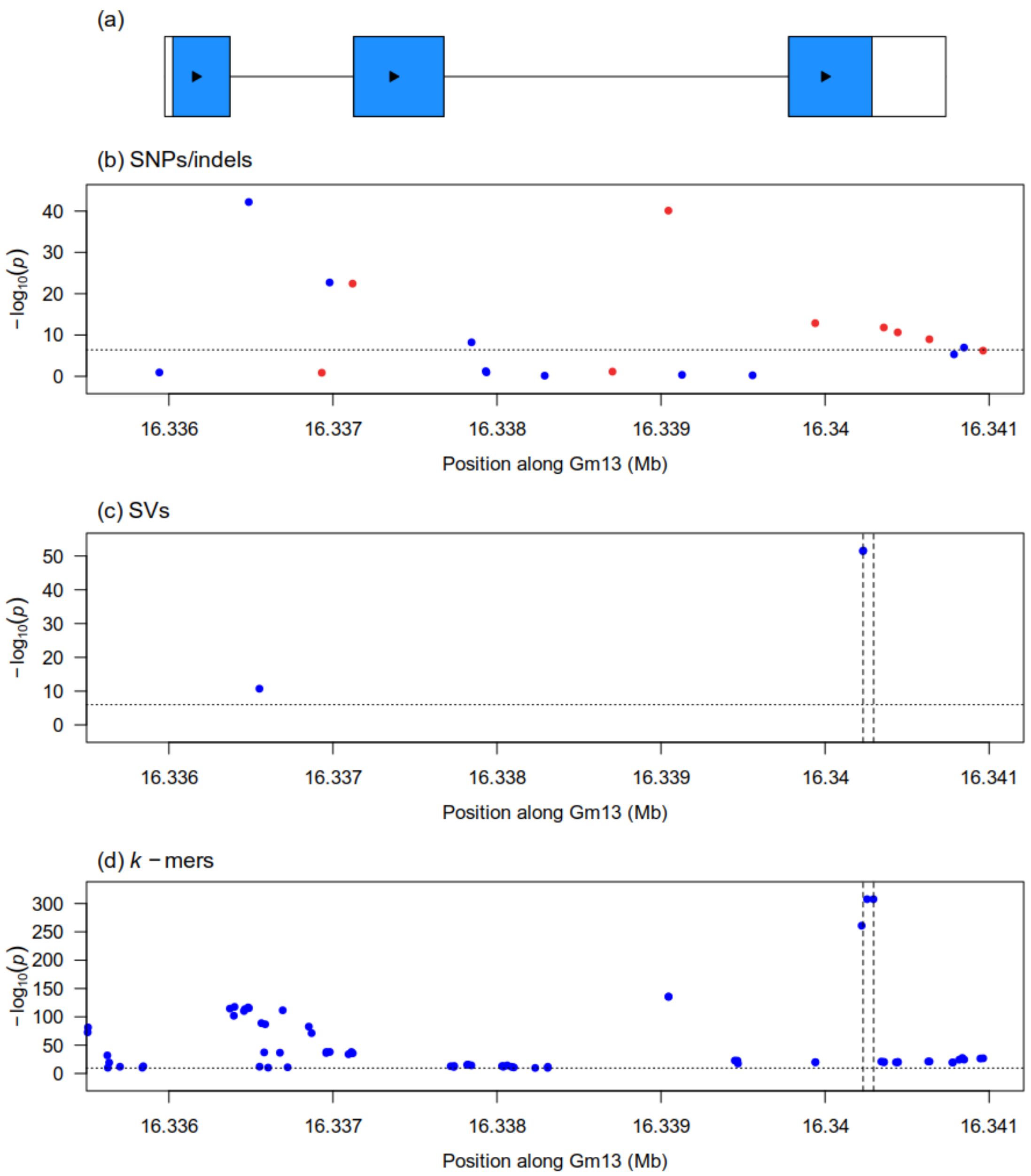


Figure S3: Zoomed-in Manhattan plots generated from the GWAS analysis of flower color at the Glyma.13g072100 gene associated with the *W1* locus using three genotype datasets: (b) Platypus (SNPs and indels), (c) Paragraph (SVs), (d) *k*-mers presence/absence. Vertical dotted lines in panels (c) and (d) indicate the location of the causal SV at this locus. Horizontal dotted lines indicate the 5% family-wise error-rate significance threshold determined from a randomization approach. In the case of SNPs/indels, blue points denote markers used in the original analysis, whereas red points denote markers that had originally been pruned but whose *p*-values were computed after signal discovery. Panel (a) shows gene models over the plotting interval. Exons are represented by rectangles whereas introns are represented by horizontal lines. Coding sequences are shown in blue and the direction of transcription is indicated by arrows.

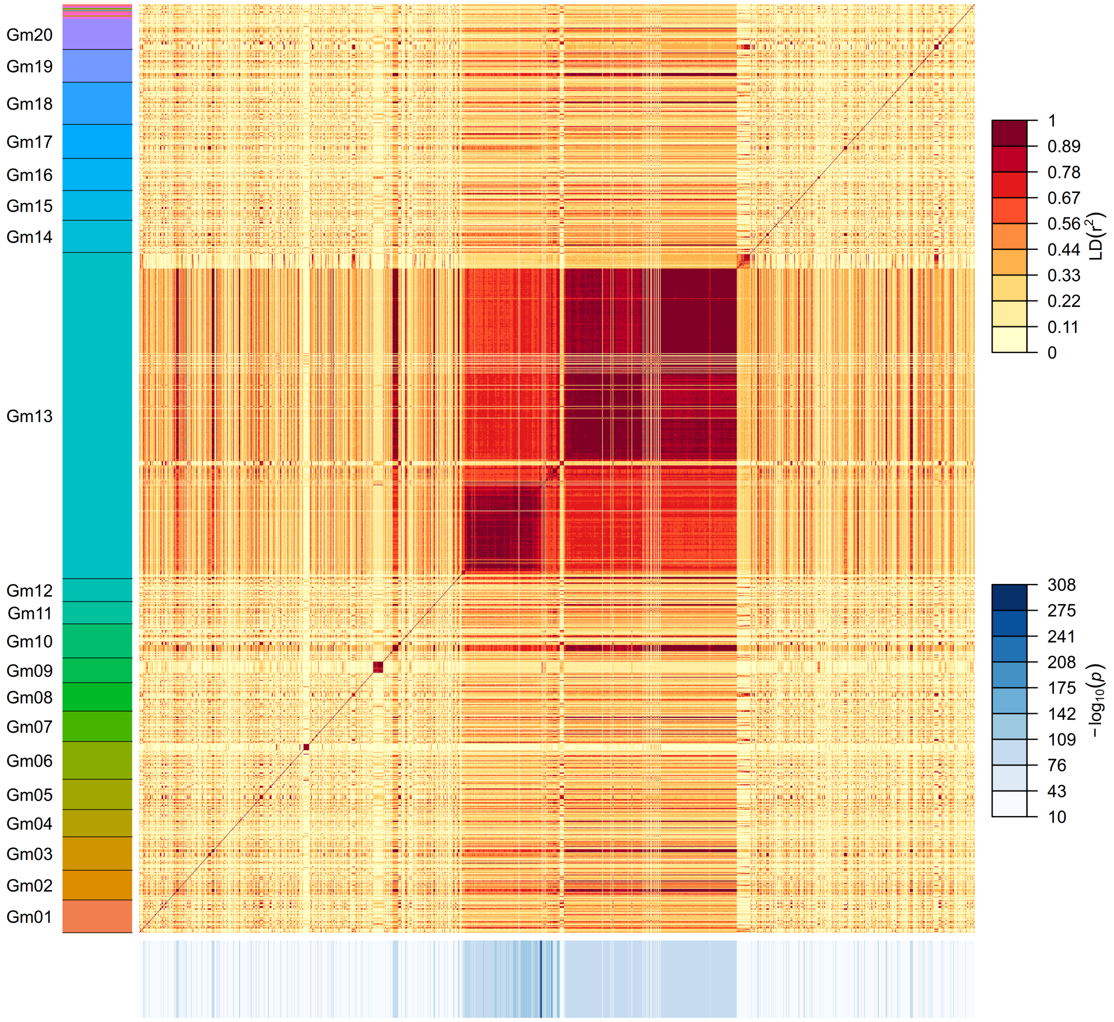


Figure S4: Pairwise LD among 1500 significant $k$-mers identified for flower color. $k$-mers are sorted along the y-axis according to their putative position along the reference assembly version 4 of Williams82, as identified by "Gm" chromosome labels. Sequences that lack a "Gm" label (top of the y-axis) represent unanchored scaffolds. $k$-mers are represented in the same order along the x- and y-axis. The colored rectangles drawn below the x-axis represent the $-\log_{10}(p)$ of each $k$-mer.

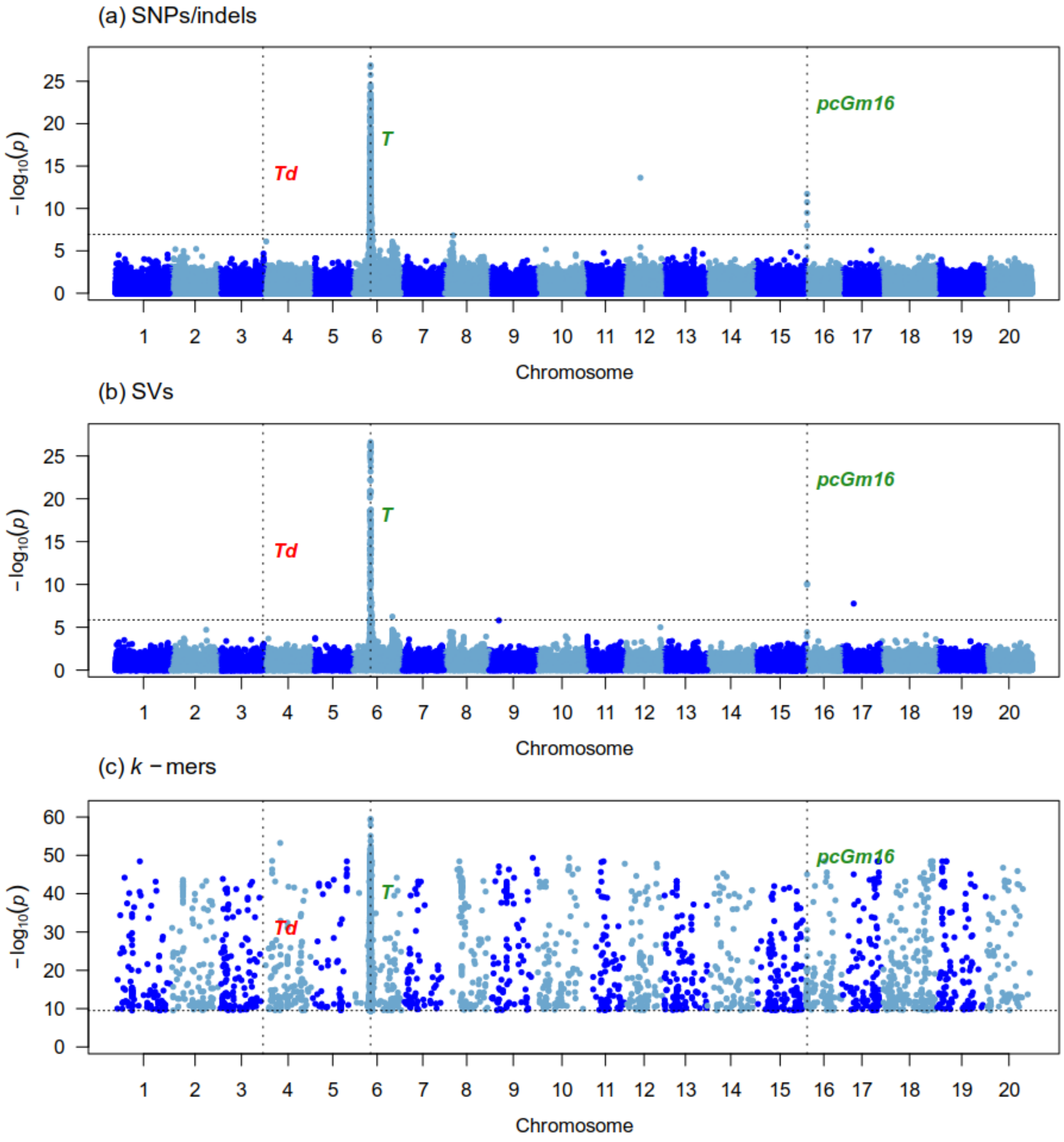


Figure S5: Manhattan plots generated from the GWAS analysis of pubescence color (first GWAS) on 342 samples using three genotype datasets : (a) Platypus (SNPs and indels), (b) Paragraph (SVs), (c) $k$-mers presence/absence. The x-axis shows the position along the reference assembly version 4 of Williams82. Each point shows the $-\log_{10}(p)$ associated with a particular marker or $k$-mer. Horizontal dotted lines indicate the 5% family-wise error-rate significance threshold determined from a randomization approach. Vertical dotted lines indicate the position of signals associated with the trait. Documented loci are colored according to whether they were found (green) or not (red) by a particular approach. The "Gm" prefix has been left out of chromosome names for simplicity.

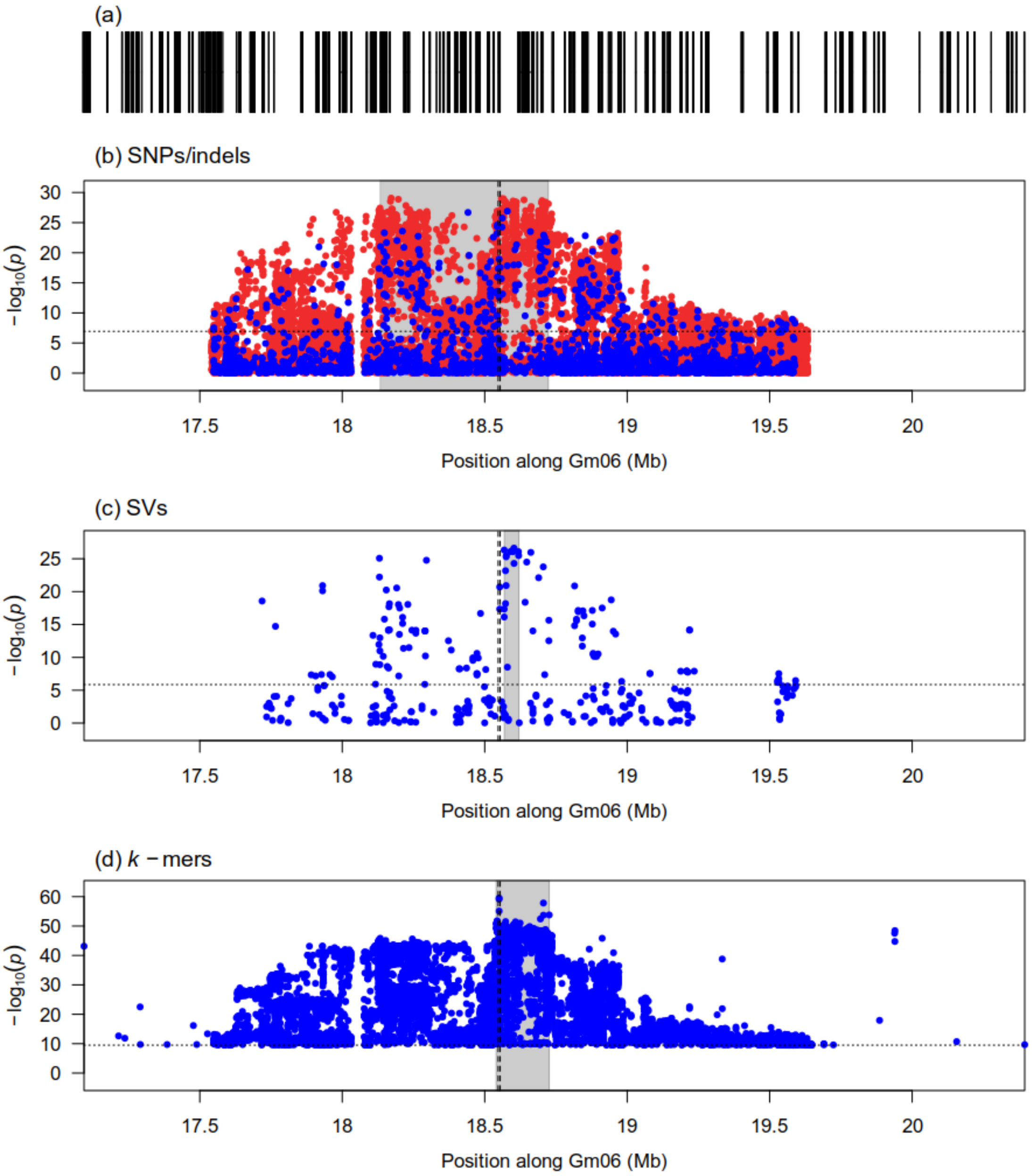


Figure S6: Zoomed-in Manhattan plots of signals detected by the GWAS analysis of pubescence color (first GWAS) at the $T$ locus using three genotype datasets: (b) Platypus (SNPs and indels), (c) Paragraph (SVs), (d) $k$-mers presence/absence. Panel (a) shows gene models over the genomic interval. Horizontal dotted lines indicate the 5% family-wise error-rate significance threshold determined from a randomization approach. Vertical dotted lines indicate the location of the Glyma.06g202300 gene associated with the locus. Gray shaded rectangles indicate the region delimited by the top 5% (SVs) or top 1% (SNPs/indels and $k$-mers) associations in the signal region. In the case of SNPs/indels, blue points denote markers used in the original analysis, whereas red points denote markers that had originally been pruned but whose $p$-values were computed after signal discovery.

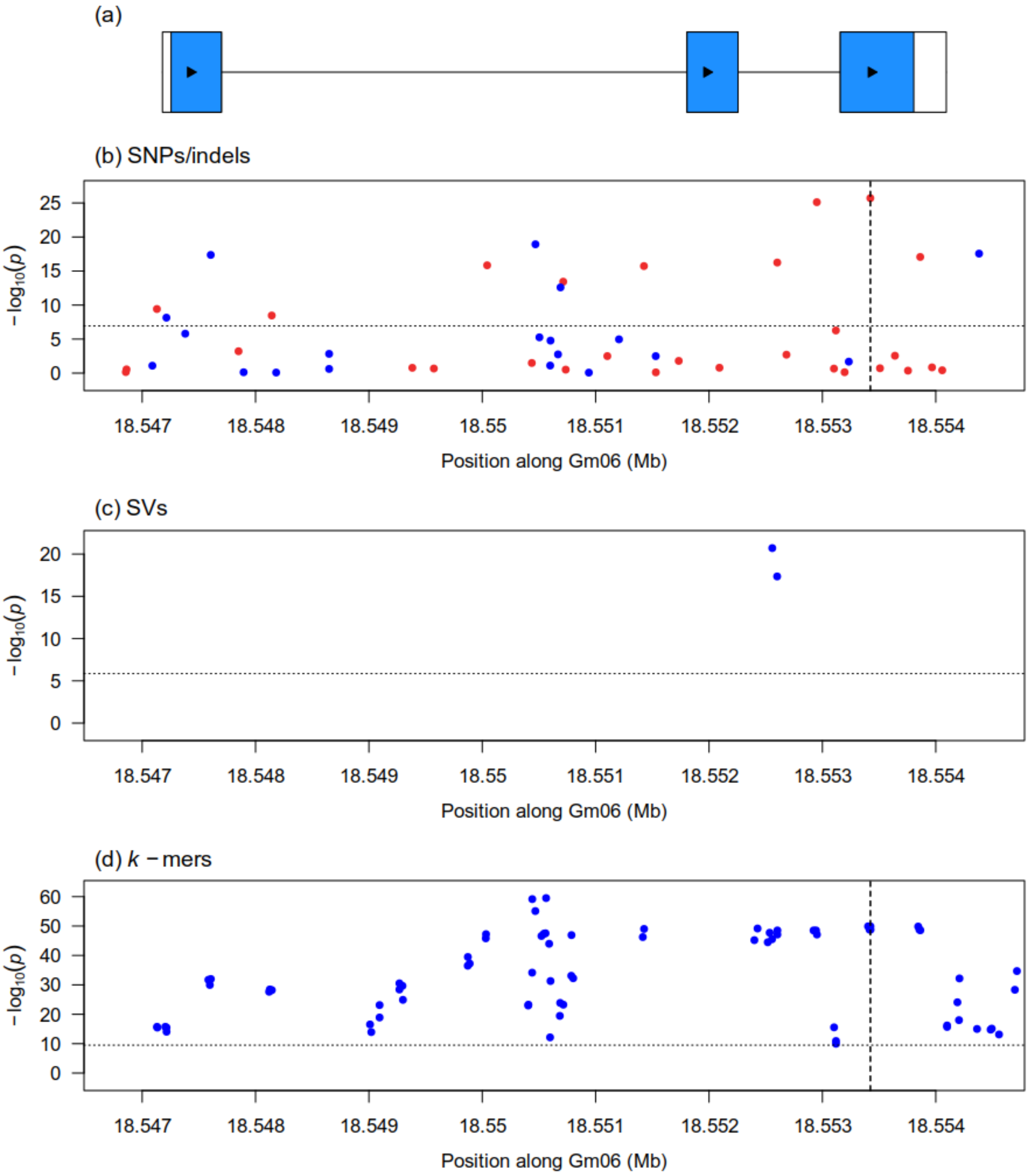


Figure S7: Zoomed-in Manhattan plots generated from the GWAS analysis of pubescence color (first GWAS) at the Glyma.06g202300 gene associated with the $T$ locus using three genotype datasets: (b) Platypus (SNPs and indels), (c) Paragraph (SVs), (d) $k$-mers presence/absence. Vertical dotted lines in panels (b) and (d) indicate the location of the causal indel at this locus. Horizontal dotted lines indicate the 5% family-wise error-rate significance threshold determined from a randomization approach. In the case of SNPs/indels, blue points denote markers used in the original analysis, whereas red points denote markers that had originally been pruned but whose $p$-values were computed after signal discovery. Panel (a) shows gene models over the plotting interval. Exons are represented by rectangles whereas introns are represented by horizontal lines. Coding sequences are shown in blue and the direction of transcription is indicated by arrows.

(a)

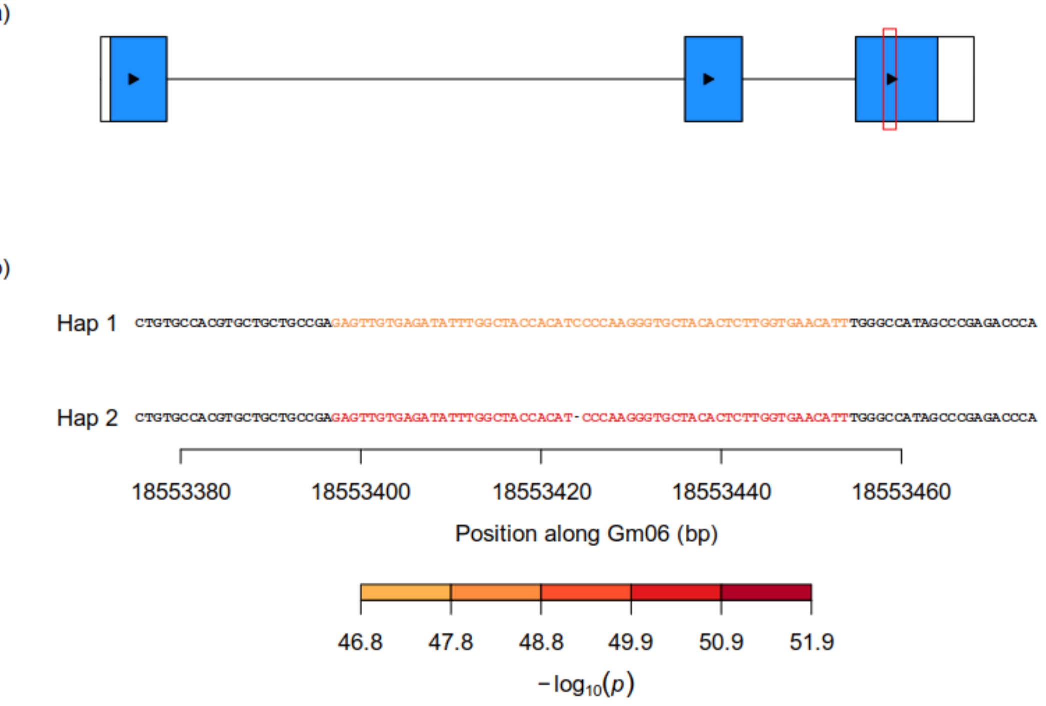


(c)

| | Hap 1 | Hap 2 |
|---|---|---|
| Gray | 22 | 134 |
| Light tawny | 37 | 0 |
| Tawny | 148 | 0 |

Figure S8: Identification of a causal indel underlying significant $k$-mers at the Glyma.06g202300 gene associated with the $T$ locus for pubescence color (first GWAS). (a) Gene model of Glyma.06g202300. Exons are represented by rectangles whereas introns are represented by horizontal lines. Coding sequences are shown in blue and the direction of transcription is indicated by arrows. The red rectangle identifies the region that is zoomed-in in panel (b). (b) Nucleotide sequences of haplotypes observed in at least five samples across the dataset. Individual nucleotides are colored according to the $-\log_{10}(p)$ of the most significant $k$-mer overlapping them. Dashes indicate gaps in haplotype sequence alignment whereas vertical lines indicate differences in sequence between two haplotypes. (c) Contingency table of the phenotypes and haplotypes observed in the dataset. Haplotypes correspond to those shown in panel (b).

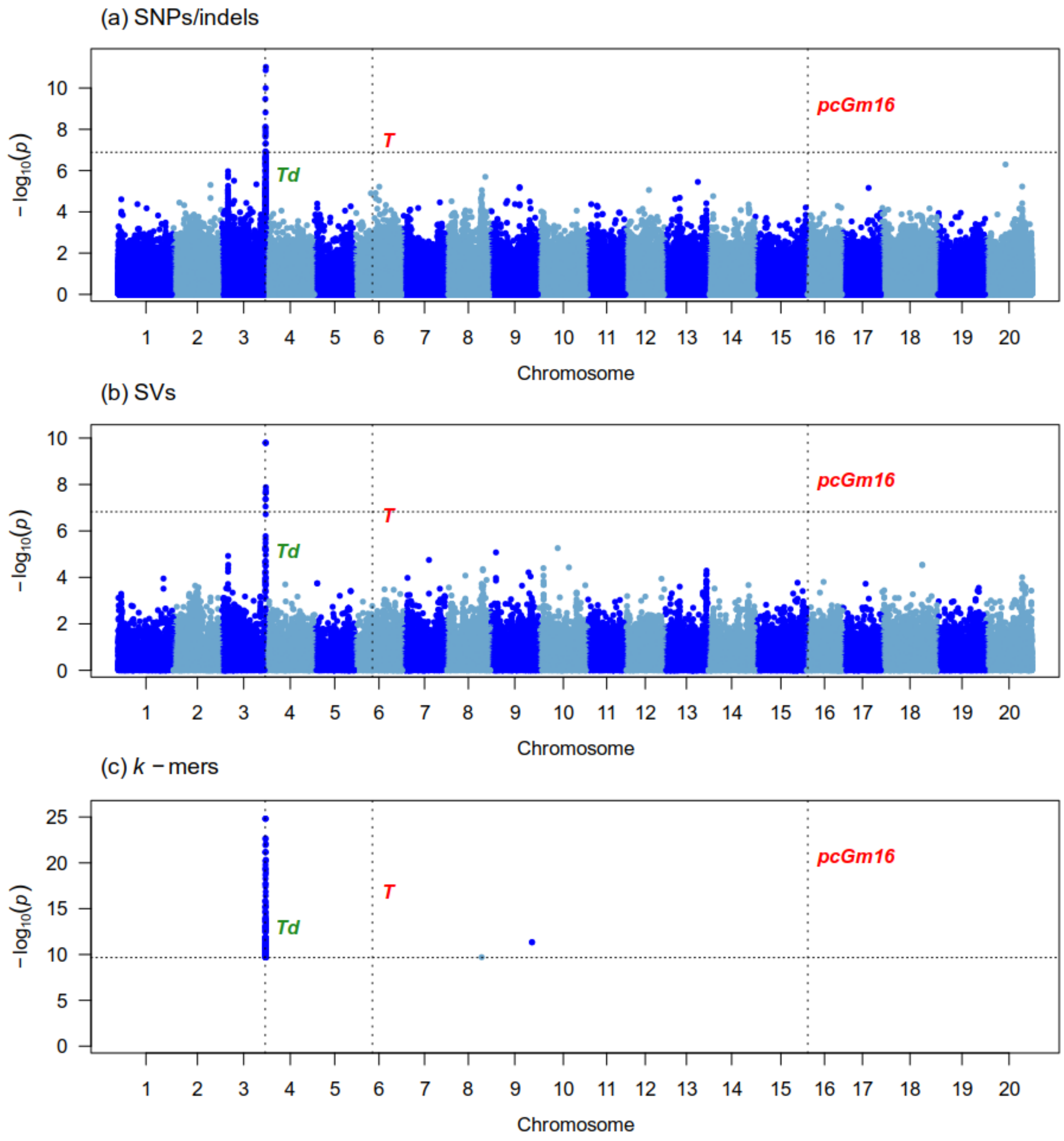


Figure S9: Manhattan plots generated from the GWAS analysis of pubescence color (second GWAS) on 185 samples using three genotype datasets : (a) Platypus (SNPs and indels), (b) Paragraph (SVs), (c) $k$-mers presence/absence. The x-axis shows the position along the reference assembly version 4 of Williams82. Each point shows the $-\log_{10}(p)$ associated with a particular marker or $k$-mer. Horizontal dotted lines indicate the 5% family-wise error-rate significance threshold determined from a randomization approach. Vertical dotted lines indicate the position of signals associated with the trait. Documented loci are colored according to whether they were found (green) or not (red) by a particular approach. The "Gm" prefix has been left out of chromosome names for simplicity.

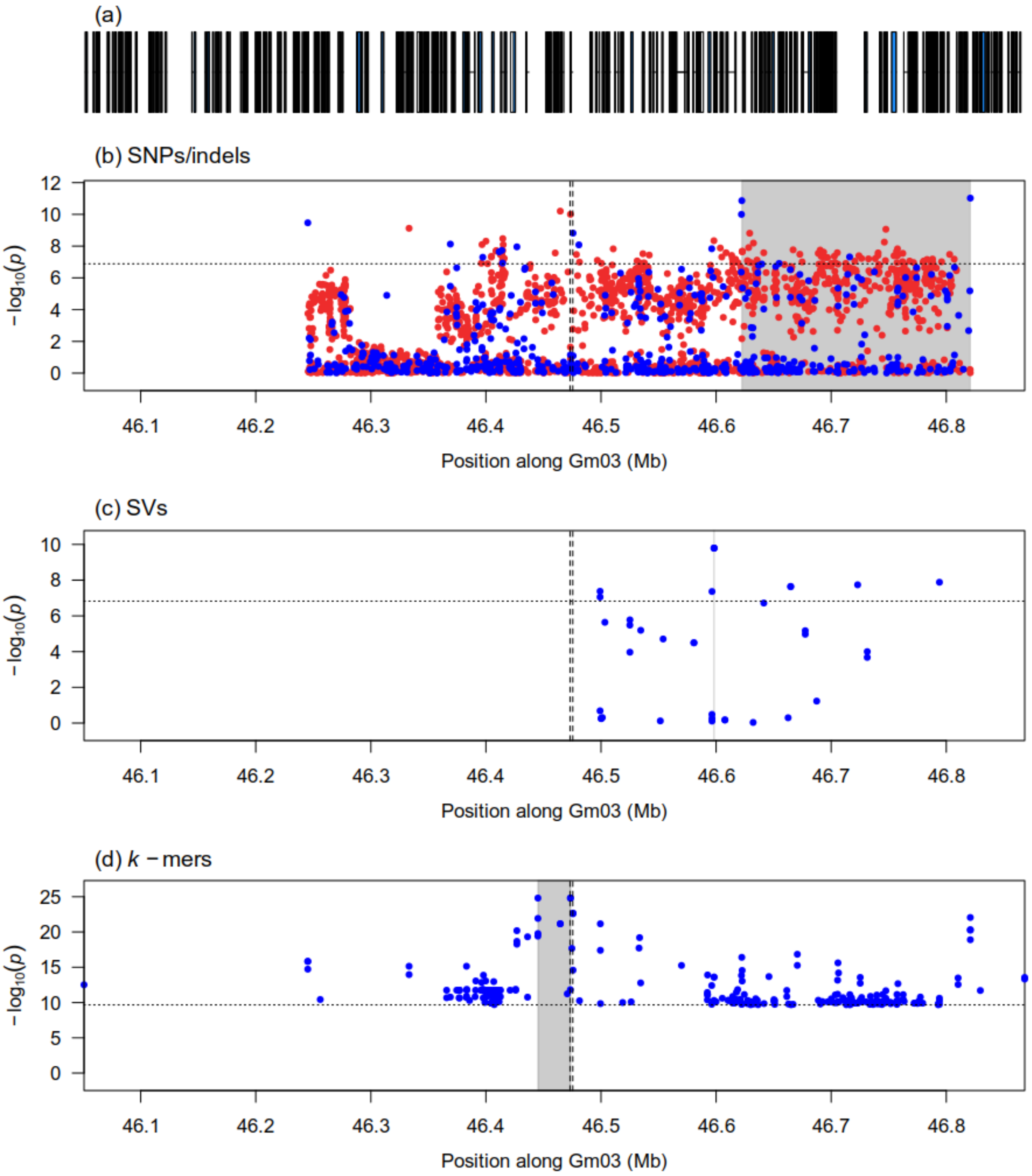


Figure S10: Zoomed-in Manhattan plots of signals detected by the GWAS analysis of pubescence color (second GWAS) at the *Td* locus using three genotype datasets: (b) Platypus (SNPs and indels), (c) Paragraph (SVs), (d) $k$-mers presence/absence. Panel (a) shows gene models over the genomic interval. Horizontal dotted lines indicate the 5% family-wise error-rate significance threshold determined from a randomization approach. Vertical dotted lines indicate the location of the Glyma.03g258700 gene associated with the locus. Gray shaded rectangles indicate the region delimited by the top 5% (SVs) or top 1% (SNPs/indels and $k$-mers) associations in the signal region. In the case of SNPs/indels, blue points denote markers used in the original analysis, whereas red points denote markers that had originally been pruned but whose $p$-values were computed after signal discovery.

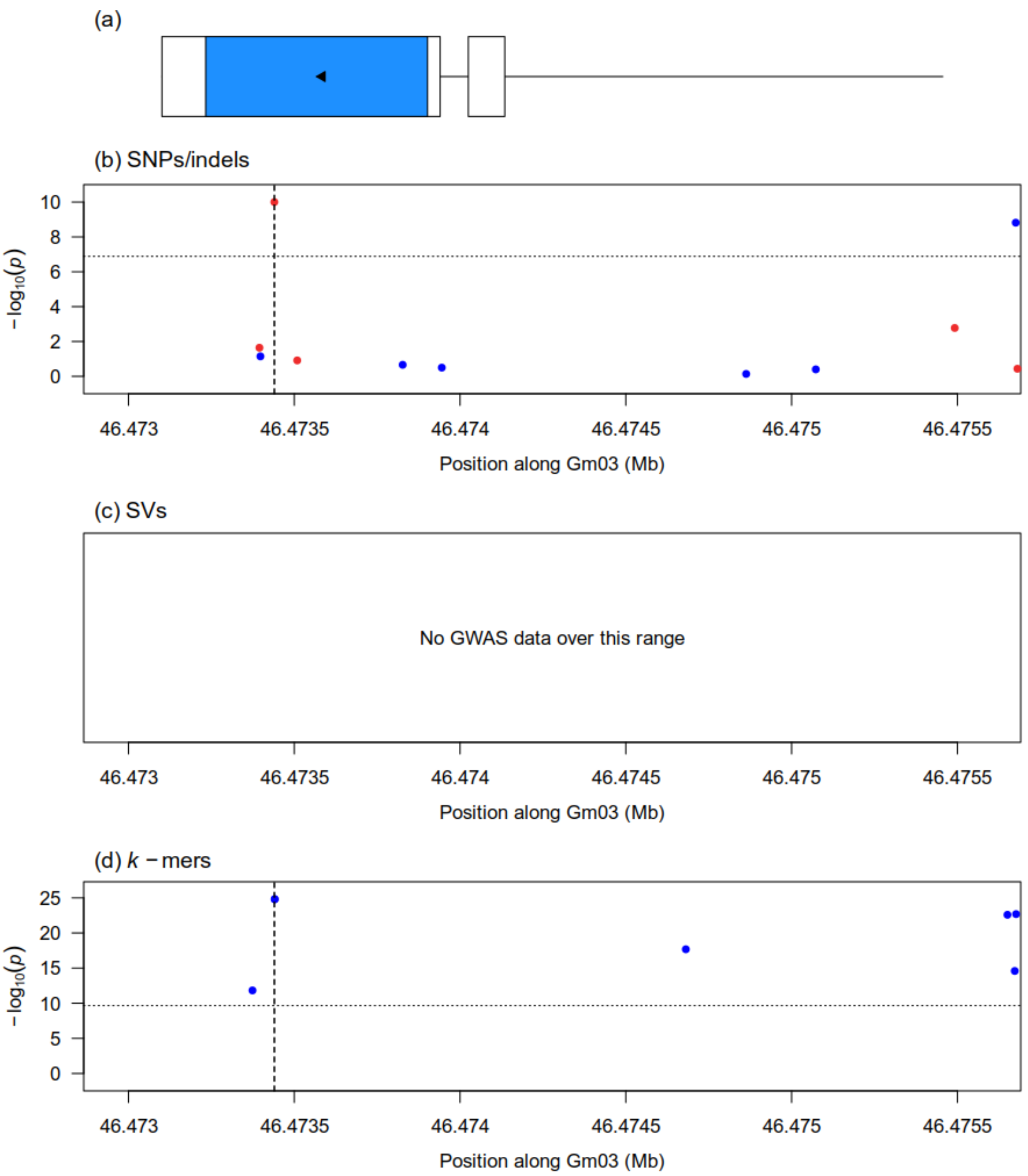


Figure S11: Zoomed-in Manhattan plots generated from the GWAS analysis of pubescence color (second GWAS) at the Glyma.03g258700 gene associated with the *Td* locus using three genotype datasets: (b) Platypus (SNPs and indels), (c) Paragraph (SVs), (d) *k*-mers presence/absence. Vertical dotted lines in panels (b) and (d) indicate the location of the causal SNP at this locus. Horizontal dotted lines indicate the 5% family-wise error-rate significance threshold determined from a randomization approach. In the case of SNPs/indels, blue points denote markers used in the original analysis, whereas red points denote markers that had originally been pruned but whose *p*-values were computed after signal discovery. Panel (a) shows gene models over the plotting interval. Exons are represented by rectangles whereas introns are represented by horizontal lines. Coding sequences are shown in blue and the direction of transcription is indicated by arrows.

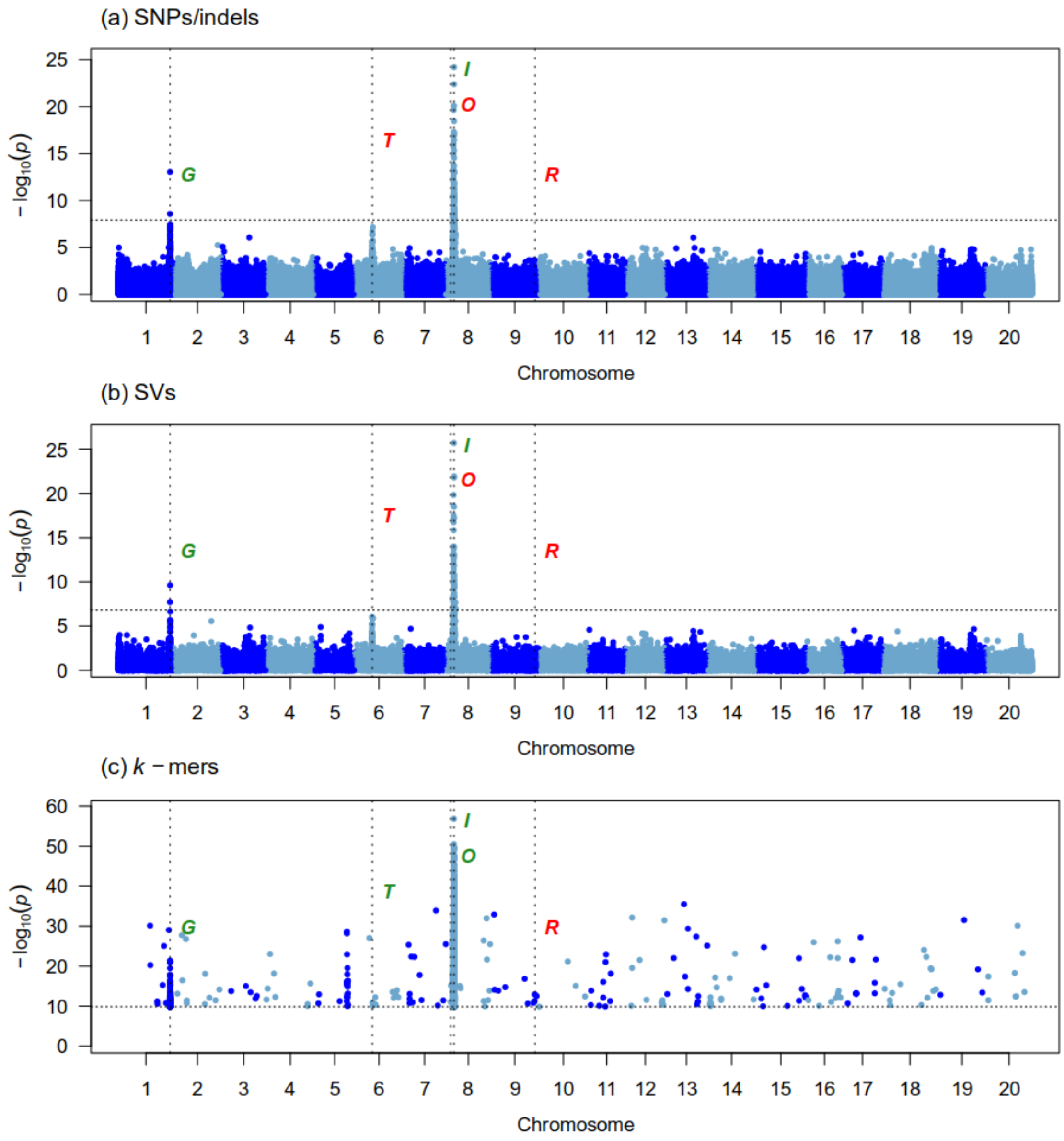


Figure S12: Manhattan plots generated from the GWAS analysis of seed coat color (first GWAS) on 342 samples using three genotype datasets : (a) Platypus (SNPs and indels), (b) Paragraph (SVs), (c) $k$-mers presence/absence. The x-axis shows the position along the reference assembly version 4 of Williams82. Each point shows the $-\log_{10}(p)$ associated with a particular marker or $k$-mer. Horizontal dotted lines indicate the 5% family-wise error-rate significance threshold determined from a randomization approach. Vertical dotted lines indicate the position of signals associated with the trait. Documented loci are colored according to whether they were found (green) or not (red) by a particular approach. The "Gm" prefix has been left out of chromosome names for simplicity.

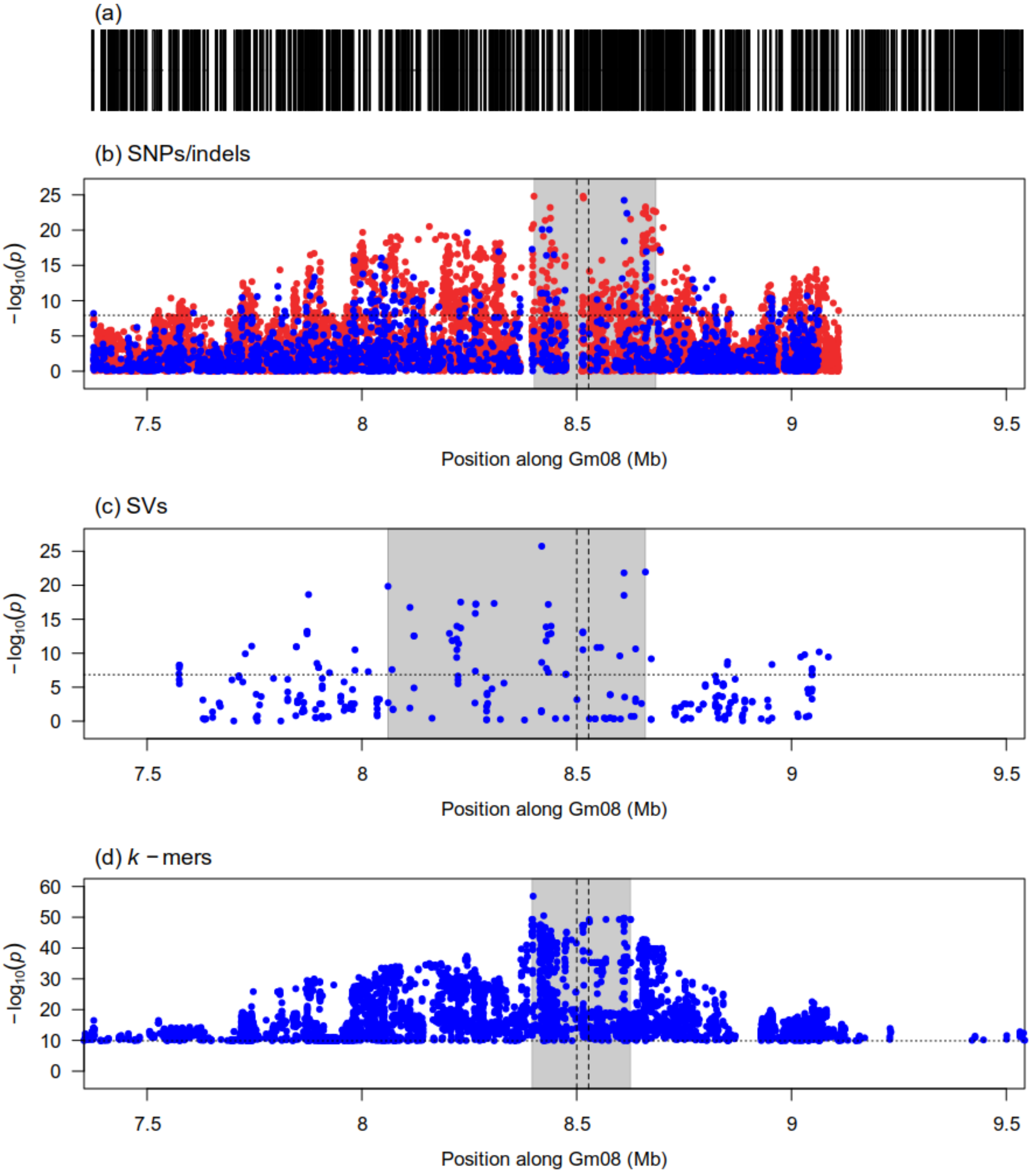


Figure S13: Zoomed-in Manhattan plots of signals detected by the GWAS analysis of seed coat color (first GWAS) at the *I* locus using three genotype datasets: (b) Platypus (SNPs and indels), (c) Paragraph (SVs), (d) $k$-mers presence/absence. Panel (a) shows gene models over the genomic interval. Horizontal dotted lines indicate the 5% family-wise error-rate significance threshold determined from a randomization approach. Vertical dotted lines indicate the boundaries of the tandem duplication/inversion identified as the causal variant at this locus. Gray shaded rectangles indicate the region delimited by the top 5% (SVs) or top 1% (SNPs/indels and $k$-mers) associations in the signal region. In the case of SNPs/indels, blue points denote markers used in the original analysis, whereas red points denote markers that had originally been pruned but whose $p$-values were computed after signal discovery.

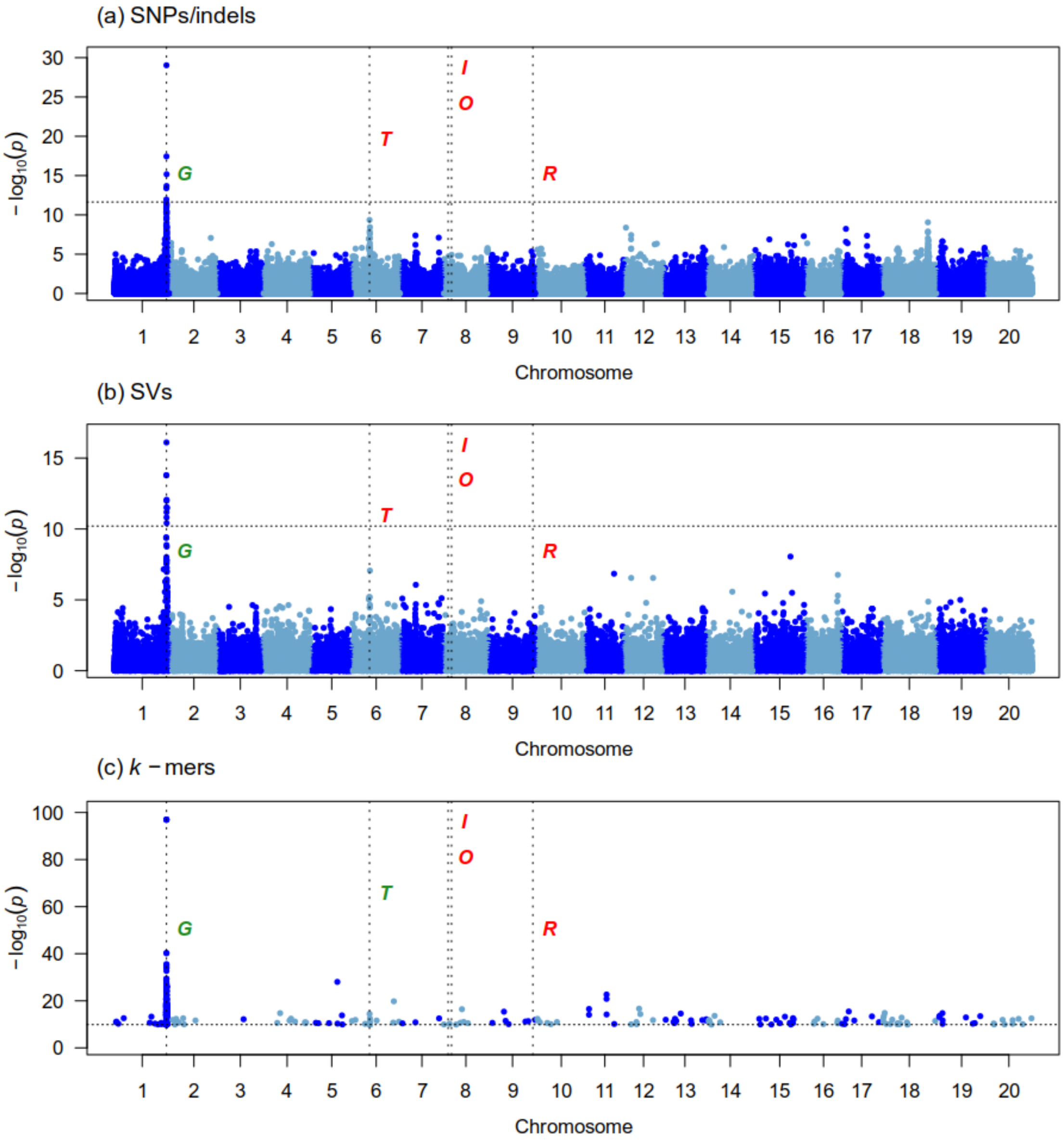


Figure S14: Manhattan plots generated from the GWAS analysis of seed coat color (second GWAS) on 261 samples using three genotype datasets : (a) Platypus (SNPs and indels), (b) Paragraph (SVs), (c) $k$-mers presence/absence. The x-axis shows the position along the reference assembly version 4 of Williams82. Each point shows the $-\log_{10}(p)$ associated with a particular marker or $k$-mer. Horizontal dotted lines indicate the 5% family-wise error-rate significance threshold determined from a randomization approach. Vertical dotted lines indicate the position of signals associated with the trait. Documented loci are colored according to whether they were found (green) or not (red) by a particular approach. The "Gm" prefix has been left out of chromosome names for simplicity.

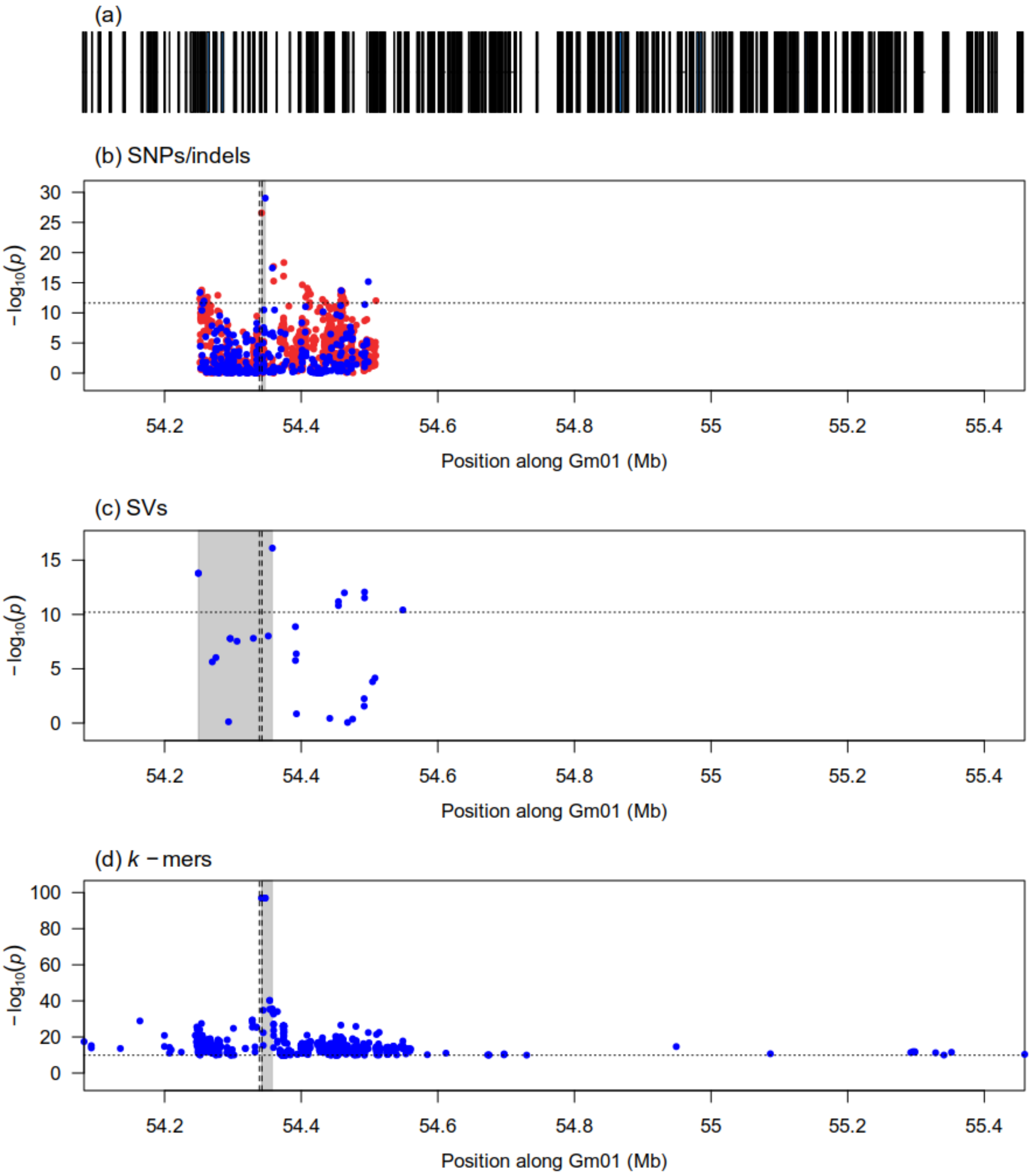


Figure S15: Zoomed-in Manhattan plots of signals detected by the GWAS analysis of seed coat color (second GWAS) at the $G$ locus using three genotype datasets: (b) Platypus (SNPs and indels), (c) Paragraph (SVs), (d) $k$-mers presence/absence. Panel (a) shows gene models over the genomic interval. Horizontal dotted lines indicate the 5% family-wise error-rate significance threshold determined from a randomization approach. Vertical dotted lines indicate the location of the Glyma.01g198500 gene associated with the locus. Gray shaded rectangles indicate the region delimited by the top 5% (SVs) or top 1% (SNPs/indels and $k$-mers) associations in the signal region. In the case of SNPs/indels, blue points denote markers used in the original analysis, whereas red points denote markers that had originally been pruned but whose $p$-values were computed after signal discovery.

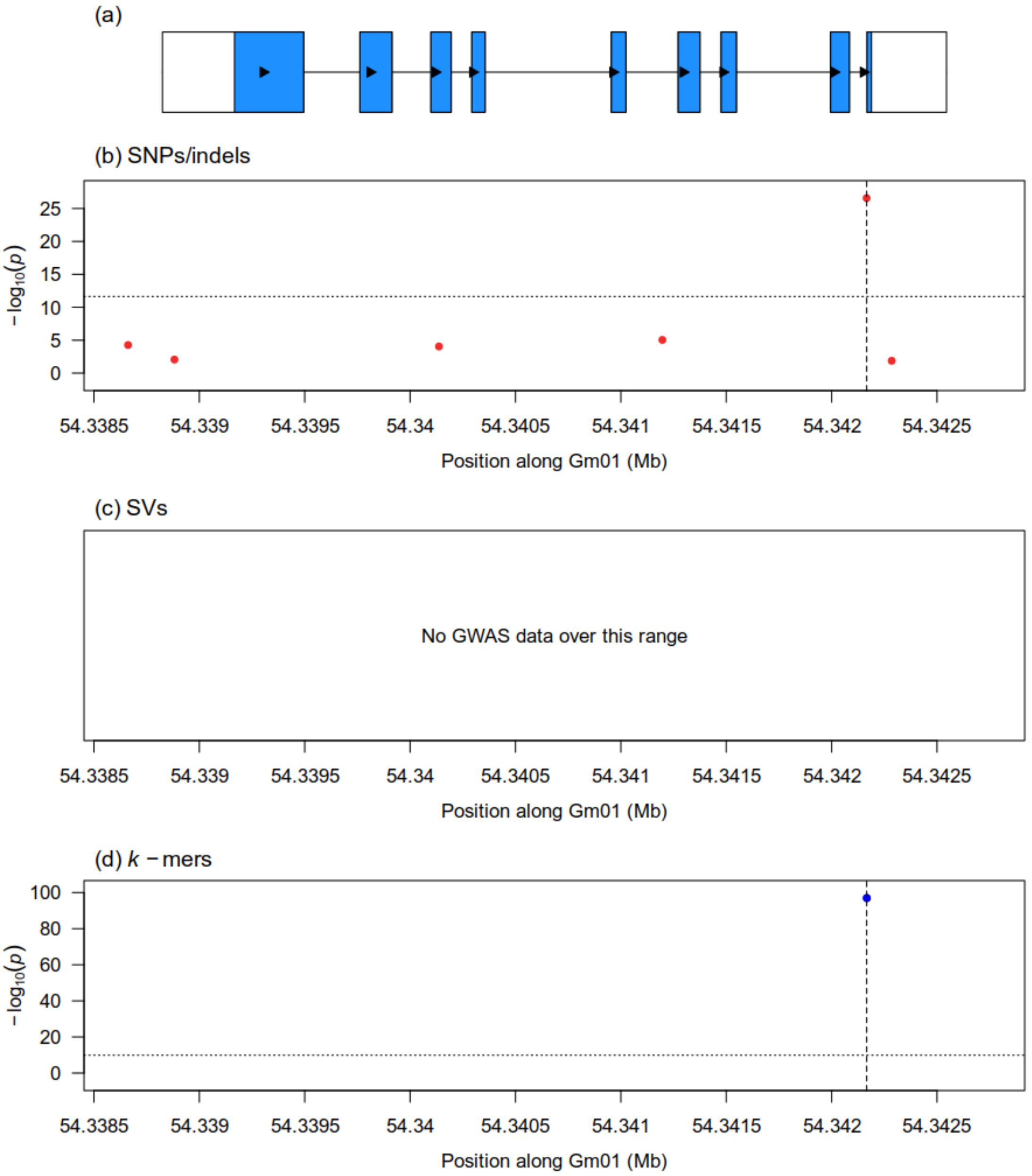


Figure S16: Zoomed-in Manhattan plots generated from the GWAS analysis of seed coat color (second GWAS) at the Glyma.01g198500 gene associated with the *G* locus using three genotype datasets: (b) Platypus (SNPs and indels), (c) Paragraph (SVs), (d) $k$-mers presence/absence. Vertical dotted lines in panels (b) and (d) indicate the location of the causal SNP at this locus. Horizontal dotted lines indicate the 5% family-wise error-rate significance threshold determined from a randomization approach. In the case of SNPs/indels, blue points denote markers used in the original analysis, whereas red points denote markers that had originally been pruned but whose $p$-values were computed after signal discovery. Panel (a) shows gene models over the plotting interval. Exons are represented by rectangles whereas introns are represented by horizontal lines. Coding sequences are shown in blue and the direction of transcription is indicated by arrows.

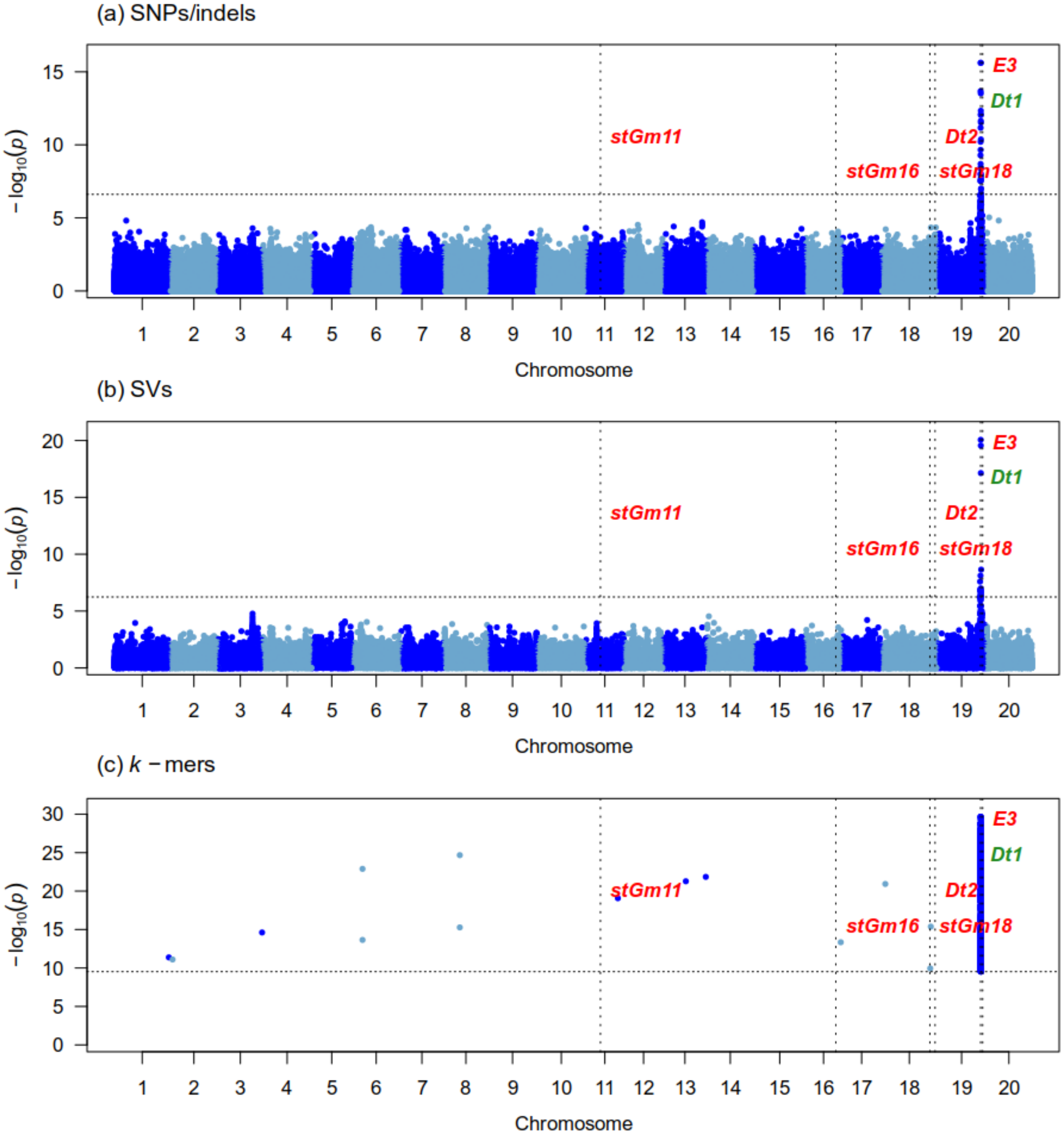


Figure S17: Manhattan plots generated from the GWAS analysis of stem termination type (first GWAS) on 349 samples using three genotype datasets : (a) Platypus (SNPs and indels), (b) Paragraph (SVs), (c) $k$-mers presence/absence. The x-axis shows the position along the reference assembly version 4 of Williams82. Each point shows the $-\log_{10}(p)$ associated with a particular marker or $k$-mer. Horizontal dotted lines indicate the 5% family-wise error-rate significance threshold determined from a randomization approach. Vertical dotted lines indicate the position of signals associated with the trait. Documented loci are colored according to whether they were found (green) or not (red) by a particular approach. The "Gm" prefix has been left out of chromosome names for simplicity.

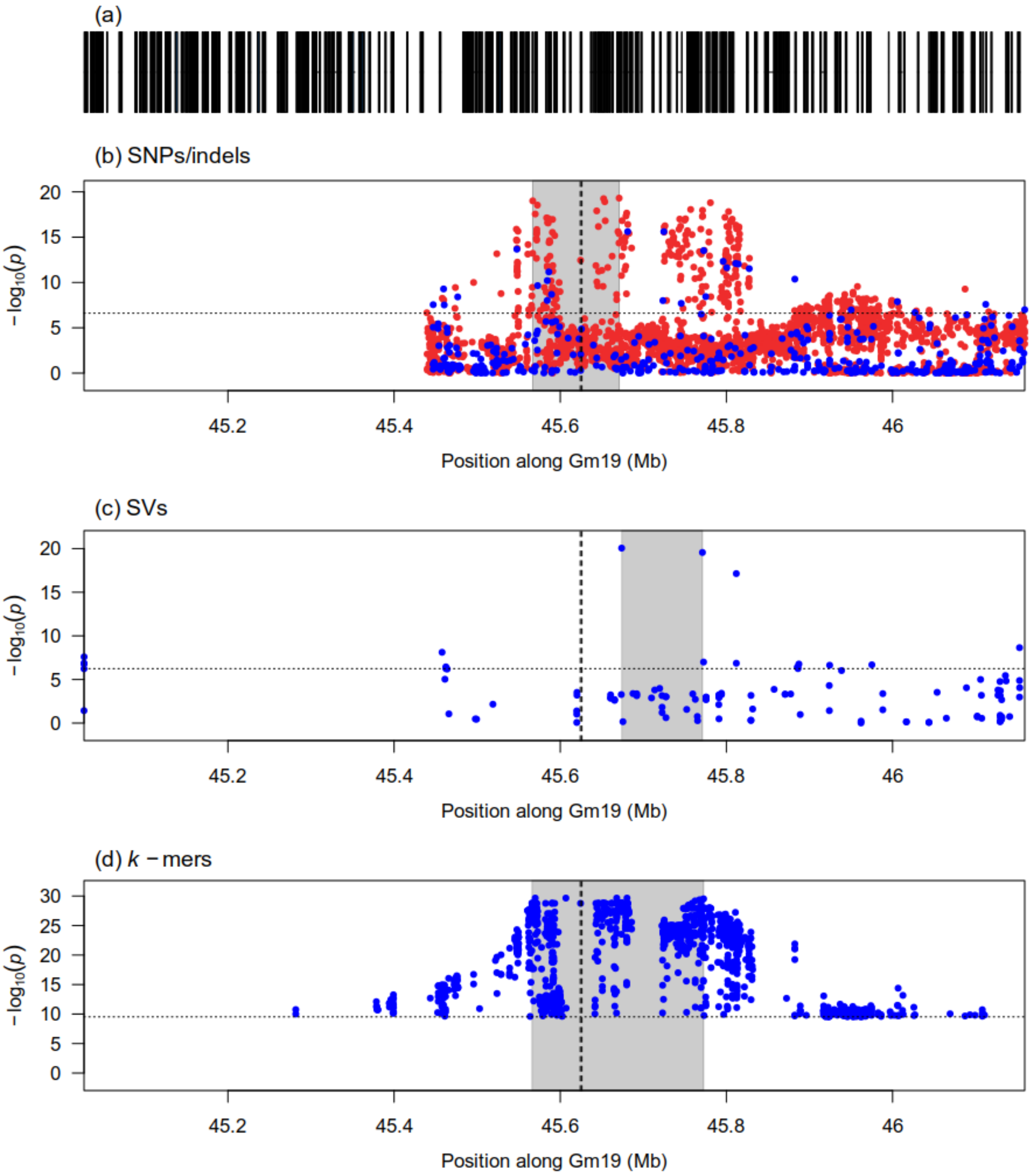


Figure S18: Zoomed-in Manhattan plots of signals detected by the GWAS analysis of stem termination type (first GWAS) at the *Dt1* locus using three genotype datasets: (b) Platypus (SNPs and indels), (c) Paragraph (SVs), (d) $k$-mers presence/absence. Panel (a) shows gene models over the genomic interval. Horizontal dotted lines indicate the 5% family-wise error-rate significance threshold determined from a randomization approach. Vertical dotted lines indicate the location of the Glyma.19g194300 gene associated with the locus. Gray shaded rectangles indicate the region delimited by the top 5% (SVs) or top 1% (SNPs/indels and $k$-mers) associations in the signal region. In the case of SNPs/indels, blue points denote markers used in the original analysis, whereas red points denote markers that had originally been pruned but whose $p$-values were computed after signal discovery.

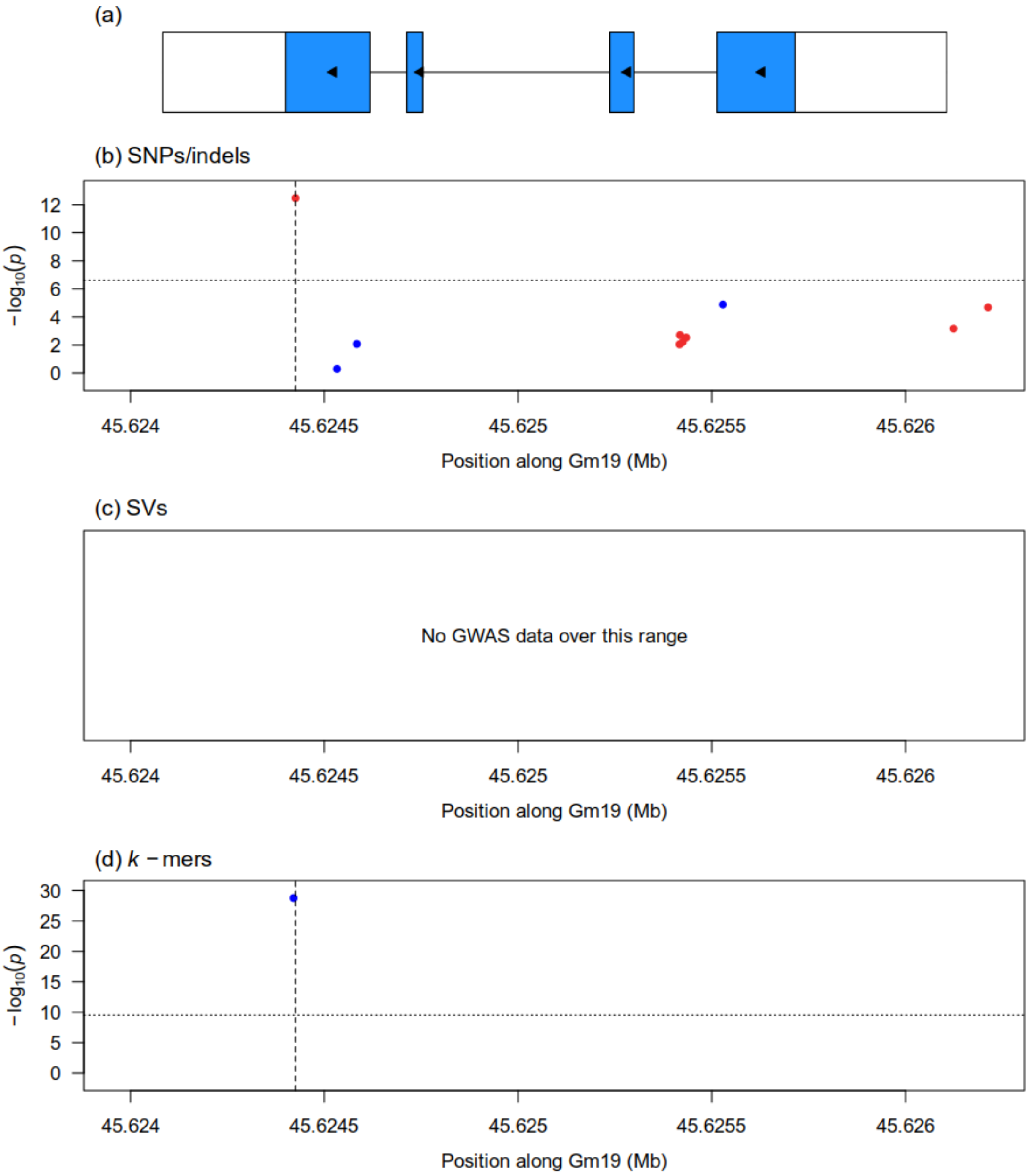


Figure S19: Zoomed-in Manhattan plots generated from the GWAS analysis of stem termination type (first GWAS) at the Glyma.19g194300 gene associated with the *Dt1* locus using three genotype datasets: (b) Platypus (SNPs and indels), (c) Paragraph (SVs), (d) $k$-mers presence/absence. Vertical dotted lines in panels (b) and (d) indicate the location of a causal SNP at this locus. Horizontal dotted lines indicate the 5% family-wise error-rate significance threshold determined from a randomization approach. In the case of SNPs/indels, blue points denote markers used in the original analysis, whereas red points denote markers that had originally been pruned but whose $p$-values were computed after signal discovery. Panel (a) shows gene models over the plotting interval. Exons are represented by rectangles whereas introns are represented by horizontal lines. Coding sequences are shown in blue and the direction of transcription is indicated by arrows.

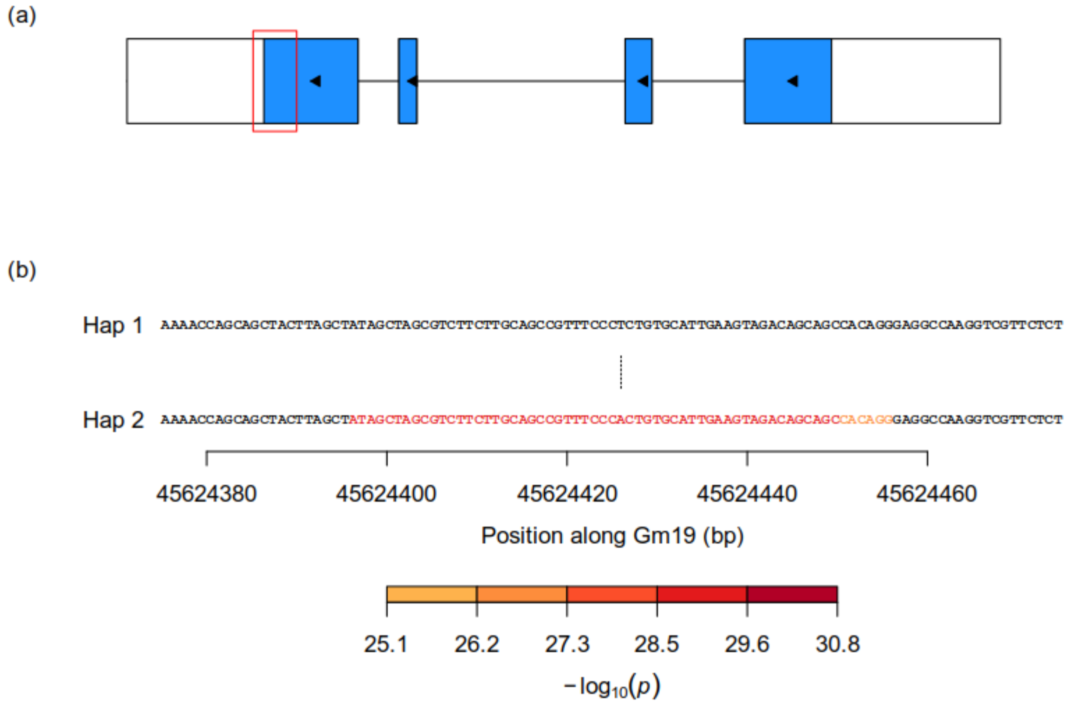


(c)

| | Hap 1 | Hap 2 |
|---|---|---|
| Determinate | 37 | 52 |
| Indeterminate | 231 | 1 |
| Semi-determinate | 23 | 5 |

Figure S20: Identification of a causal SNP underlying significant $k$-mers at the Glyma.19g194300 gene associated with the *Dt1* locus for stem termination type (first GWAS). (a) Gene model of Glyma.19g194300. Exons are represented by rectangles whereas introns are represented by horizontal lines. Coding sequences are shown in blue and the direction of transcription is indicated by arrows. The red rectangle identifies the region that is zoomed-in in panel (b). (b) Nucleotide sequences of haplotypes observed in at least five samples across the dataset. Individual nucleotides are colored according to the $-\log_{10}(p)$ of the most significant $k$-mer overlapping them. Dashes indicate gaps in haplotype sequence alignment whereas vertical lines indicate differences in sequence between two haplotypes. (c) Contingency table of the phenotypes and haplotypes observed in the dataset. Haplotypes correspond to those shown in panel (b).

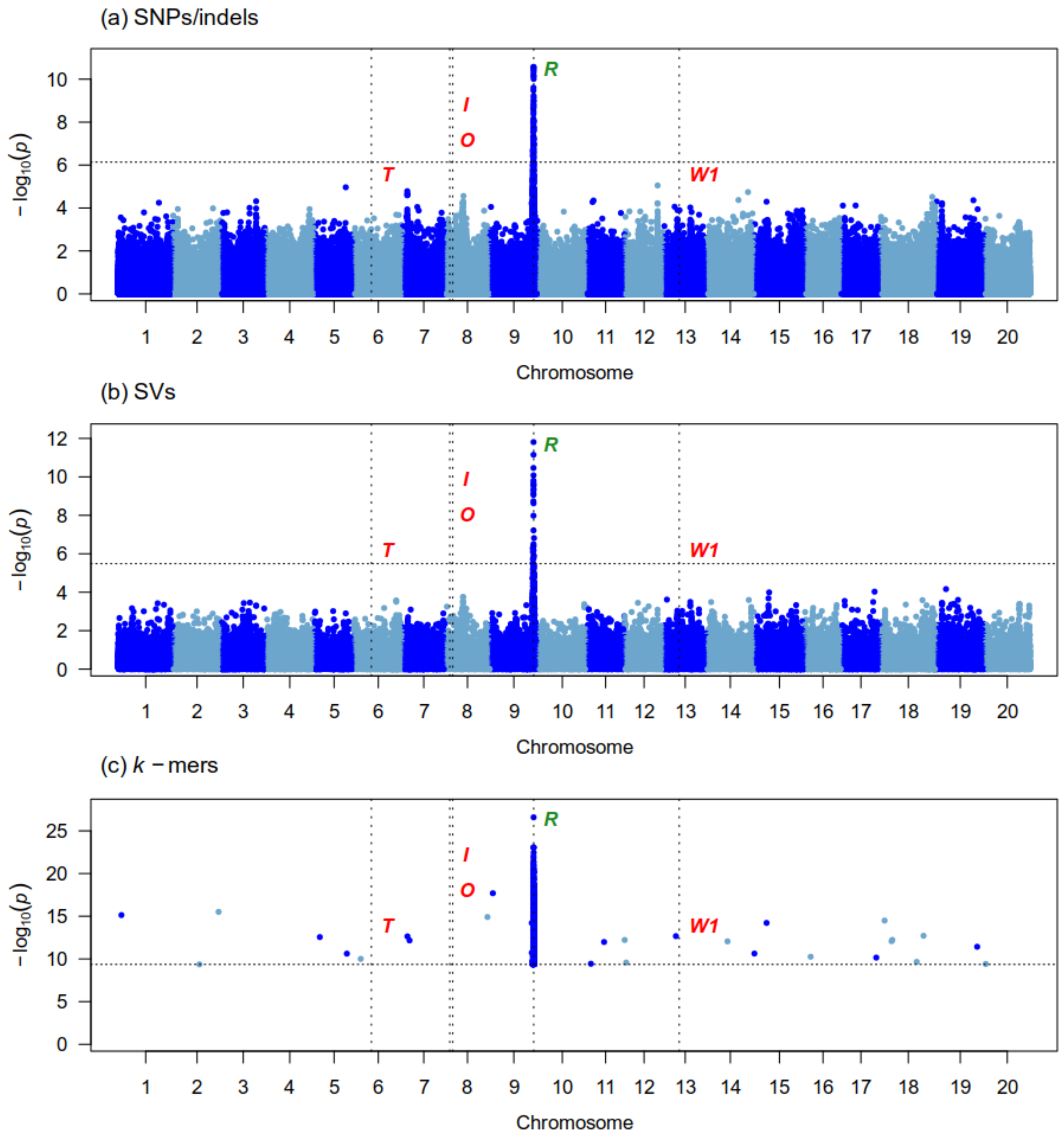


Figure S21: Manhattan plots generated from the GWAS analysis of hilum color (second GWAS) on 160 samples using three genotype datasets : (a) Platypus (SNPs and indels), (b) Paragraph (SVs), (c) $k$-mers presence/absence. The x-axis shows the position along the reference assembly version 4 of Williams82. Each point shows the $-\log_{10}(p)$ associated with a particular marker or $k$-mer. Horizontal dotted lines indicate the 5% family-wise error-rate significance threshold determined from a randomization approach. Vertical dotted lines indicate the position of signals associated with the trait. Documented loci are colored according to whether they were found (green) or not (red) by a particular approach. The "Gm" prefix has been left out of chromosome names for simplicity.

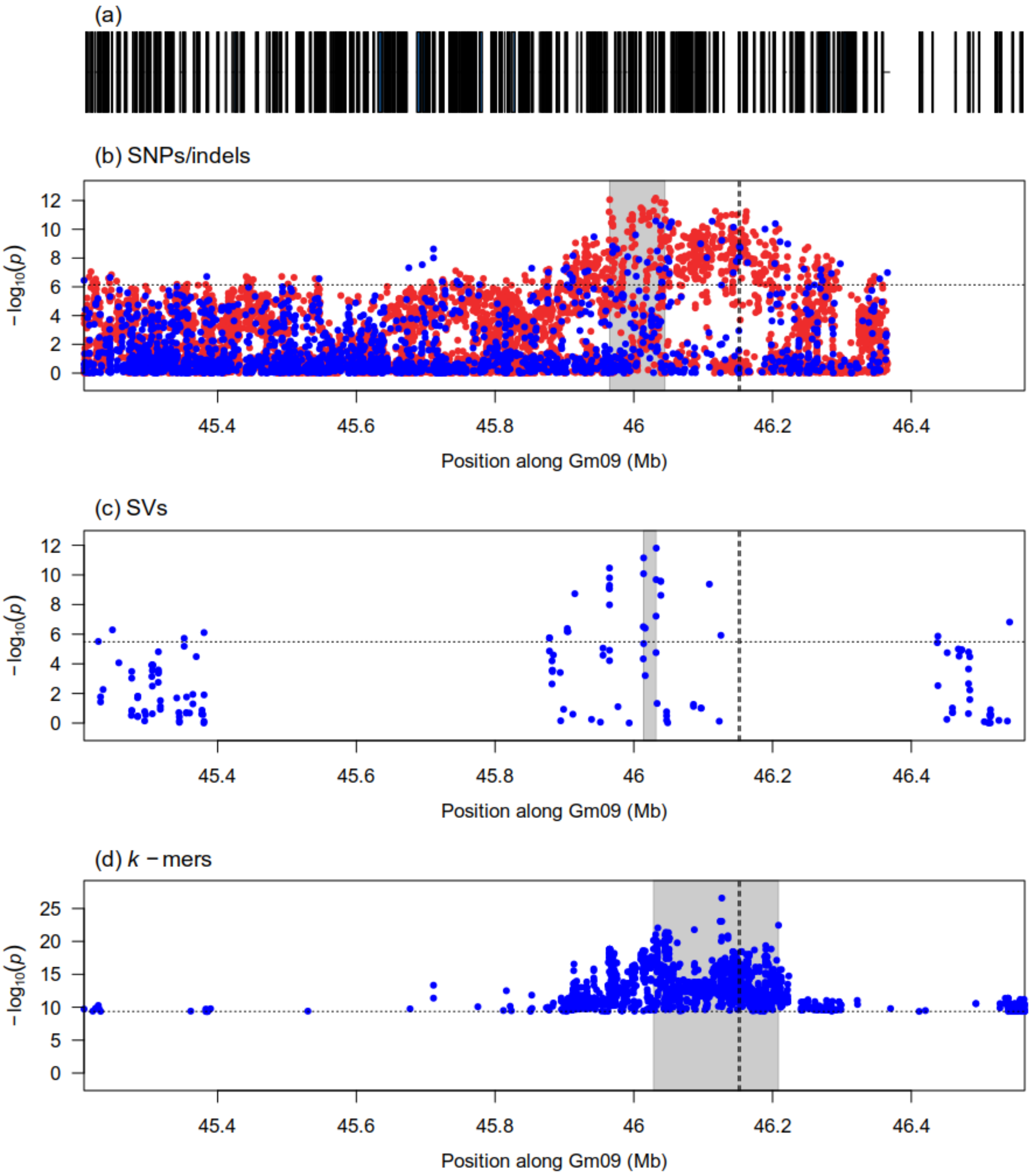


Figure S22: Zoomed-in Manhattan plots of signals detected by the GWAS analysis of hilum color (second GWAS) at the $R$ locus using three genotype datasets: (b) Platypus (SNPs and indels), (c) Paragraph (SVs), (d) $k$-mers presence/absence. Panel (a) shows gene models over the genomic interval. Horizontal dotted lines indicate the 5% family-wise error-rate significance threshold determined from a randomization approach. Vertical dotted lines indicate the location of the Glyma.09g235100 gene associated with the locus. Gray shaded rectangles indicate the region delimited by the top 5% (SVs) or top 1% (SNPs/indels and $k$-mers) associations in the signal region. In the case of SNPs/indels, blue points denote markers used in the original analysis, whereas red points denote markers that had originally been pruned but whose $p$-values were computed after signal discovery.

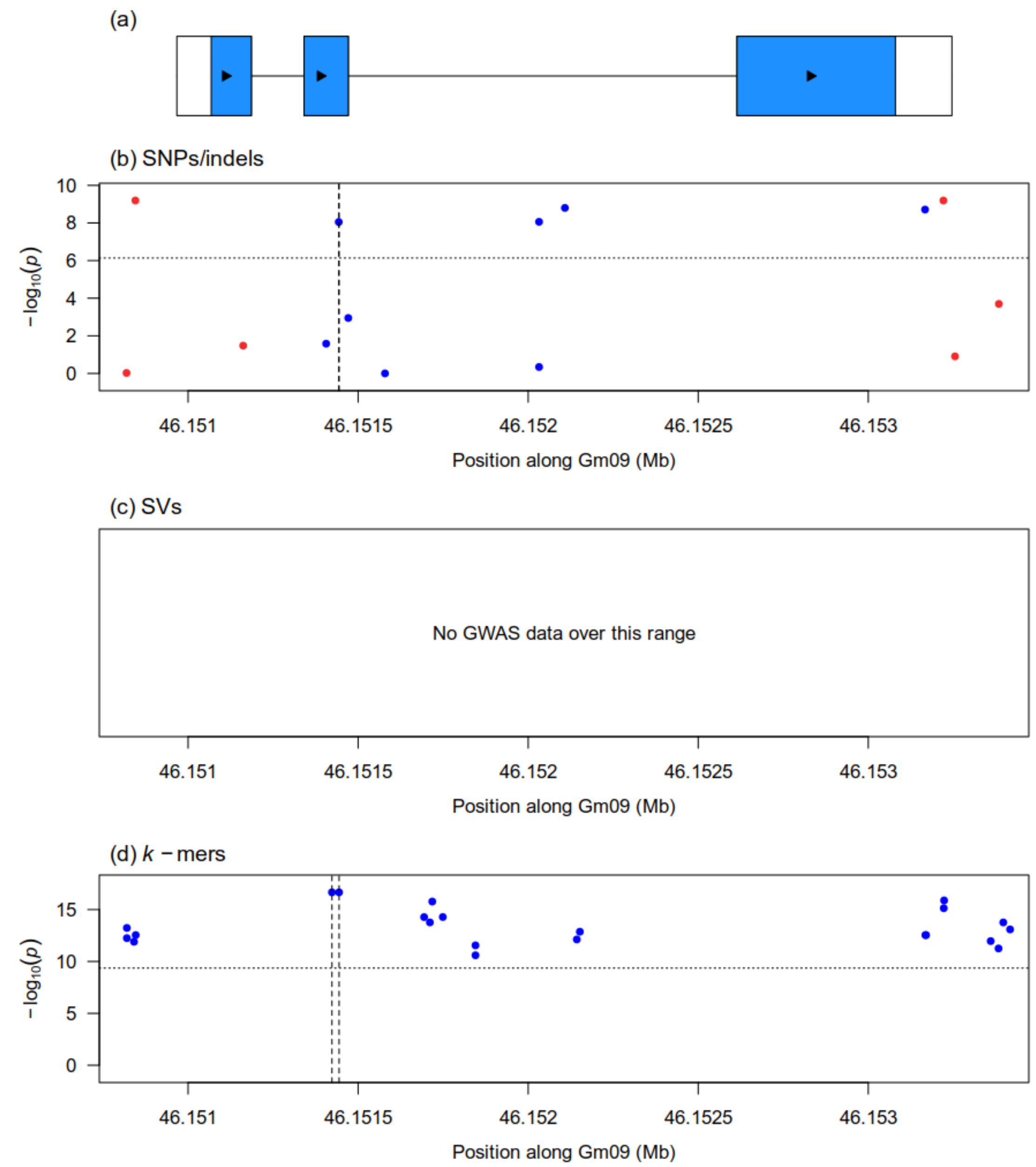


Figure S23: Zoomed-in Manhattan plots generated from the GWAS analysis of hilum color (second GWAS) at the Glyma.09g235100 gene associated with the $R$ locus using three genotype datasets: (b) Platypus (SNPs and indels), (c) Paragraph (SVs), (d) $k$-mers presence/absence. Vertical dotted lines indicate the location an indel (panels b and d) and a SNP (panel d) that are putatively causal at this locus. Horizontal dotted lines indicate the 5% family-wise error-rate significance threshold determined from a randomization approach. In the case of SNPs/indels, blue points denote markers used in the original analysis, whereas red points denote markers that had originally been pruned but whose $p$-values were computed after signal discovery. Panel (a) shows gene models over the plotting interval. Exons are represented by rectangles whereas introns are represented by horizontal lines. Coding sequences are shown in blue and the direction of transcription is indicated by arrows.

(a)

(b)

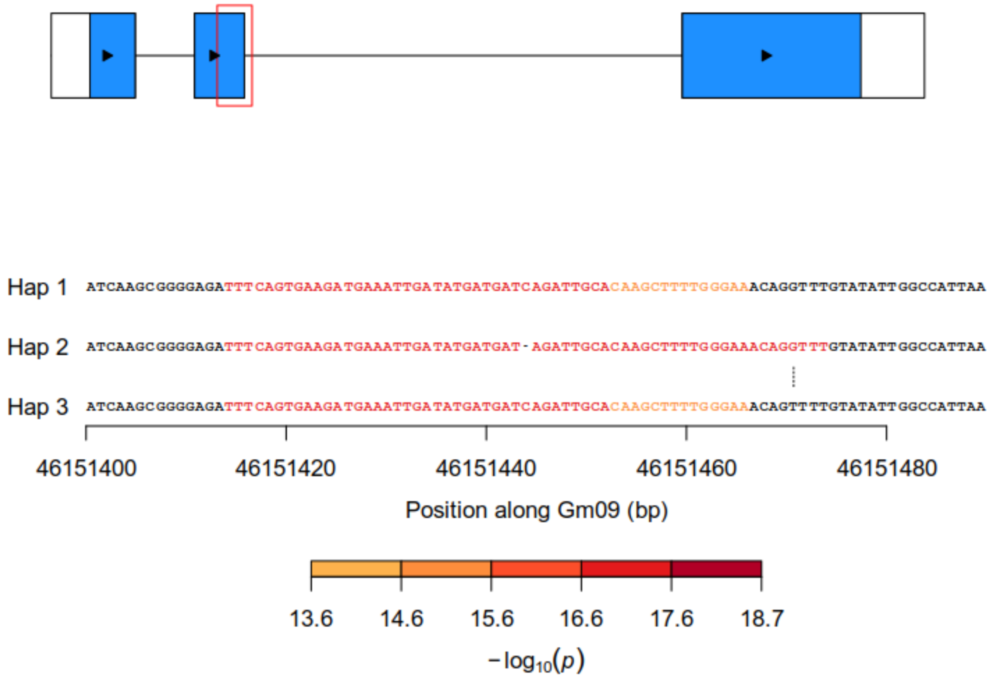


(c)

| | Hap 1 | Hap 2 | Hap 3 |
|---|---|---|---|
| Black | 69 | 0 | 0 |
| Brown | 14 | 18 | 14 |

Figure S24: Identification of a SNP and an indel that are putatively causal underlying significant $k$-mers at the Glyma.09g235100 gene associated with the $R$ locus for hilum color (second GWAS). (a) Gene model of Glyma.09g235100. Exons are represented by rectangles whereas introns are represented by horizontal lines. Coding sequences are shown in blue and the direction of transcription is indicated by arrows. The red rectangle identifies the region that is zoomed-in in panel (b). (b) Nucleotide sequences of haplotypes observed in at least five samples across the dataset. Individual nucleotides are colored according to the $-\log_{10}(p)$ of the most significant $k$-mer overlapping them. Dashes indicate gaps in haplotype sequence alignment whereas vertical lines indicate differences in sequence between two haplotypes. (c) Contingency table of the phenotypes and haplotypes observed in the dataset. Haplotypes correspond to those shown in panel (b).

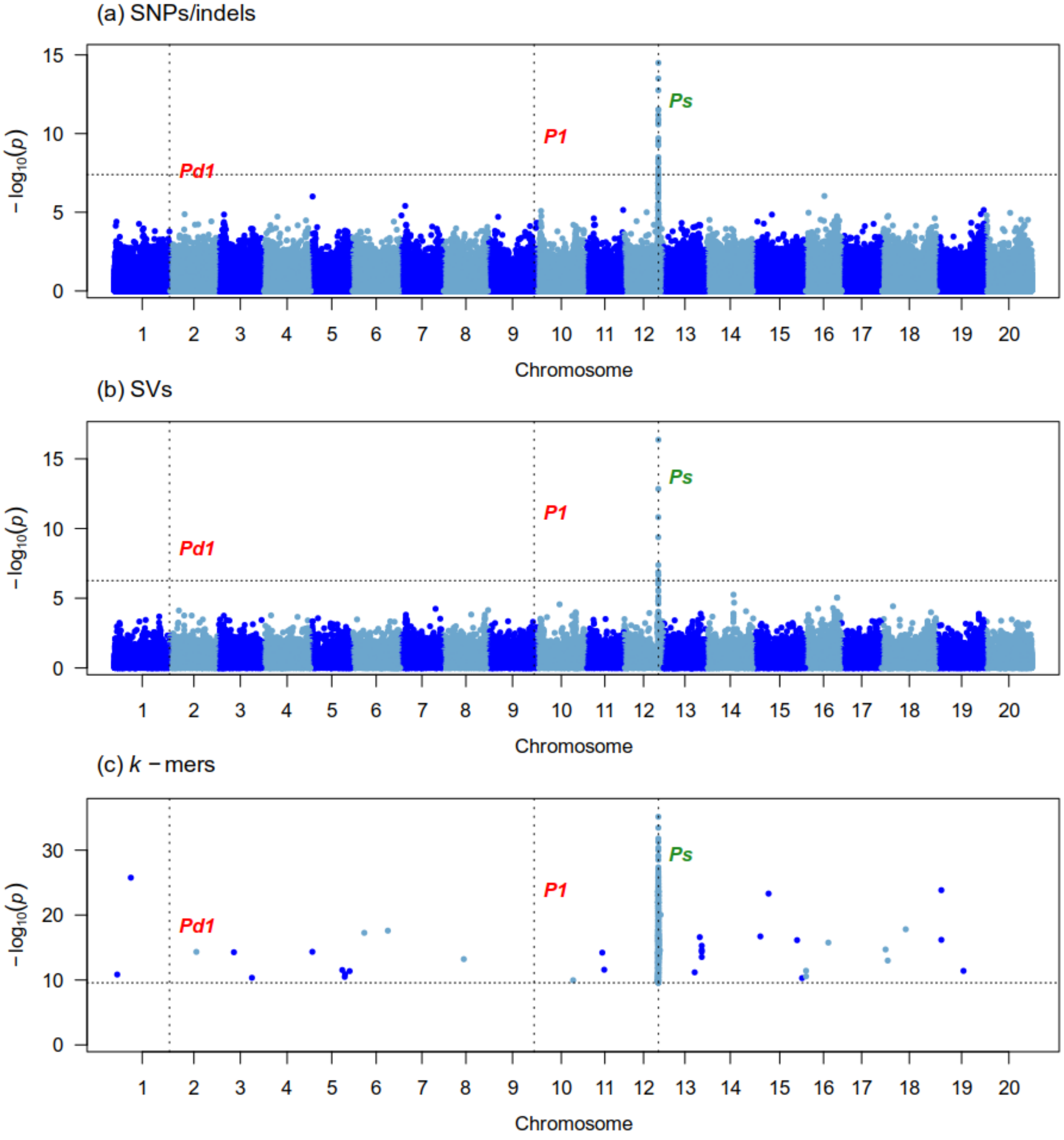


Figure S25: Manhattan plots generated from the GWAS analysis of pubescence density on 353 samples using three genotype datasets : (a) Platypus (SNPs and indels), (b) Paragraph (SVs), (c) $k$-mers presence/absence. The x-axis shows the position along the reference assembly version 4 of Williams82. Each point shows the $-\log_{10}(p)$ associated with a particular marker or $k$-mer. Horizontal dotted lines indicate the 5% family-wise error-rate significance threshold determined from a randomization approach. Vertical dotted lines indicate the position of signals associated with the trait. Documented loci are colored according to whether they were found (green) or not (red) by a particular approach. The "Gm" prefix has been left out of chromosome names for simplicity.

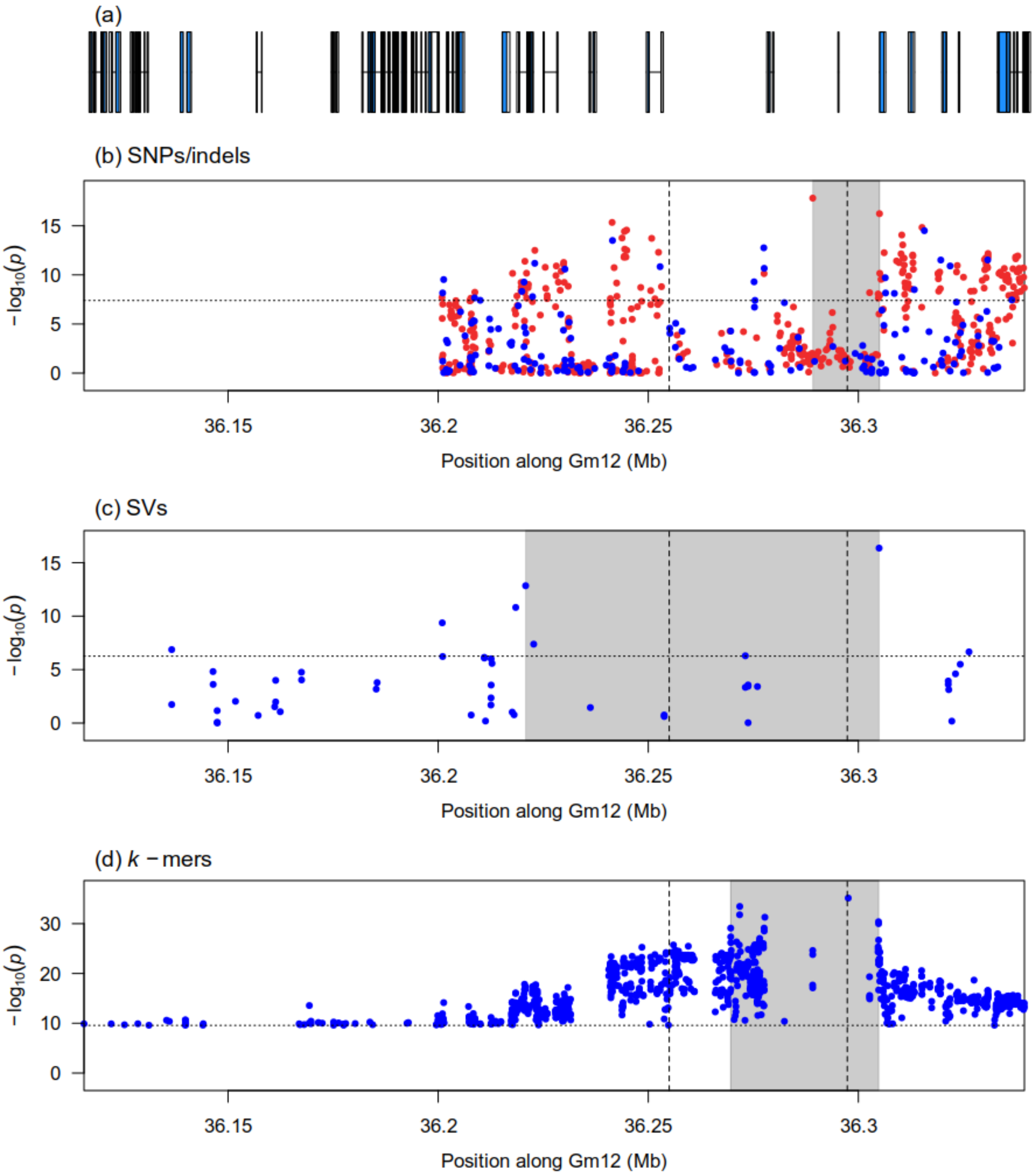


Figure S26: Zoomed-in Manhattan plots of signals detected by the GWAS analysis of pubescence density at the *Ps* locus using three genotype datasets: (b) Platypus (SNPs and indels), (c) Paragraph (SVs), (d) $k$-mers presence/absence. Panel (a) shows gene models over the genomic interval. Horizontal dotted lines indicate the 5% family-wise error-rate significance threshold determined from a randomization approach. Vertical dotted lines indicate the boundaries of the causal CNV overlapping the Glyma.12g187200 gene associated with the locus. Gray shaded rectangles indicate the region delimited by the top 5% (SVs) or top 1% (SNPs/indels and $k$-mers) associations in the signal region. In the case of SNPs/indels, blue points denote markers used in the original analysis, whereas red points denote markers that had originally been pruned but whose $p$-values were computed after signal discovery.

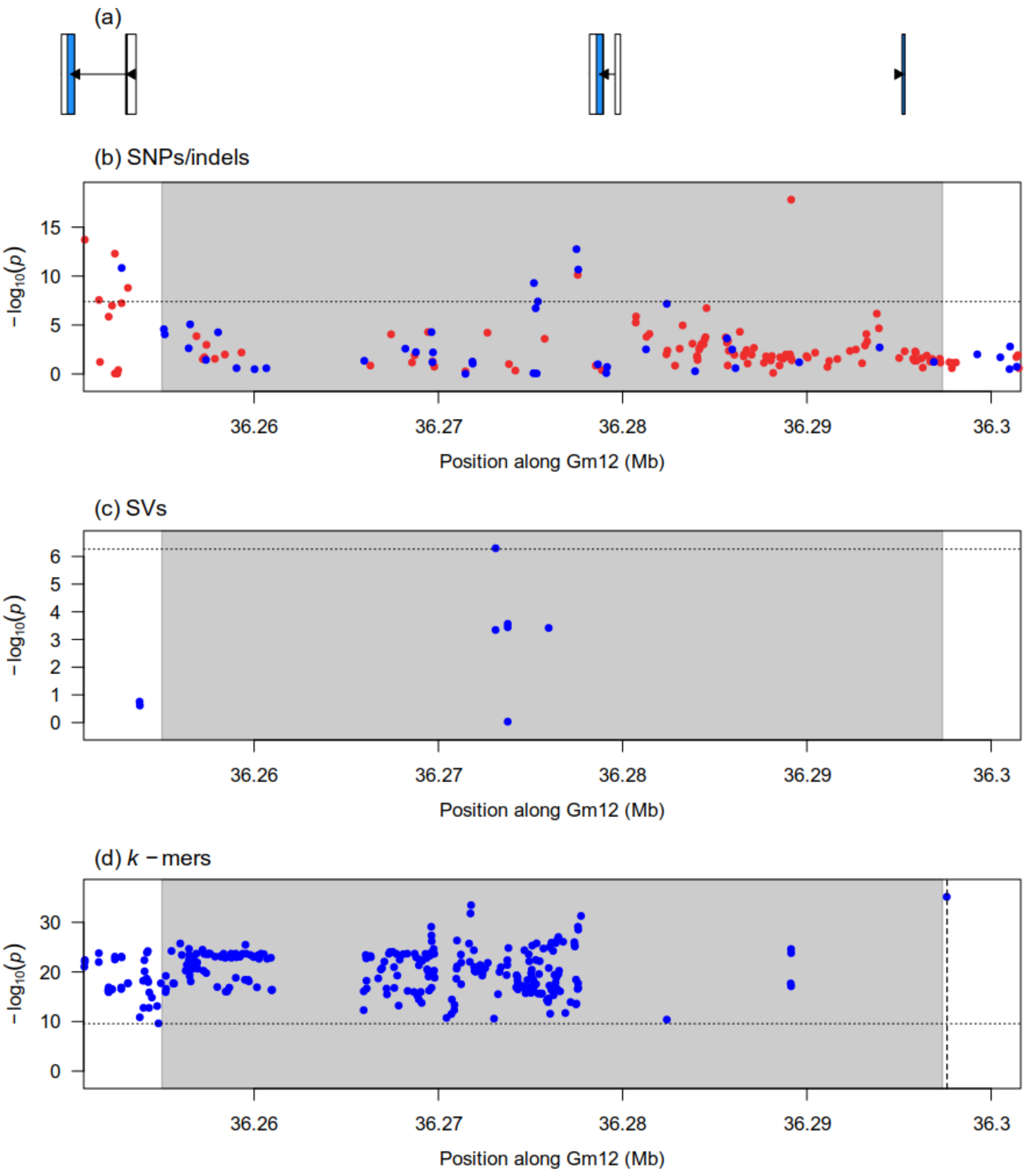


Figure S27: Zoomed-in Manhattan plots generated from the GWAS analysis of pubescence density at the location of the causal CNV overlapping the Glyma.12g187200 gene. associated with the *Ps* locus using three genotype datasets: (b) Platypus (SNPs and indels), (c) Paragraph (SVs), (d) $k$-mers presence/absence. The vertical dotted line in panel (d) indicates the location of the most significant $k$-mer, which is associated with the causal CNV at this locus. The shaded gray rectangles show the boundaries of the causal CNV. Horizontal dotted lines indicate the 5% family-wise error-rate significance threshold determined from a randomization approach. In the case of SNPs/indels, blue points denote markers used in the original analysis, whereas red points denote markers that had originally been pruned but whose $p$-values were computed after signal discovery. Panel (a) shows gene models over the plotting interval. Exons are represented by rectangles whereas introns are represented by horizontal lines. Coding sequences are shown in blue and the direction of transcription is indicated by arrows.

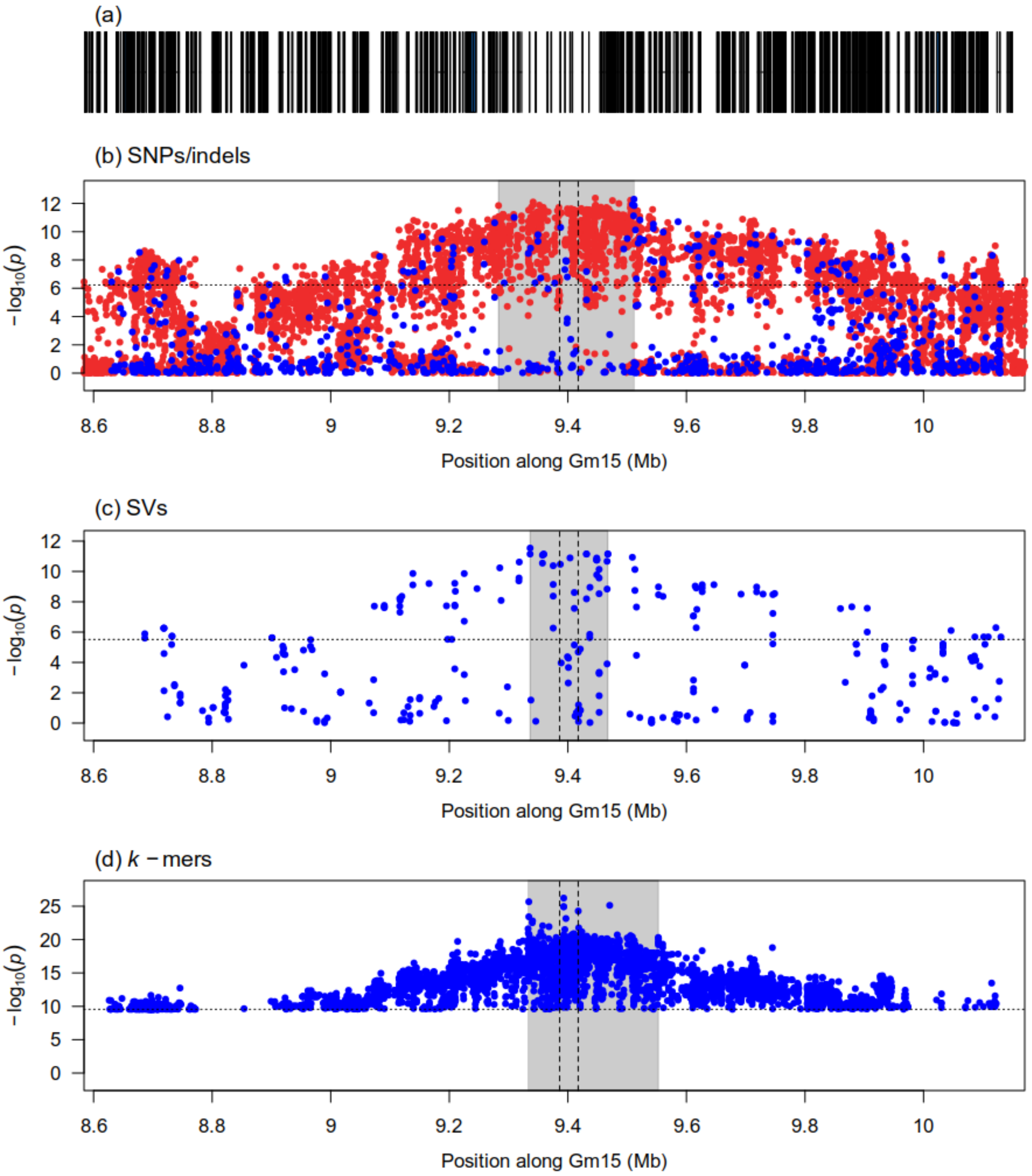


Figure S28: Zoomed-in Manhattan plots of signals detected by the GWAS analysis of seed coat luster (third GWAS) at the *B* locus using three genotype datasets: (b) Platypus (SNPs and indels), (c) Paragraph (SVs), (d) *k*-mers presence/absence. Panel (a) shows gene models over the genomic interval. Horizontal dotted lines indicate the 5% family-wise error-rate significance threshold determined from a randomization approach. Vertical dotted lines indicate the boundaries of the causal CNV associated with the locus. Gray shaded rectangles indicate the region delimited by the top 5% (SVs) or top 1% (SNPs/indels and *k*-mers) associations in the signal region. In the case of SNPs/indels, blue points denote markers used in the original analysis, whereas red points denote markers that had originally been pruned but whose *p*-values were computed after signal discovery.

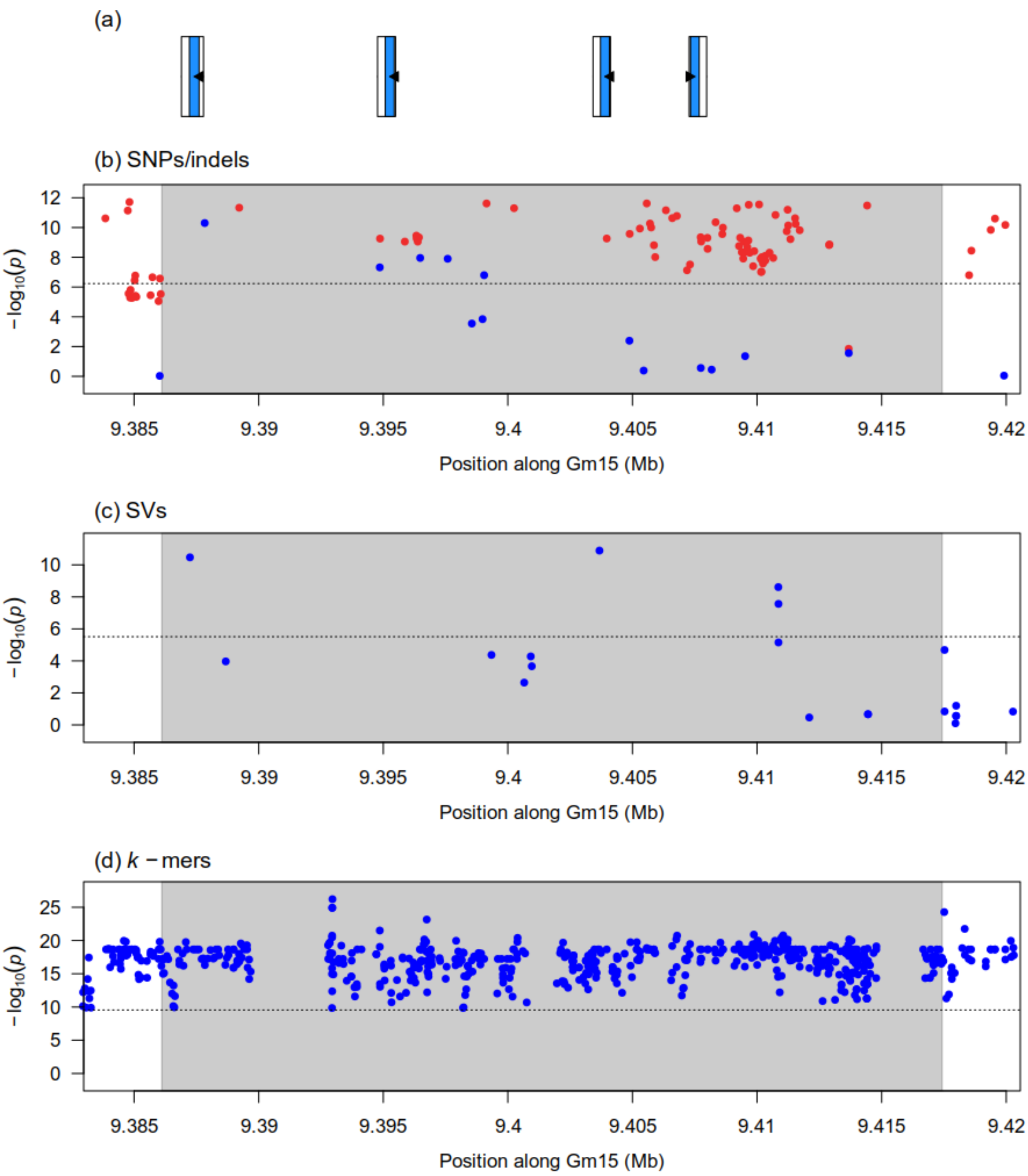


Figure S29: Zoomed-in Manhattan plots generated from the GWAS analysis of seed coat luster (third GWAS) at the location of the causal CNV associated with the *B* locus using three genotype datasets: (b) Platypus (SNPs and indels), (c) Paragraph (SVs), (d) *k*-mers presence/absence. The shaded gray rectangles show the boundaries of the causal CNV. Horizontal dotted lines indicate the 5% family-wise error-rate significance threshold determined from a randomization approach. In the case of SNPs/indels, blue points denote markers used in the original analysis, whereas red points denote markers that had originally been pruned but whose *p*-values were computed after signal discovery. Panel (a) shows gene models over the plotting interval. Exons are represented by rectangles whereas introns are represented by horizontal lines. Coding sequences are shown in blue and the direction of transcription is indicated by arrows.

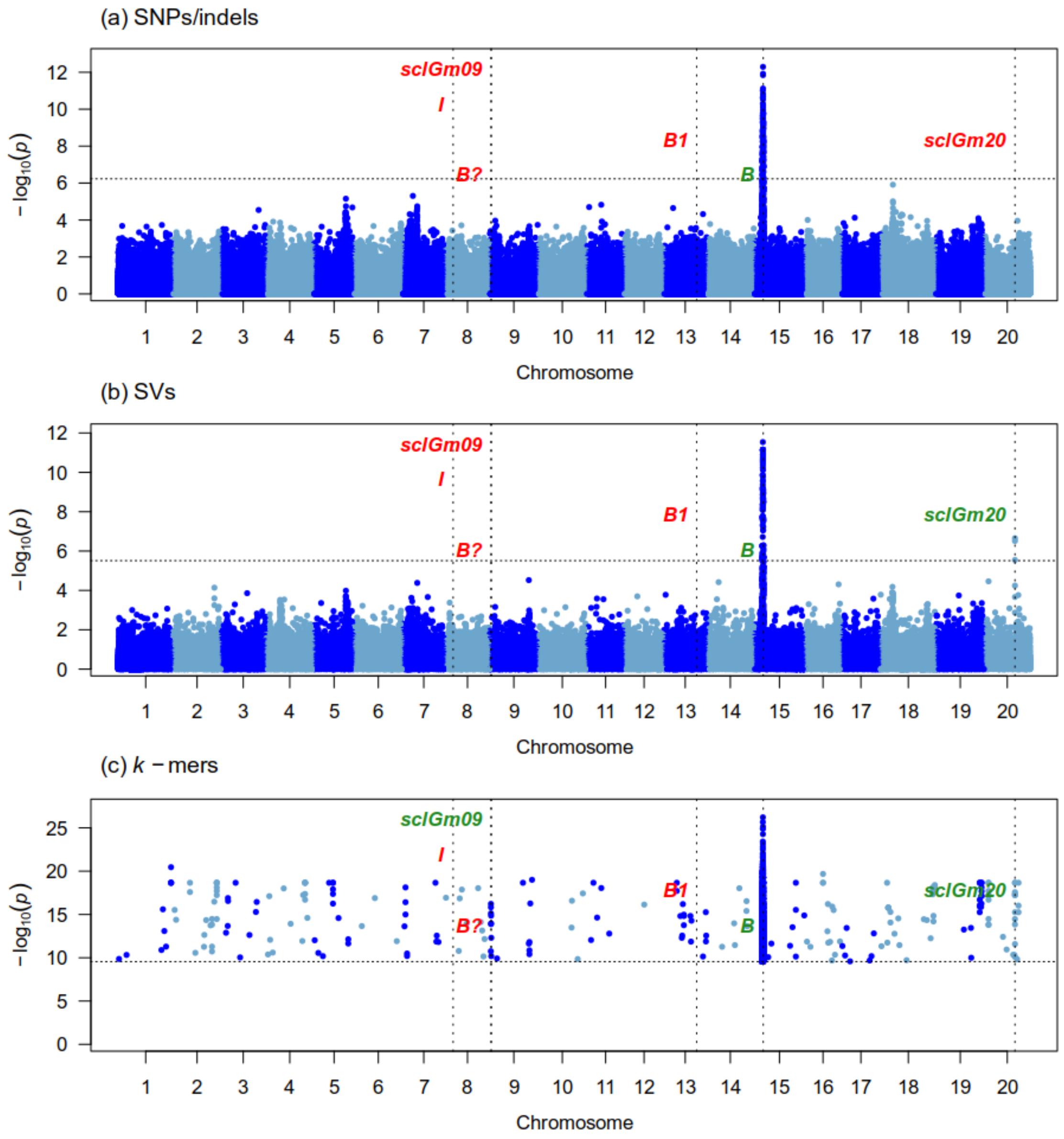


Figure S30: Manhattan plots generated from the GWAS analysis of seed coat luster (third GWAS) on 151 samples using three genotype datasets : (a) Platypus (SNPs and indels), (b) Paragraph (SVs), (c) *k*-mers presence/absence. The x-axis shows the position along the reference assembly version 4 of Williams82. Each point shows the $-\log_{10}(p)$ associated with a particular marker or *k*-mer. Horizontal dotted lines indicate the 5% family-wise error-rate significance threshold determined from a randomization approach. Vertical dotted lines indicate the position of signals associated with the trait. Documented loci are colored according to whether they were found (green) or not (red) by a particular approach. The "Gm" prefix has been left out of chromosome names for simplicity.

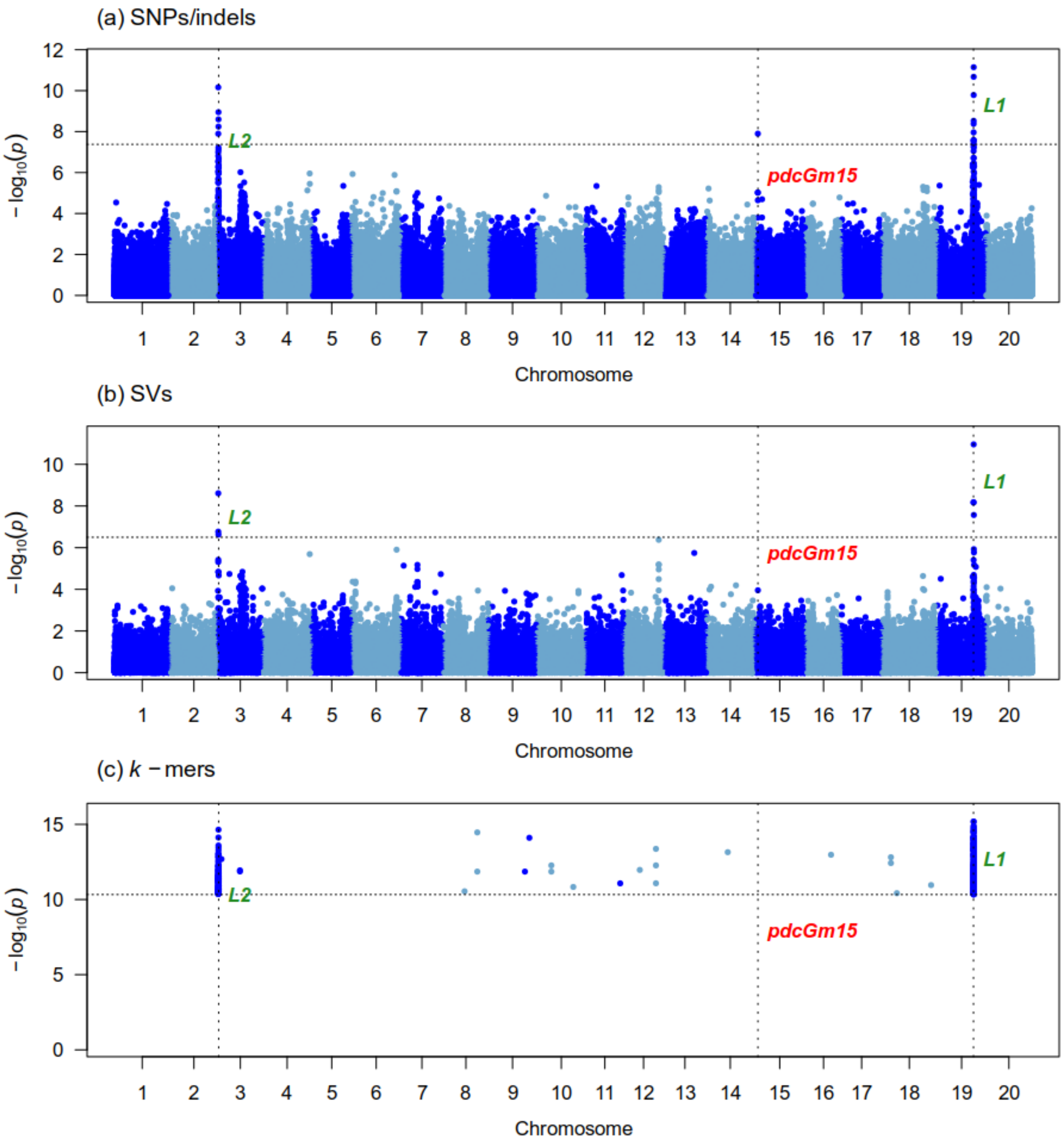


Figure S31: Manhattan plots generated from the GWAS analysis of pod color (first GWAS) on 355 samples using three genotype datasets : (a) Platypus (SNPs and indels), (b) Paragraph (SVs), (c) $k$-mers presence/absence. The x-axis shows the position along the reference assembly version 4 of Williams82. Each point shows the $-\log_{10}(p)$ associated with a particular marker or $k$-mer. Horizontal dotted lines indicate the 5% family-wise error-rate significance threshold determined from a randomization approach. Vertical dotted lines indicate the position of signals associated with the trait. Documented loci are colored according to whether they were found (green) or not (red) by a particular approach. The "Gm" prefix has been left out of chromosome names for simplicity.

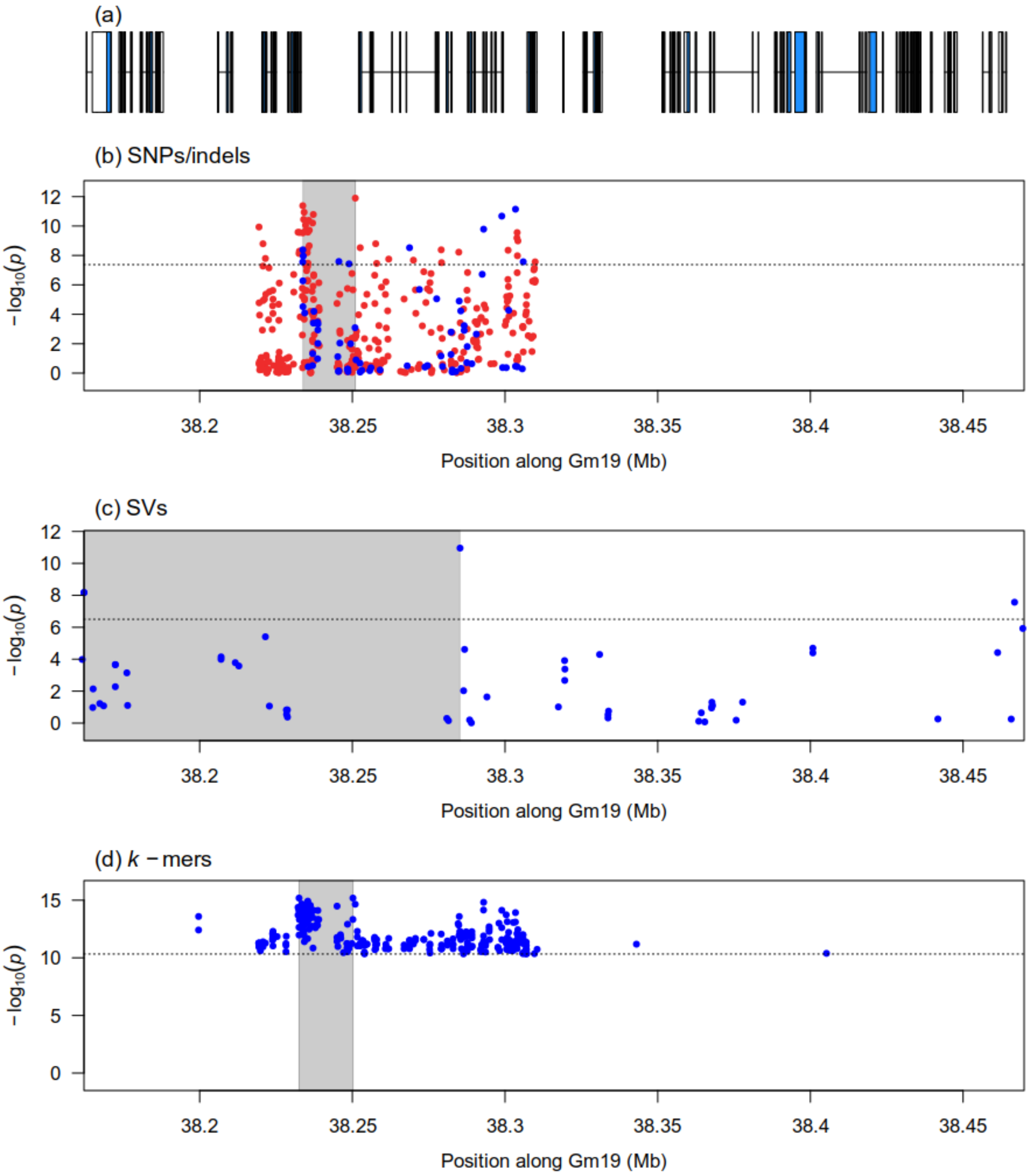


Figure S32: Zoomed-in Manhattan plots of signals detected by the GWAS analysis of pod color (first GWAS) at the *L1* locus using three genotype datasets: (b) Platypus (SNPs and indels), (c) Paragraph (SVs), (d) $k$-mers presence/absence. Panel (a) shows gene models over the genomic interval. Horizontal dotted lines indicate the 5% family-wise error-rate significance threshold determined from a randomization approach. Gray shaded rectangles indicate the region delimited by the top 5% (SVs) or top 1% (SNPs/indels and $k$-mers) associations in the signal region. In the case of SNPs/indels, blue points denote markers used in the original analysis, whereas red points denote markers that had originally been pruned but whose $p$-values were computed after signal discovery.

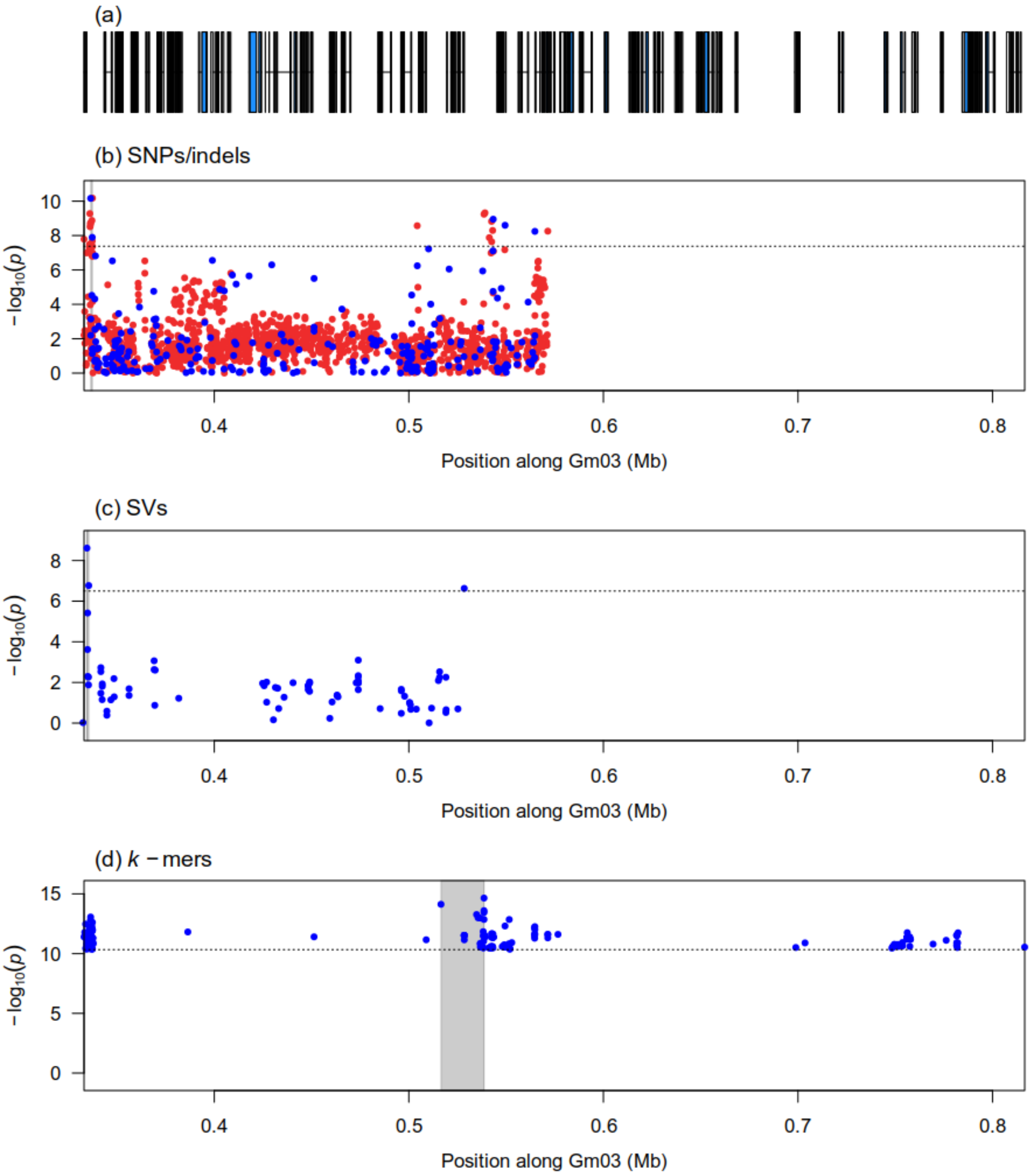


Figure S33: Zoomed-in Manhattan plots of signals detected by the GWAS analysis of pod color (first GWAS) at the *L2* locus using three genotype datasets: (b) Platypus (SNPs and indels), (c) Paragraph (SVs), (d) $k$-mers presence/absence. Panel (a) shows gene models over the genomic interval. Horizontal dotted lines indicate the 5% family-wise error-rate significance threshold determined from a randomization approach. Gray shaded rectangles indicate the region delimited by the top 5% (SVs) or top 1% (SNPs/indels and $k$-mers) associations in the signal region. In the case of SNPs/indels, blue points denote markers used in the original analysis, whereas red points denote markers that had originally been pruned but whose $p$-values were computed after signal discovery.

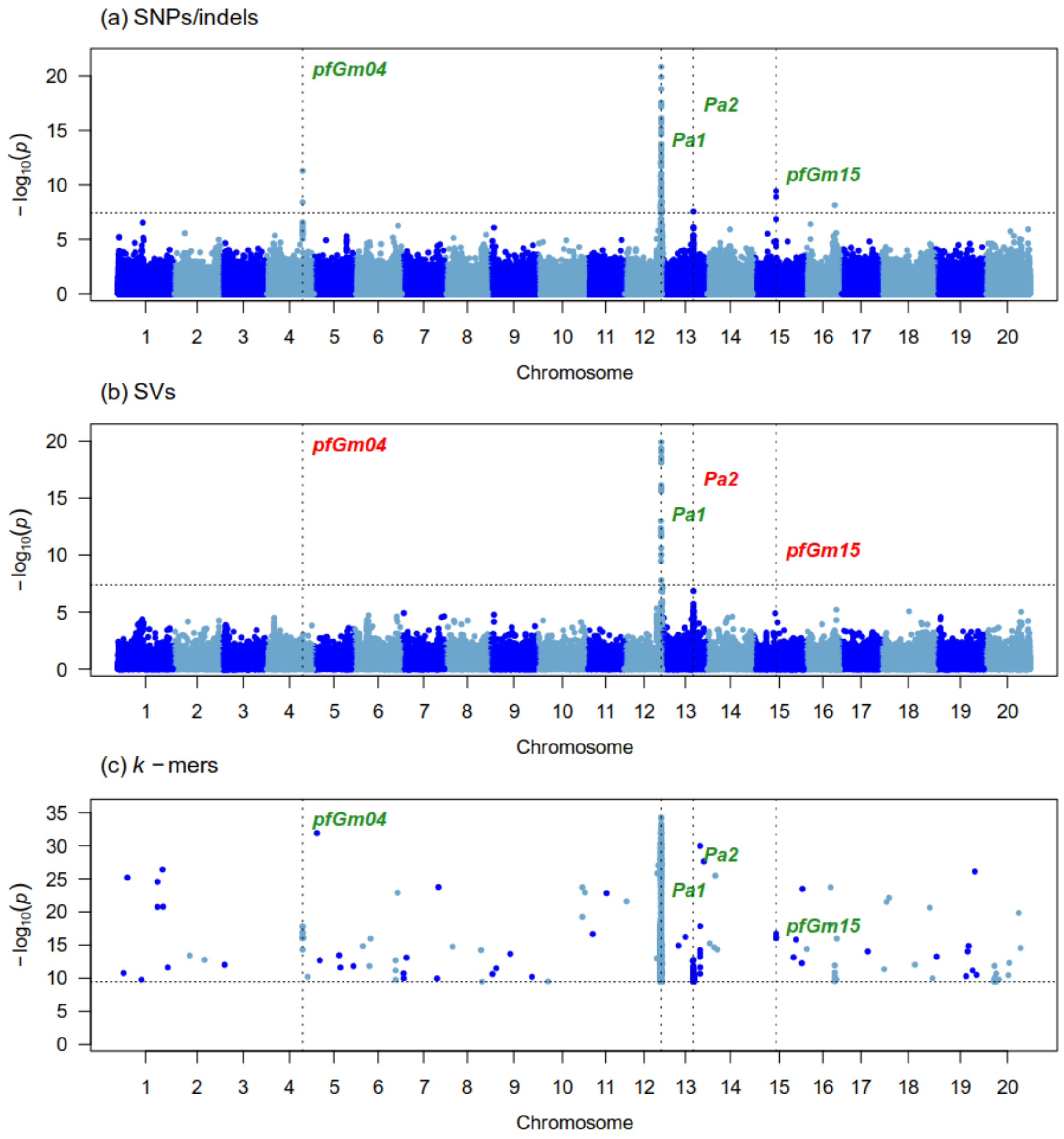


Figure S34: Manhattan plots generated from the GWAS analysis of pubescence form (first GWAS) on 348 samples using three genotype datasets : (a) Platypus (SNPs and indels), (b) Paragraph (SVs), (c) $k$-mers presence/absence. The x-axis shows the position along the reference assembly version 4 of Williams82. Each point shows the $-\log_{10}(p)$ associated with a particular marker or $k$-mer. Horizontal dotted lines indicate the 5% family-wise error-rate significance threshold determined from a randomization approach. Vertical dotted lines indicate the position of signals associated with the trait. Documented loci are colored according to whether they were found (green) or not (red) by a particular approach. The "Gm" prefix has been left out of chromosome names for simplicity.

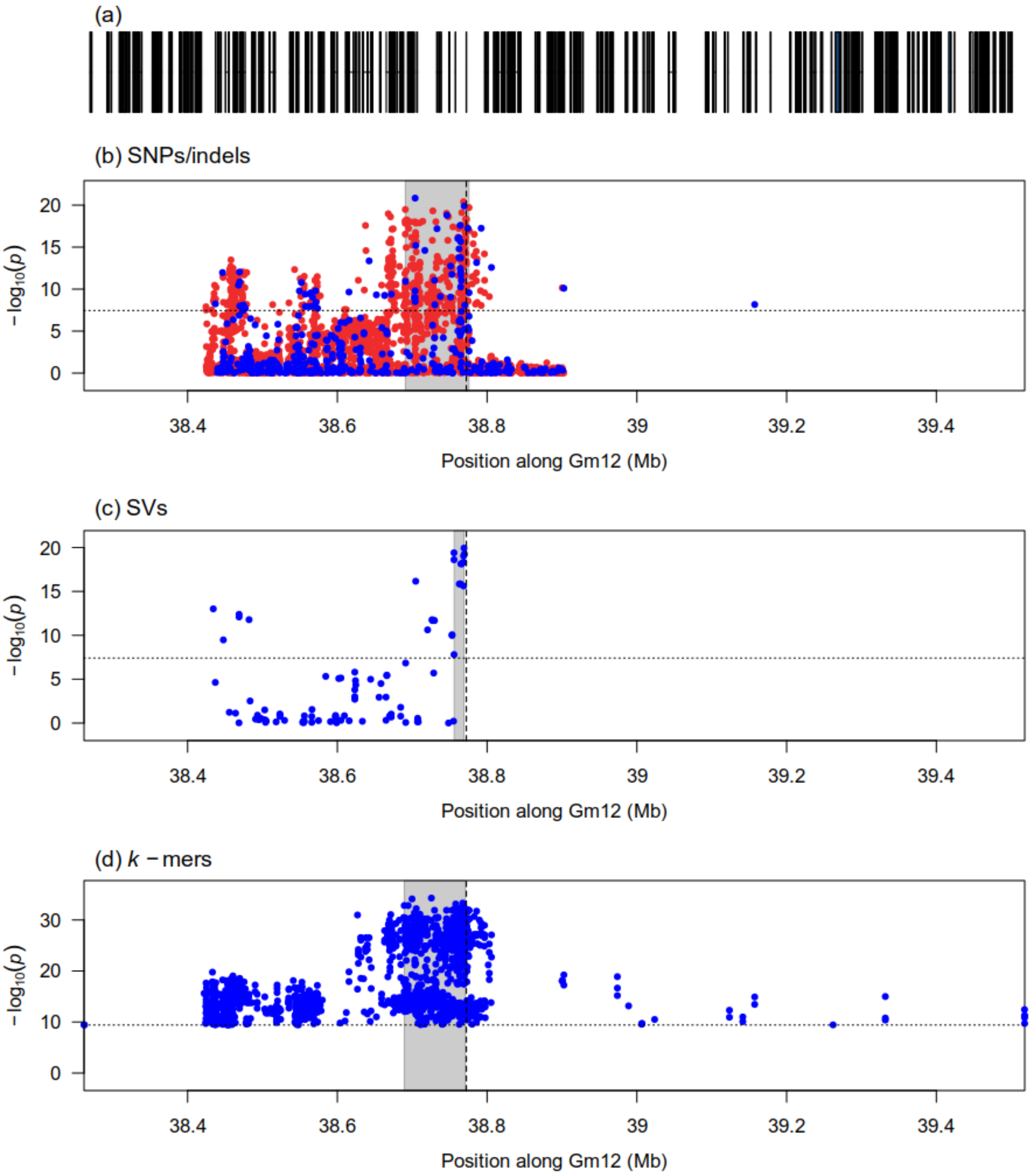


Figure S35: Zoomed-in Manhattan plots of signals detected by the GWAS analysis of pubescence form (first GWAS) at the *Pa1* locus using three genotype datasets: (b) Platypus (SNPs and indels), (c) Paragraph (SVs), (d) $k$-mers presence/absence. Panel (a) shows gene models over the genomic interval. Horizontal dotted lines indicate the 5% family-wise error-rate significance threshold determined from a randomization approach. Vertical dotted lines indicate the location of the Glyma.12g213900 gene that we suggest as a candidate for this locus. Gray shaded rectangles indicate the region delimited by the top 5% (SVs) or top 1% (SNPs/indels and $k$-mers) associations in the signal region. In the case of SNPs/indels, blue points denote markers used in the original analysis, whereas red points denote markers that had originally been pruned but whose $p$-values were computed after signal discovery.

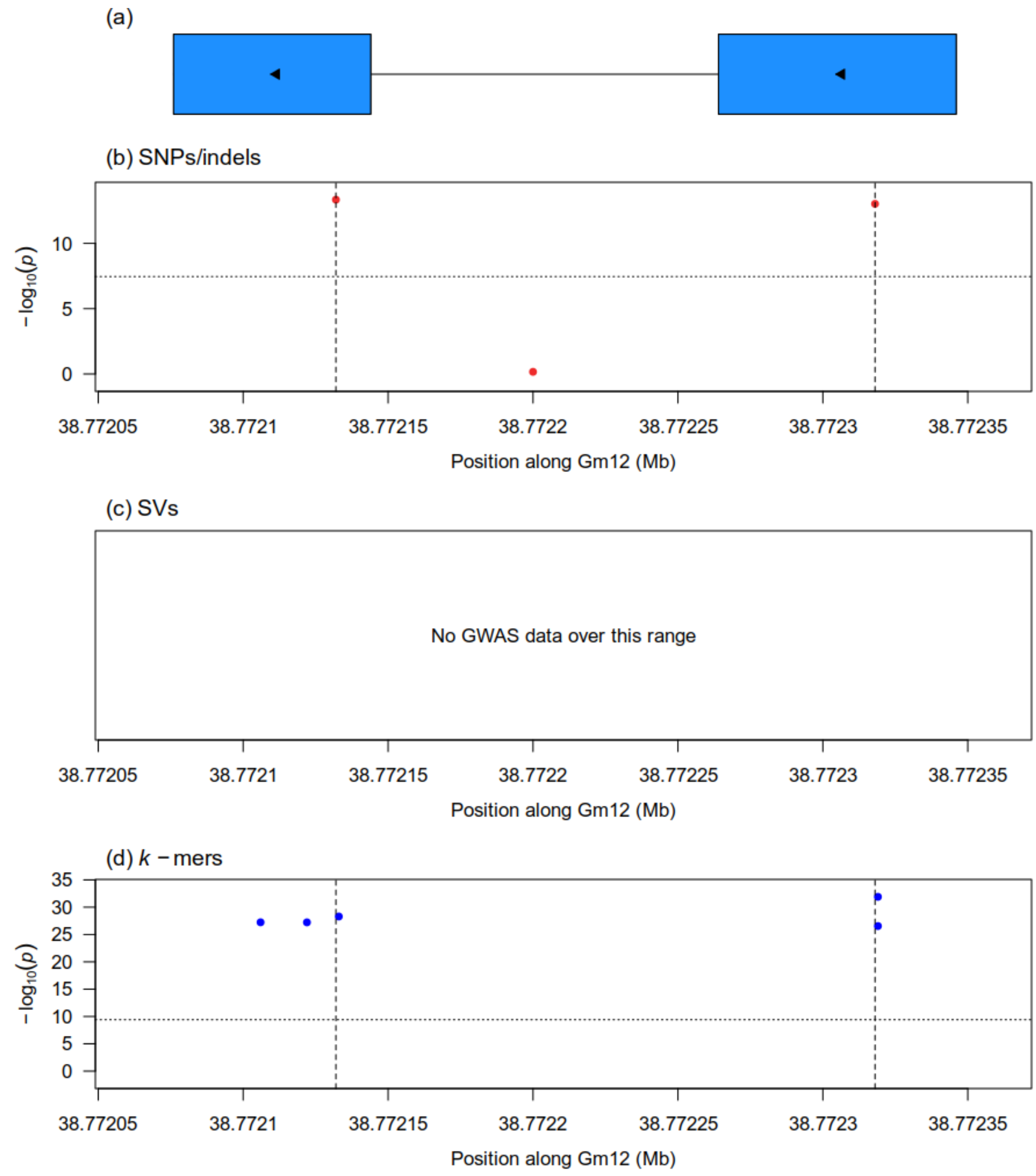


Figure S36: Zoomed-in Manhattan plots generated from the GWAS analysis of pubescence form (first GWAS) at the Glyma.12g213900 candidate gene putatively associated with the *Pa1* locus using three genotype datasets: (b) Platypus (SNPs and indels), (c) Paragraph (SVs), (d) $k$-mers presence/absence. Vertical dotted lines in panels (b) and (d) indicate the location of two highly significant non-synonymous SNPs at this gene. Horizontal dotted lines indicate the 5% family-wise error-rate significance threshold determined from a randomization approach. In the case of SNPs/indels, blue points denote markers used in the original analysis, whereas red points denote markers that had originally been pruned but whose $p$-values were computed after signal discovery. Panel (a) shows gene models over the plotting interval. Exons are represented by rectangles whereas introns are represented by horizontal lines. Coding sequences are shown in blue and the direction of transcription is indicated by arrows.

(a)

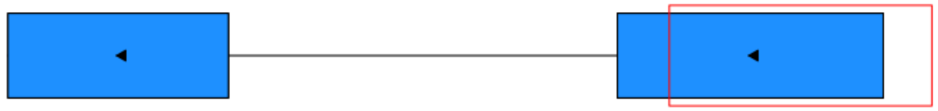

(b)

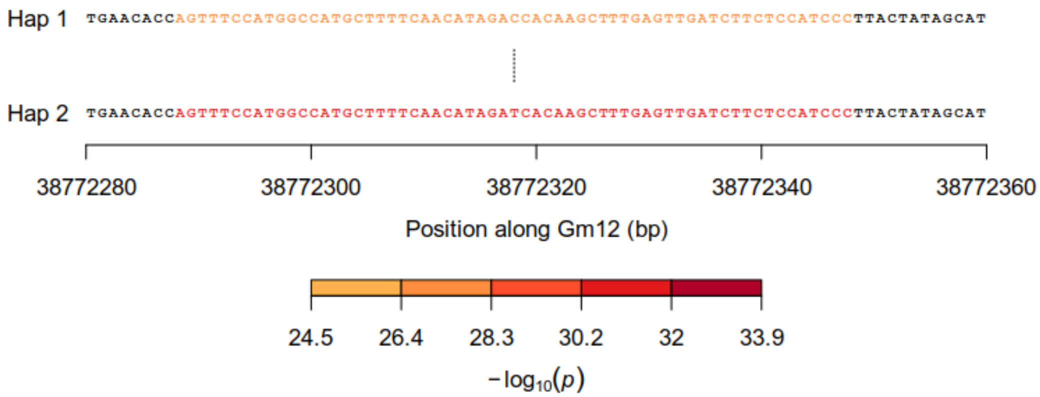


(c)

| | Hap 1 | Hap 2 |
|---|---|---|
| Appressed | 6 | 32 |
| Erect | 256 | 3 |
| Semi-appressed | 42 | 9 |

Figure S37: Identification of a non-synonymous SNP underlying significant $k$-mers at the Glyma.12g213900 candidate gene associated with the *Pa1* locus for pubescence form (first GWAS). (a) Gene model of Glyma.12g213900 candidate. Exons are represented by rectangles whereas introns are represented by horizontal lines. Coding sequences are shown in blue and the direction of transcription is indicated by arrows. The red rectangle identifies the region that is zoomed-in in panel (b). (b) Nucleotide sequences of haplotypes observed in at least five samples across the dataset. Individual nucleotides are colored according to the $-\log_{10}(p)$ of the most significant $k$-mer overlapping them. Dashes indicate gaps in haplotype sequence alignment whereas vertical lines indicate differences in sequence between two haplotypes. (c) Contingency table of the phenotypes and haplotypes observed in the dataset. Haplotypes correspond to those shown in panel (b).

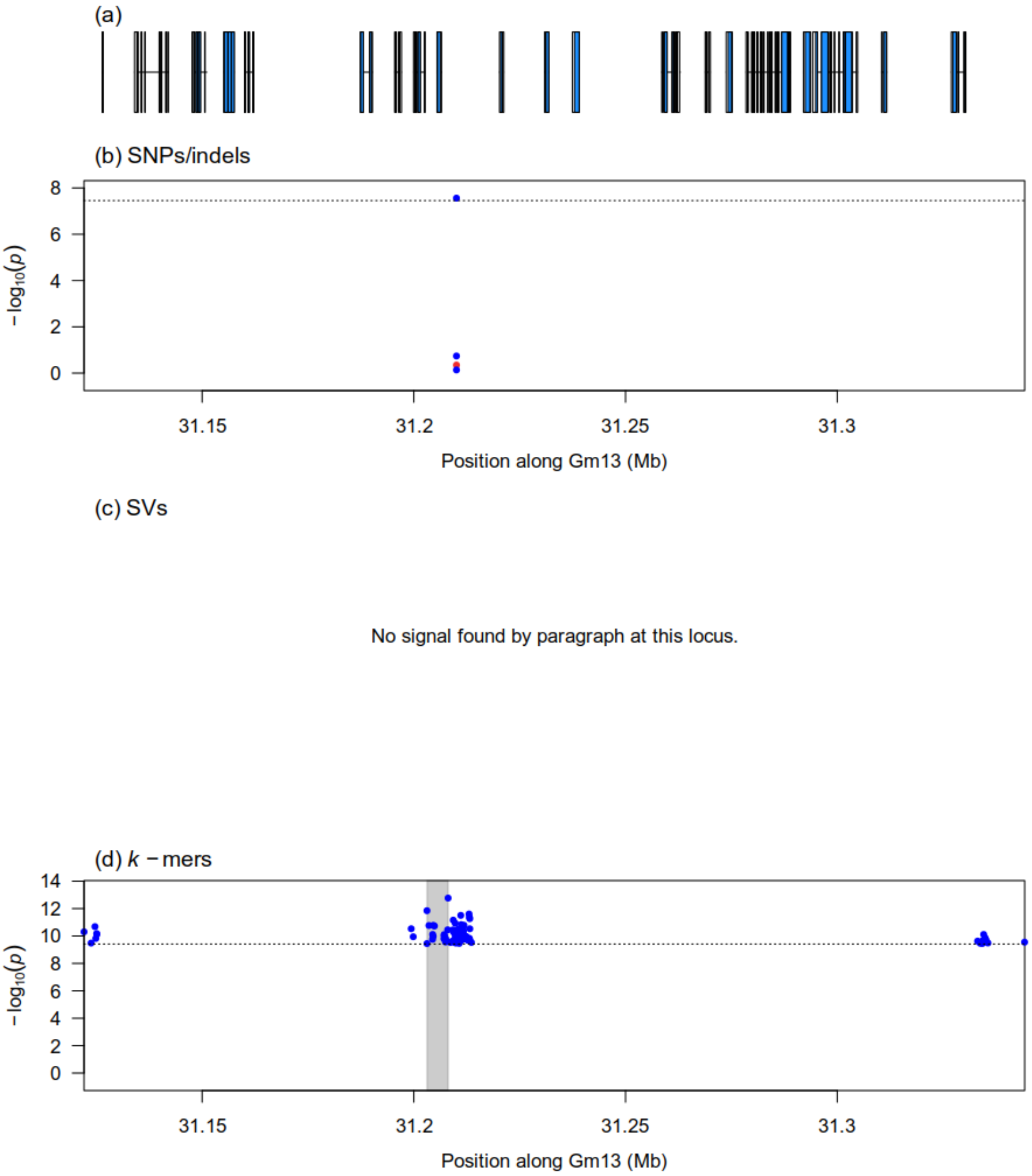


Figure S38: Zoomed-in Manhattan plots of signals detected by the GWAS analysis of pubescence form (first GWAS) at the *Pa2* locus using three genotype datasets: (b) Platypus (SNPs and indels), (c) Paragraph (SVs), (d) $k$-mers presence/absence. Panel (a) shows gene models over the genomic interval. Horizontal dotted lines indicate the 5% family-wise error-rate significance threshold determined from a randomization approach. Gray shaded rectangles indicate the region delimited by the top 5% (SVs) or top 1% (SNPs/indels and $k$-mers) associations in the signal region. In the case of SNPs/indels, blue points denote markers used in the original analysis, whereas red points denote markers that had originally been pruned but whose $p$-values were computed after signal discovery.

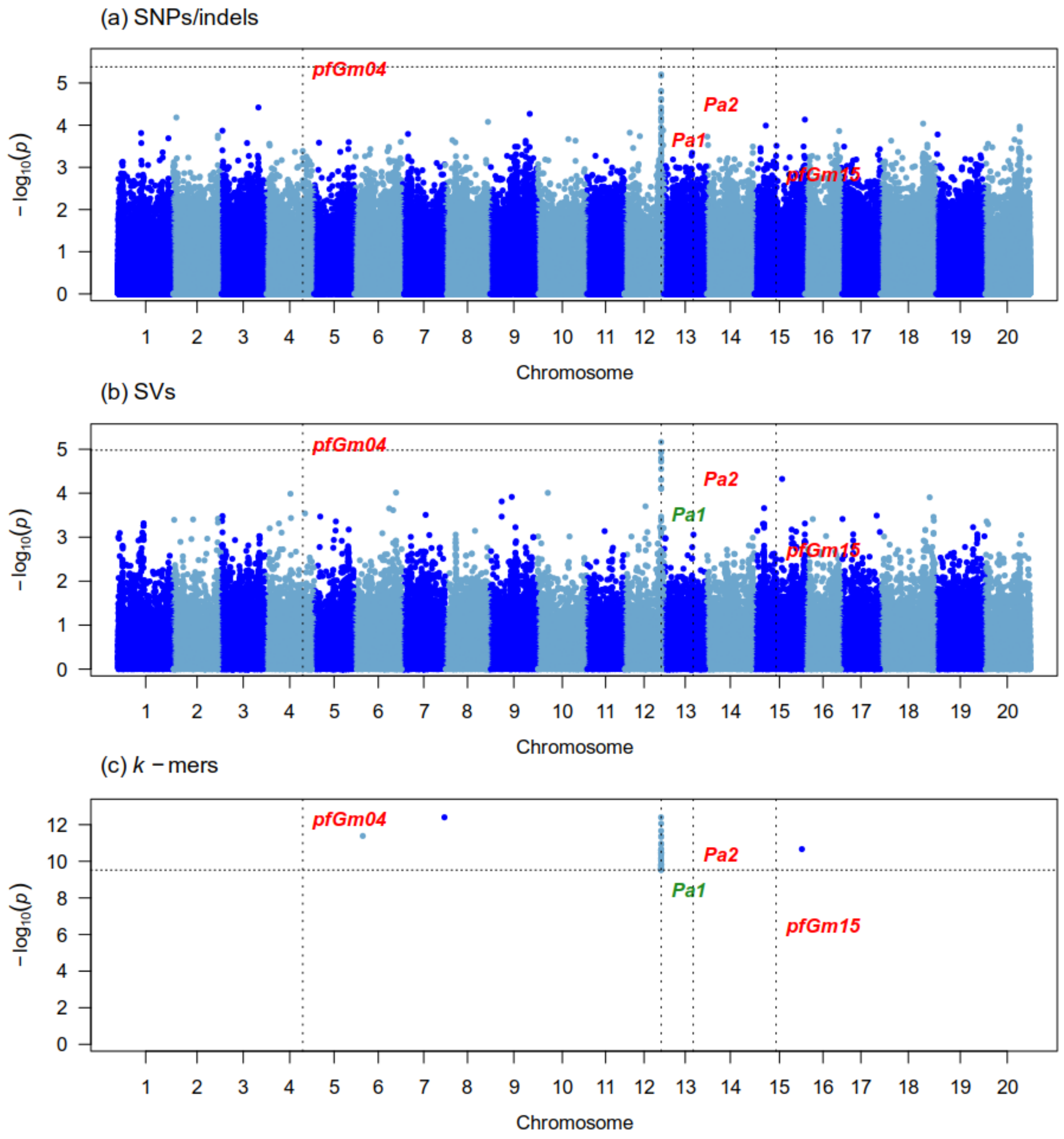


Figure S39: Manhattan plots generated from the GWAS analysis of pubescence form (second GWAS) on 89 samples using three genotype datasets : (a) Platypus (SNPs and indels), (b) Paragraph (SVs), (c) $k$-mers presence/absence. The x-axis shows the position along the reference assembly version 4 of Williams82. Each point shows the $-\log_{10}(p)$ associated with a particular marker or $k$-mer. Horizontal dotted lines indicate the 5% family-wise error-rate significance threshold determined from a randomization approach. Vertical dotted lines indicate the position of signals associated with the trait. Documented loci are colored according to whether they were found (green) or not (red) by a particular approach. The "Gm" prefix has been left out of chromosome names for simplicity.

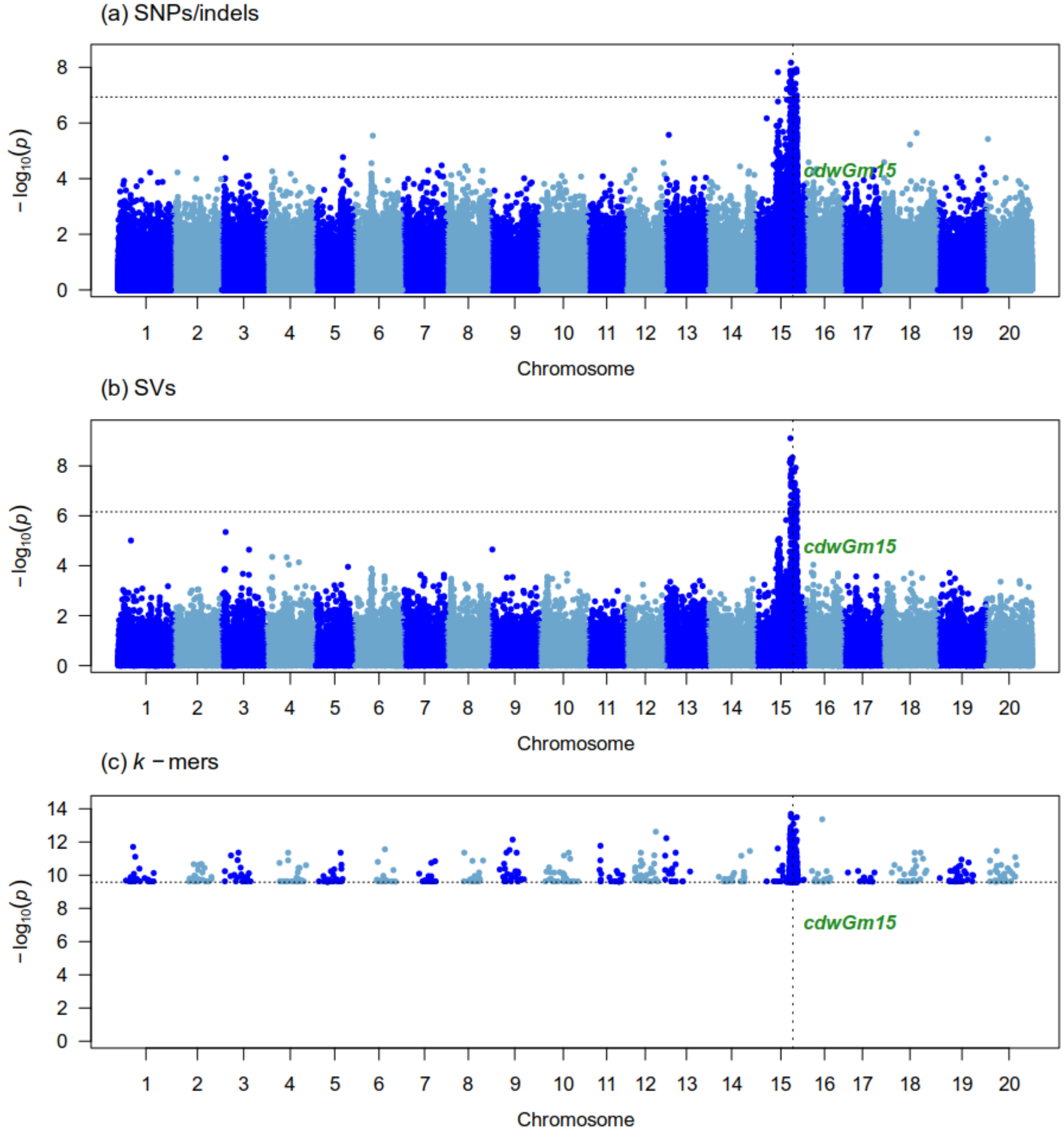


Figure S40: Manhattan plots generated from the GWAS analysis of corrected dry weight (resistance to *P. sojae*) on 362 samples using three genotype datasets : (a) Platypus (SNPs and indels), (b) Paragraph (SVs), (c) $k$-mers presence/absence. The x-axis shows the position along the reference assembly version 4 of Williams82. Each point shows the $-\log_{10}(p)$ associated with a particular marker or $k$-mer. Horizontal dotted lines indicate the 5% family-wise error-rate significance threshold determined from a randomization approach. Vertical dotted lines indicate the position of signals associated with the trait. Documented loci are colored according to whether they were found (green) or not (red) by a particular approach. The "Gm" prefix has been left out of chromosome names for simplicity.

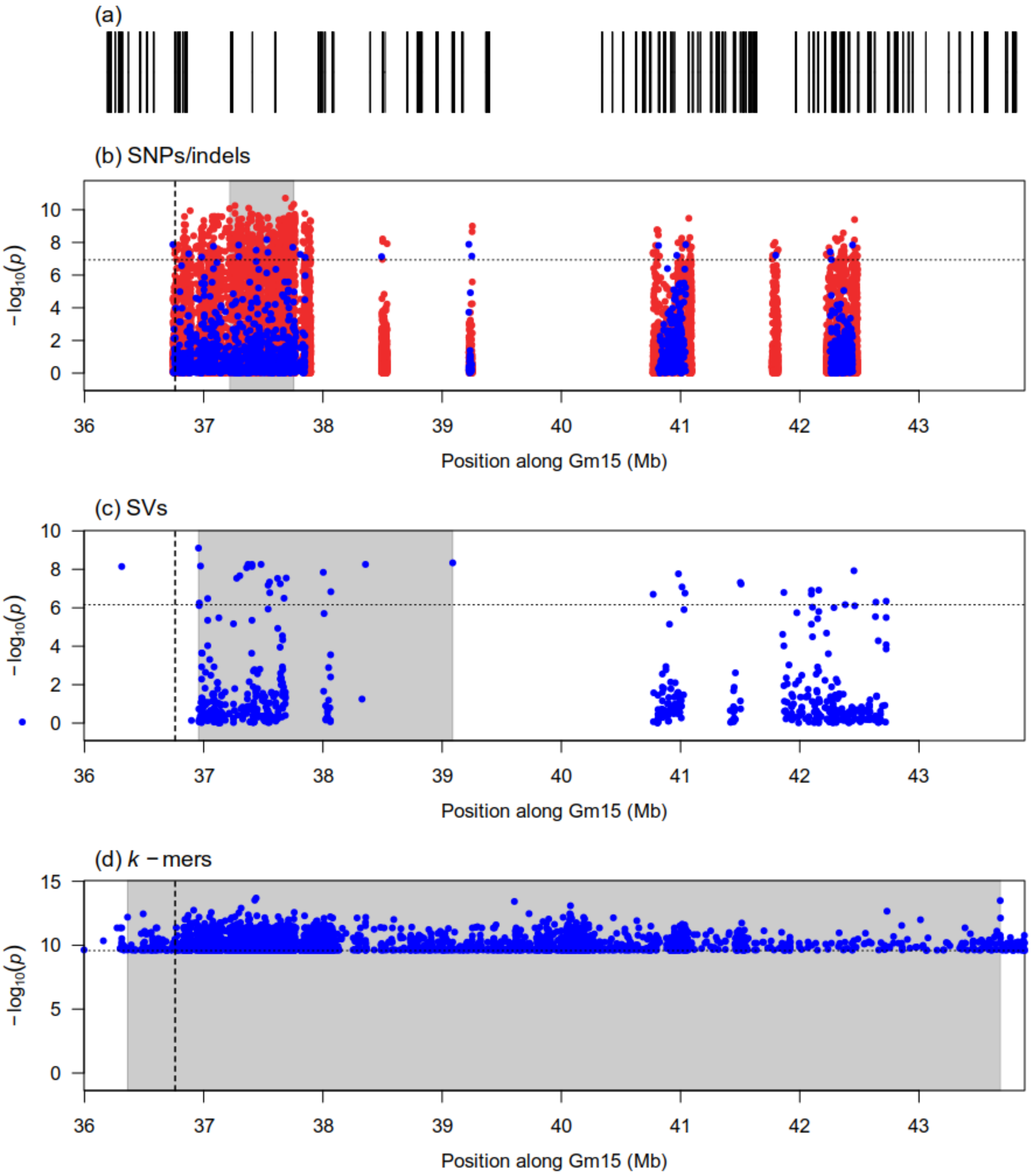


Figure S41: Zoomed-in Manhattan plots of signals detected by the GWAS analysis of corrected dry weight (resistance to *P. sojae*) at the *cdwGm15* locus using three genotype datasets: (b) Platypus (SNPs and indels), (c) Paragraph (SVs), (d) $k$-mers presence/absence. Panel (a) shows gene models over the genomic interval. Horizontal dotted lines indicate the 5% family-wise error-rate significance threshold determined from a randomization approach. Vertical dotted lines indicate the location of the Glyma.15g217100 gene suggested by de Ronne et al. (2022) as associated with the locus. Gray shaded rectangles indicate the region delimited by the top 5% (SVs) or top 1% (SNPs/indels and $k$-mers) associations in the signal region. In the case of SNPs/indels, blue points denote markers used in the original analysis, whereas red points denote markers that had originally been pruned but whose $p$-values were computed after signal discovery.

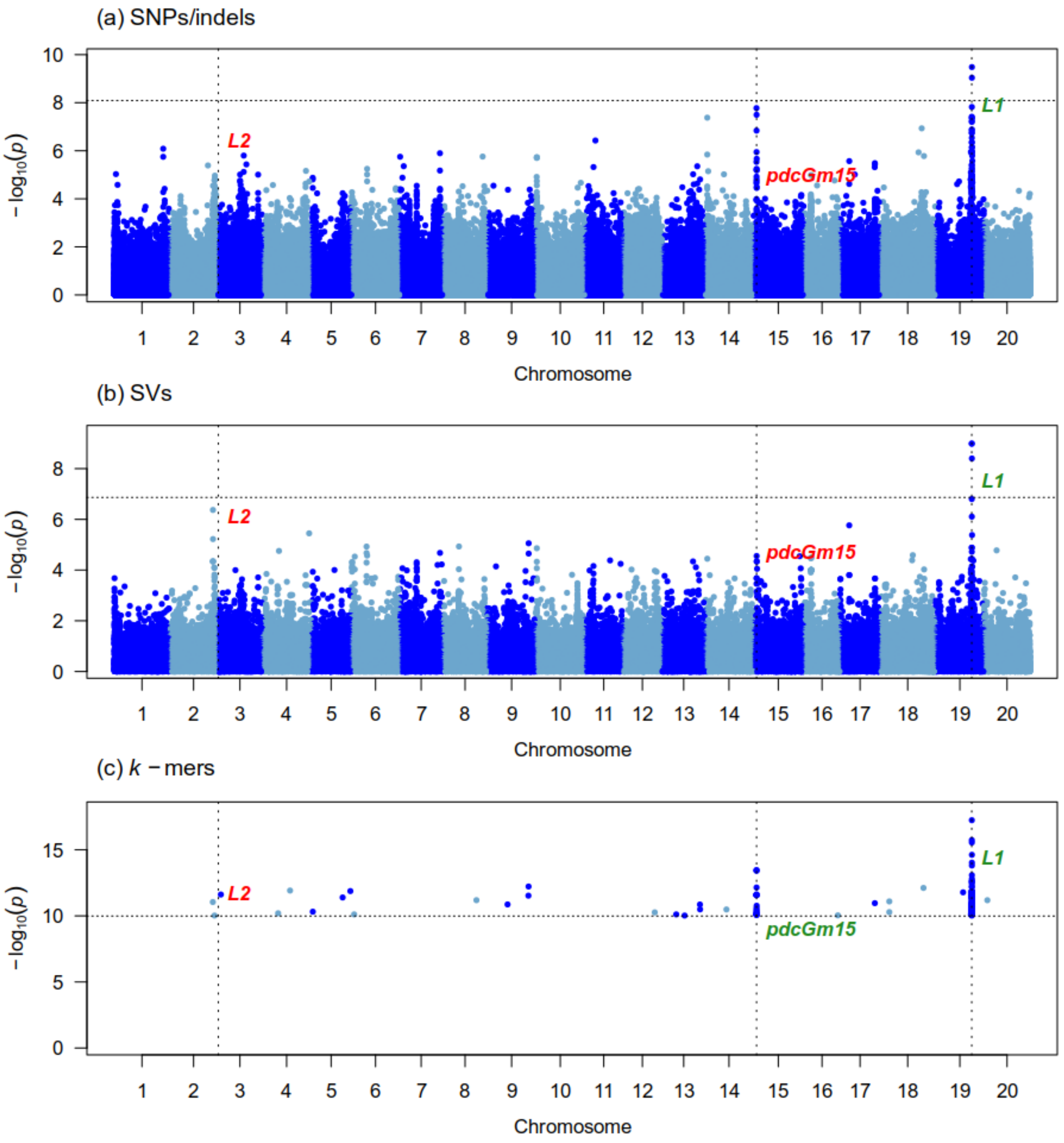


Figure S42: Manhattan plots generated from the GWAS analysis of pod color (second GWAS) on 259 samples using three genotype datasets : (a) Platypus (SNPs and indels), (b) Paragraph (SVs), (c) $k$-mers presence/absence. The x-axis shows the position along the reference assembly version 4 of Williams82. Each point shows the $-\log_{10}(p)$ associated with a particular marker or $k$-mer. Horizontal dotted lines indicate the 5% family-wise error-rate significance threshold determined from a randomization approach. Vertical dotted lines indicate the position of signals associated with the trait. Documented loci are colored according to whether they were found (green) or not (red) by a particular approach. The “Gm” prefix has been left out of chromosome names for simplicity.

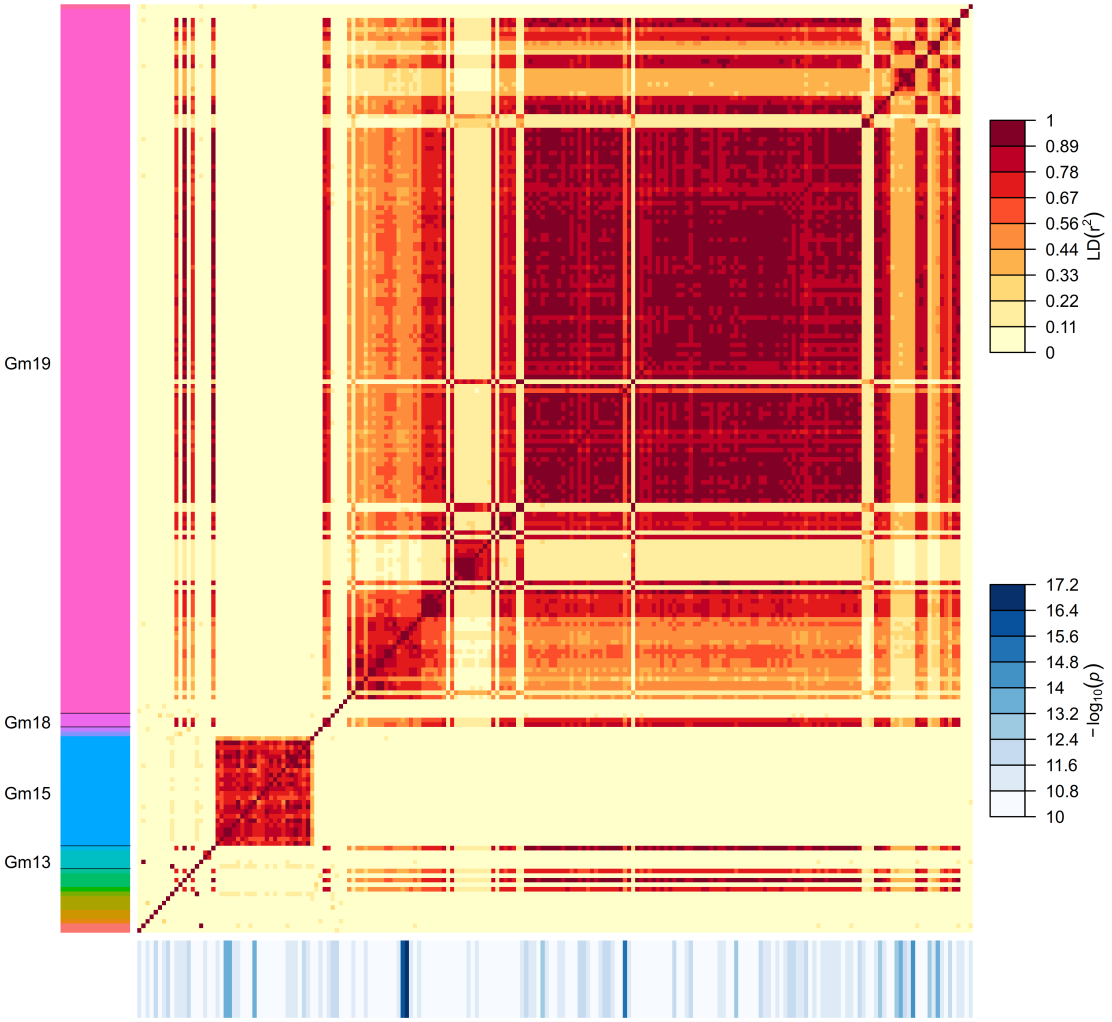


Figure S43: Pairwise LD among 203 significant $k$-mers identified for pod color (second GWAS). $k$-mers are sorted along the y-axis according to their putative position along the reference assembly version 4 of Williams82, as identified by "Gm" chromosome labels. Sequences that lack a "Gm" label (top of the y-axis) represent unanchored scaffolds. $k$-mers are represented in the same order along the x- and y-axis. The colored rectangles drawn below the x-axis represent the $-\log_{10}(p)$ of each $k$-mer.

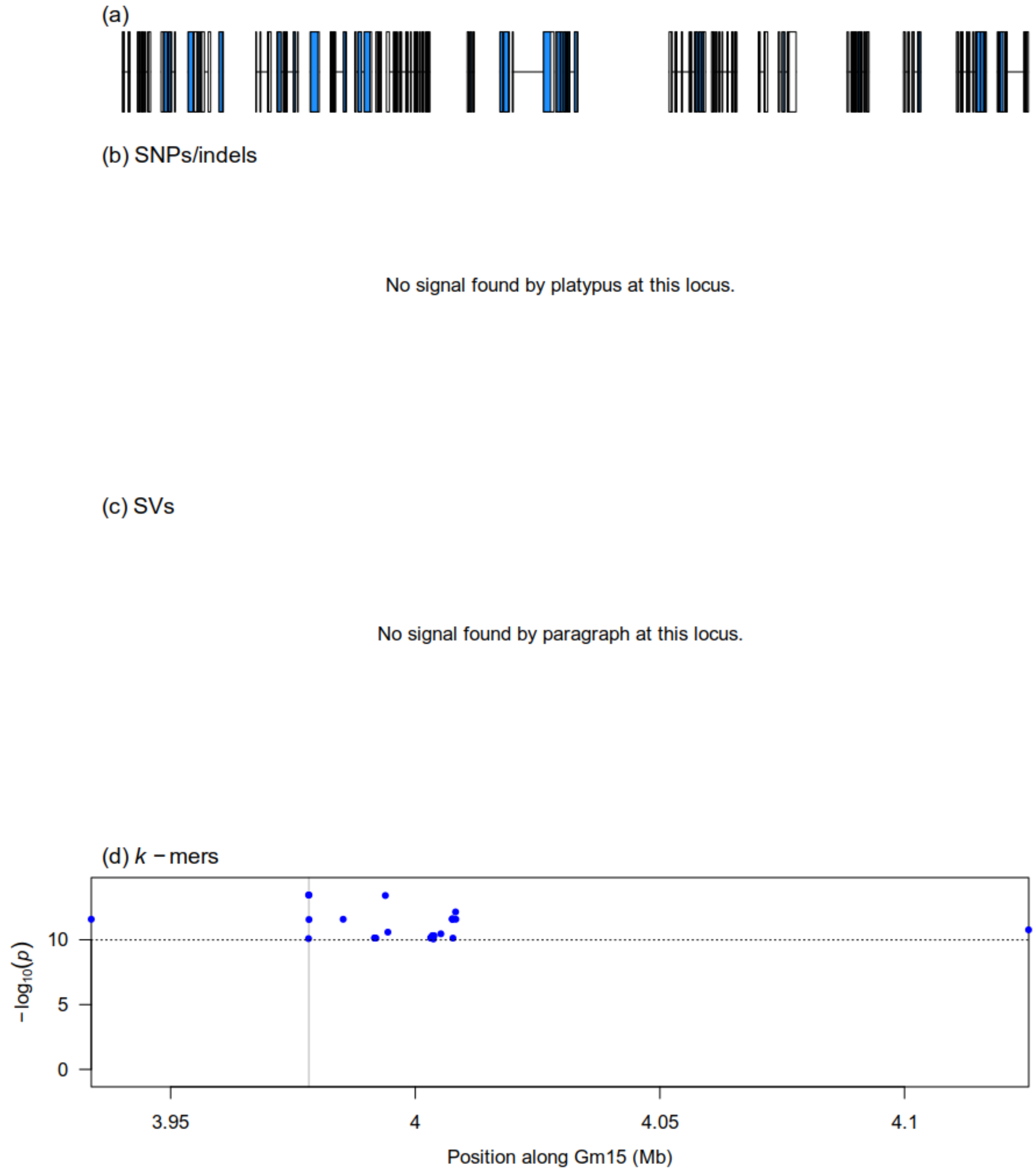


Figure S44: Zoomed-in Manhattan plots of signals detected by the GWAS analysis of pod color (second GWAS) at the newly suggested *pdcGm15* locus using three genotype datasets: (b) Platypus (SNPs and indels), (c) Paragraph (SVs), (d) $k$-mers presence/absence. Panel (a) shows gene models over the genomic interval. Horizontal dotted lines indicate the 5% family-wise error-rate significance threshold determined from a randomization approach. Gray shaded rectangles indicate the region delimited by the top 5% (SVs) or top 1% (SNPs/indels and $k$-mers) associations in the signal region. In the case of SNPs/indels, blue points denote markers used in the original analysis, whereas red points denote markers that had originally been pruned but whose $p$-values were computed after signal discovery.

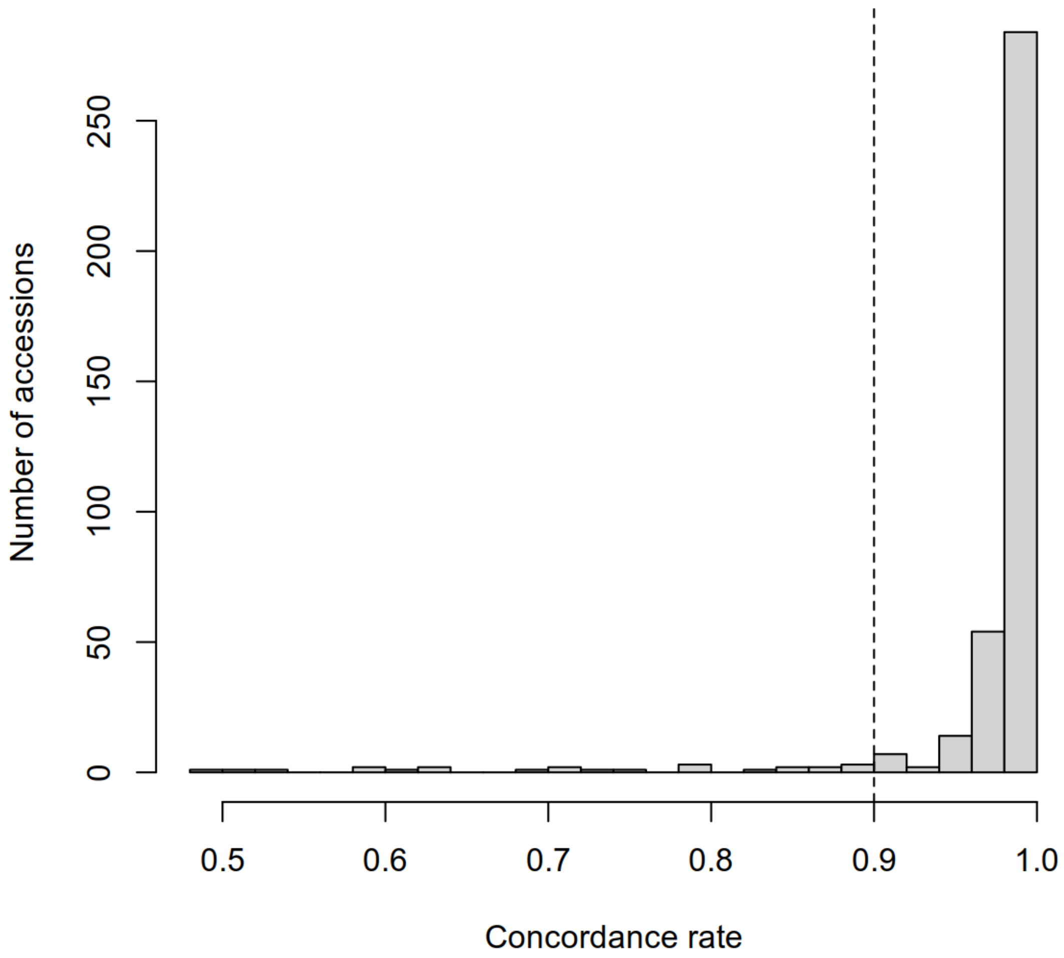


Figure S45: Histogram of concordance rates between genotypes derived from WGS data and SoySNP50K genotypes for 385 soybean accessions. The vertical line indicates the 0.9 concordance threshold that was used for filtering out mismatching samples.

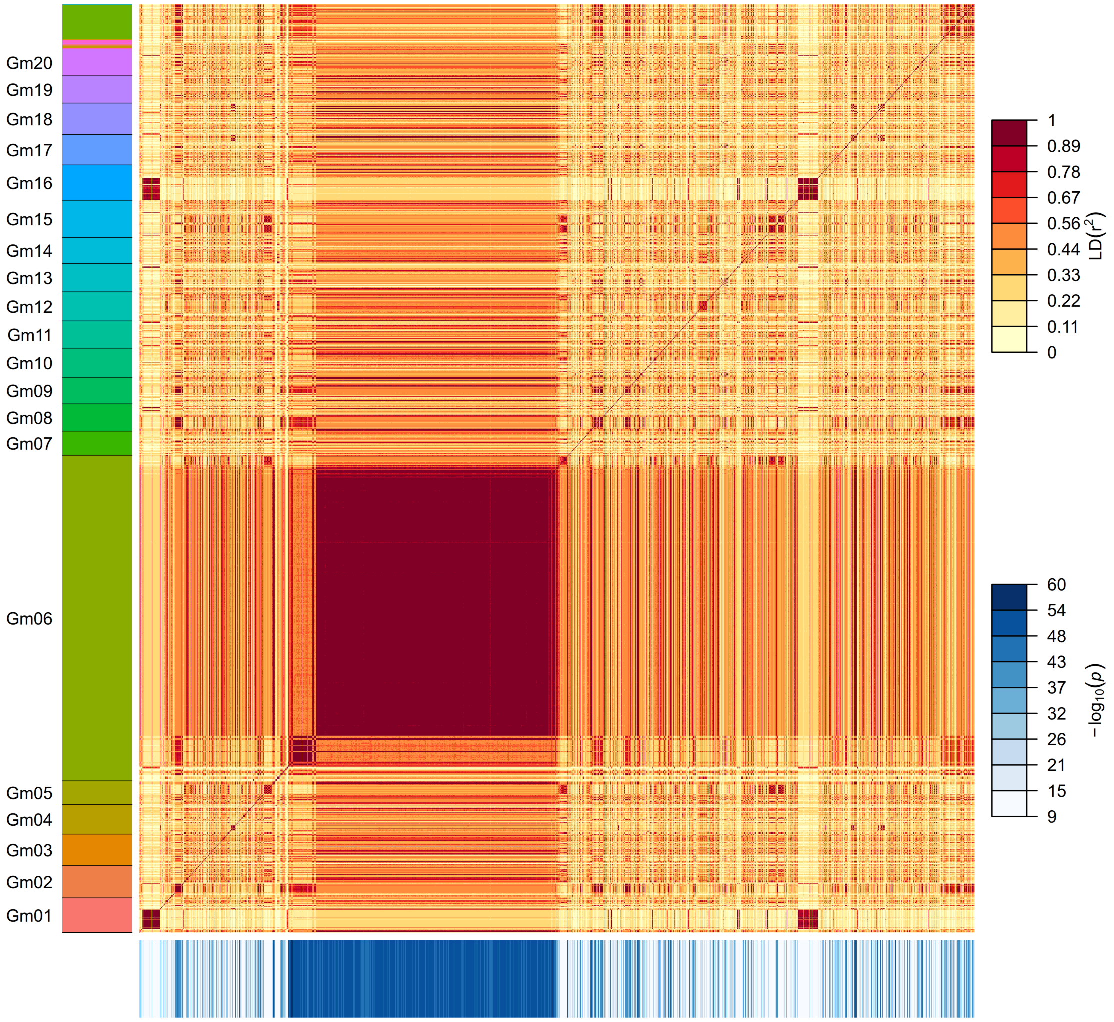


Figure S46: Pairwise LD among 1500 significant $k$-mers identified for pubescence color (first GWAS). $k$-mers are sorted along the y-axis according to their putative position along the reference assembly version 4 of Williams82, as identified by "Gm" chromosome labels. Sequences that lack a "Gm" label (top of the y-axis) represent unanchored scaffolds. $k$-mers are represented in the same order along the x- and y-axis. The colored rectangles drawn below the x-axis represent the $-\log_{10}(p)$ of each $k$-mer.

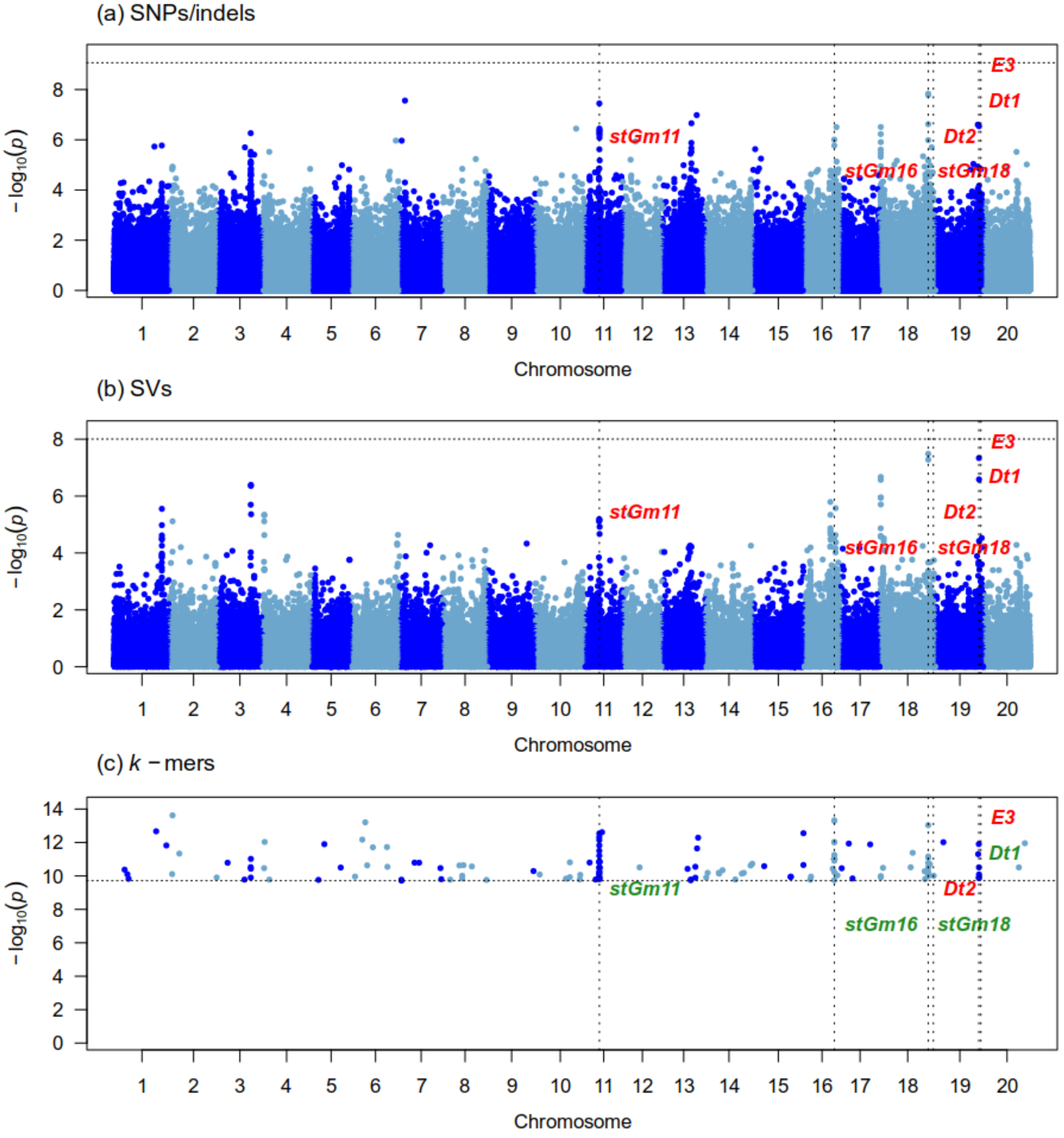


Figure S47: Manhattan plots generated from the GWAS analysis of stem termination type (second GWAS) on 260 samples using three genotype datasets : (a) Platypus (SNPs and indels), (b) Paragraph (SVs), (c) $k$-mers presence/absence. The x-axis shows the position along the reference assembly version 4 of Williams82. Each point shows the $-\log_{10}(p)$ associated with a particular marker or $k$-mer. Horizontal dotted lines indicate the 5% family-wise error-rate significance threshold determined from a randomization approach. Vertical dotted lines indicate the position of signals associated with the trait. Documented loci are colored according to whether they were found (green) or not (red) by a particular approach. The "Gm" prefix has been left out of chromosome names for simplicity.

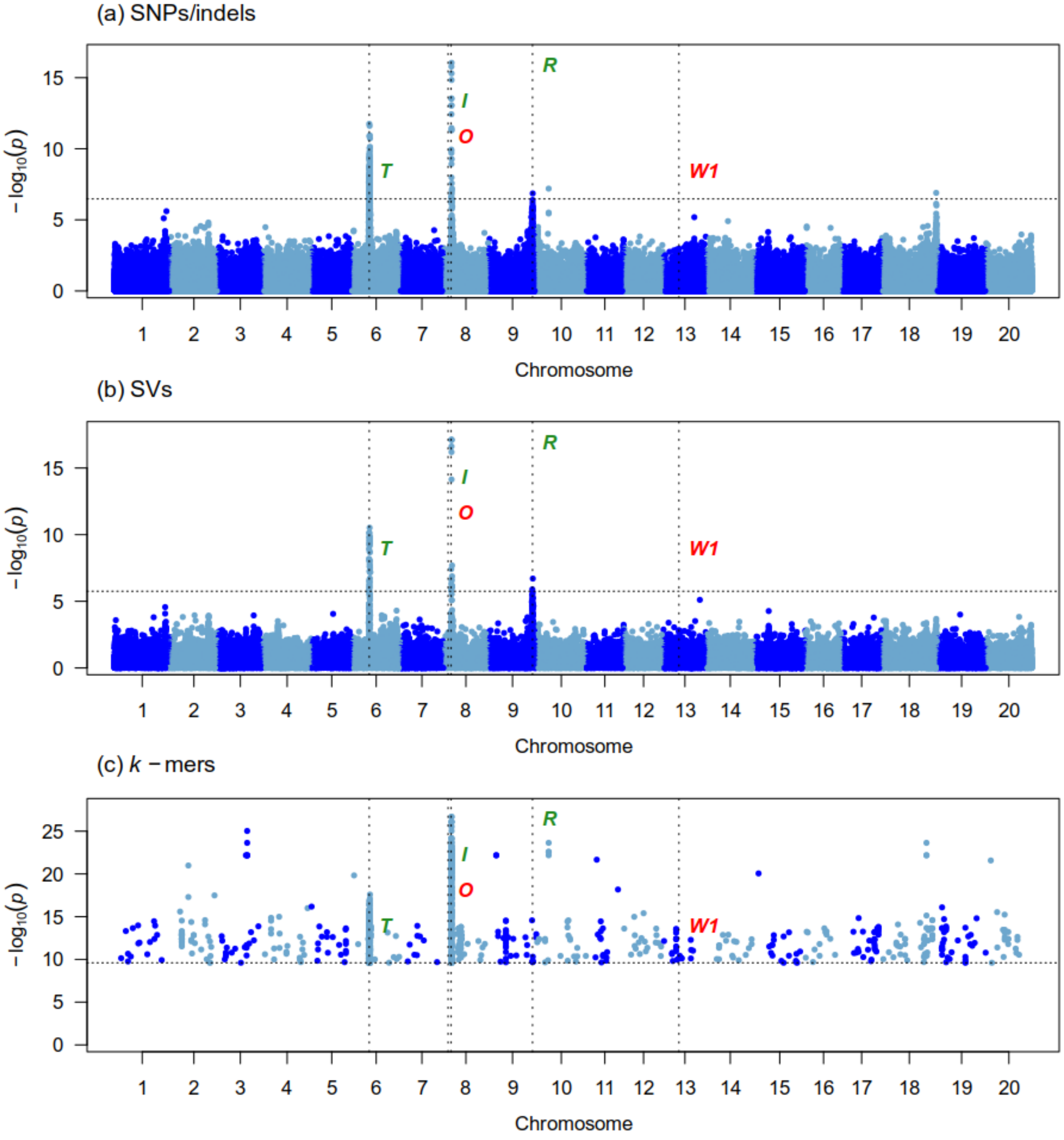


Figure S48: Manhattan plots generated from the GWAS analysis of hilum color (first GWAS) on 326 samples using three genotype datasets : (a) Platypus (SNPs and indels), (b) Paragraph (SVs), (c) $k$-mers presence/absence. The x-axis shows the position along the reference assembly version 4 of Williams82. Each point shows the $-\log_{10}(p)$ associated with a particular marker or $k$-mer. Horizontal dotted lines indicate the 5% family-wise error-rate significance threshold determined from a randomization approach. Vertical dotted lines indicate the position of signals associated with the trait. Documented loci are colored according to whether they were found (green) or not (red) by a particular approach. The "Gm" prefix has been left out of chromosome names for simplicity.

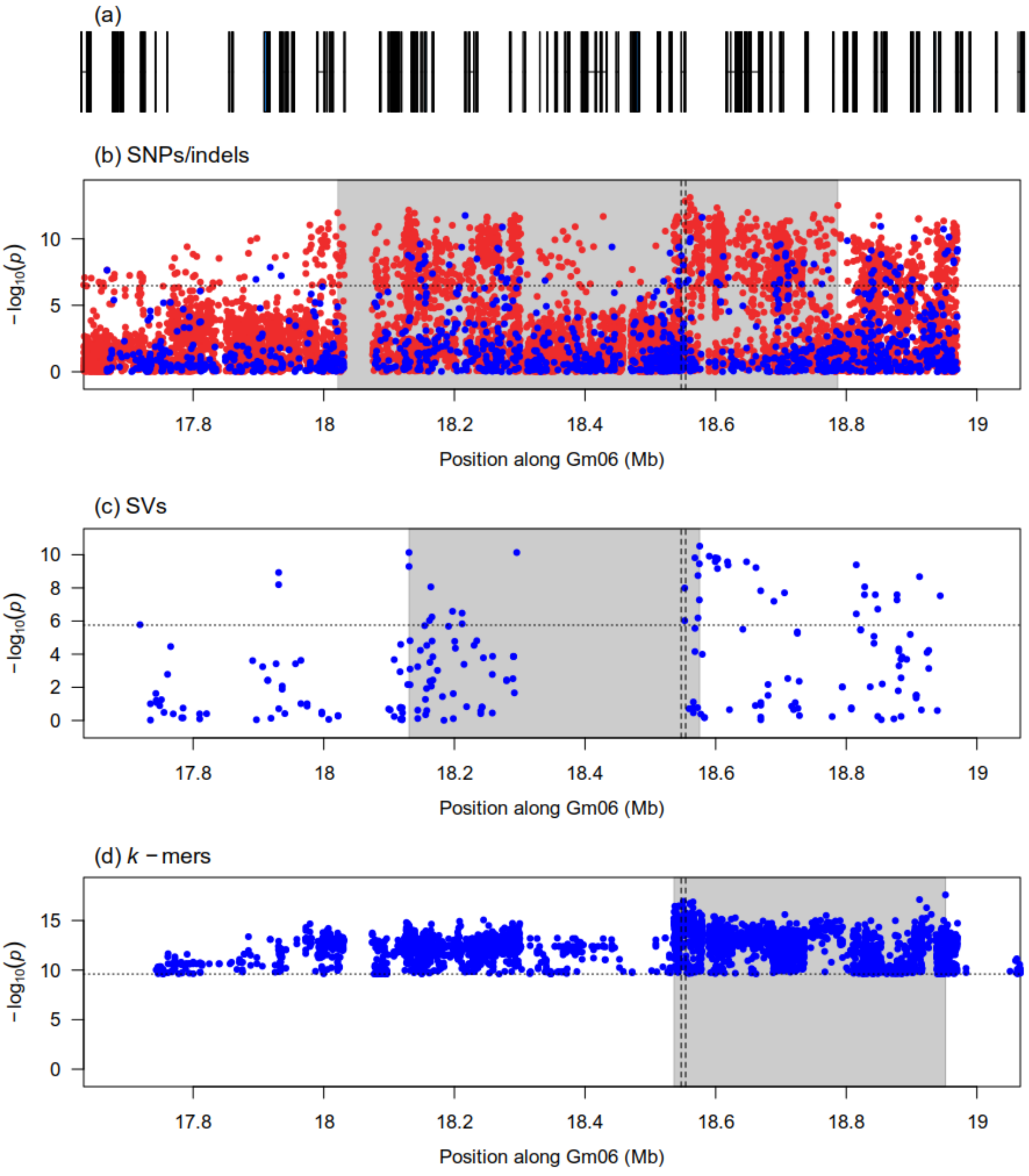


Figure S49: Zoomed-in Manhattan plots of signals detected by the GWAS analysis of hilum color (first GWAS) at the $T$ locus using three genotype datasets: (b) Platypus (SNPs and indels), (c) Paragraph (SVs), (d) $k$-mers presence/absence. Panel (a) shows gene models over the genomic interval. Horizontal dotted lines indicate the 5% family-wise error-rate significance threshold determined from a randomization approach. Vertical dotted lines indicate the location of the Glyma.06g202300 gene associated with the locus. Gray shaded rectangles indicate the region delimited by the top 5% (SVs) or top 1% (SNPs/indels and $k$-mers) associations in the signal region. In the case of SNPs/indels, blue points denote markers used in the original analysis, whereas red points denote markers that had originally been pruned but whose $p$-values were computed after signal discovery.

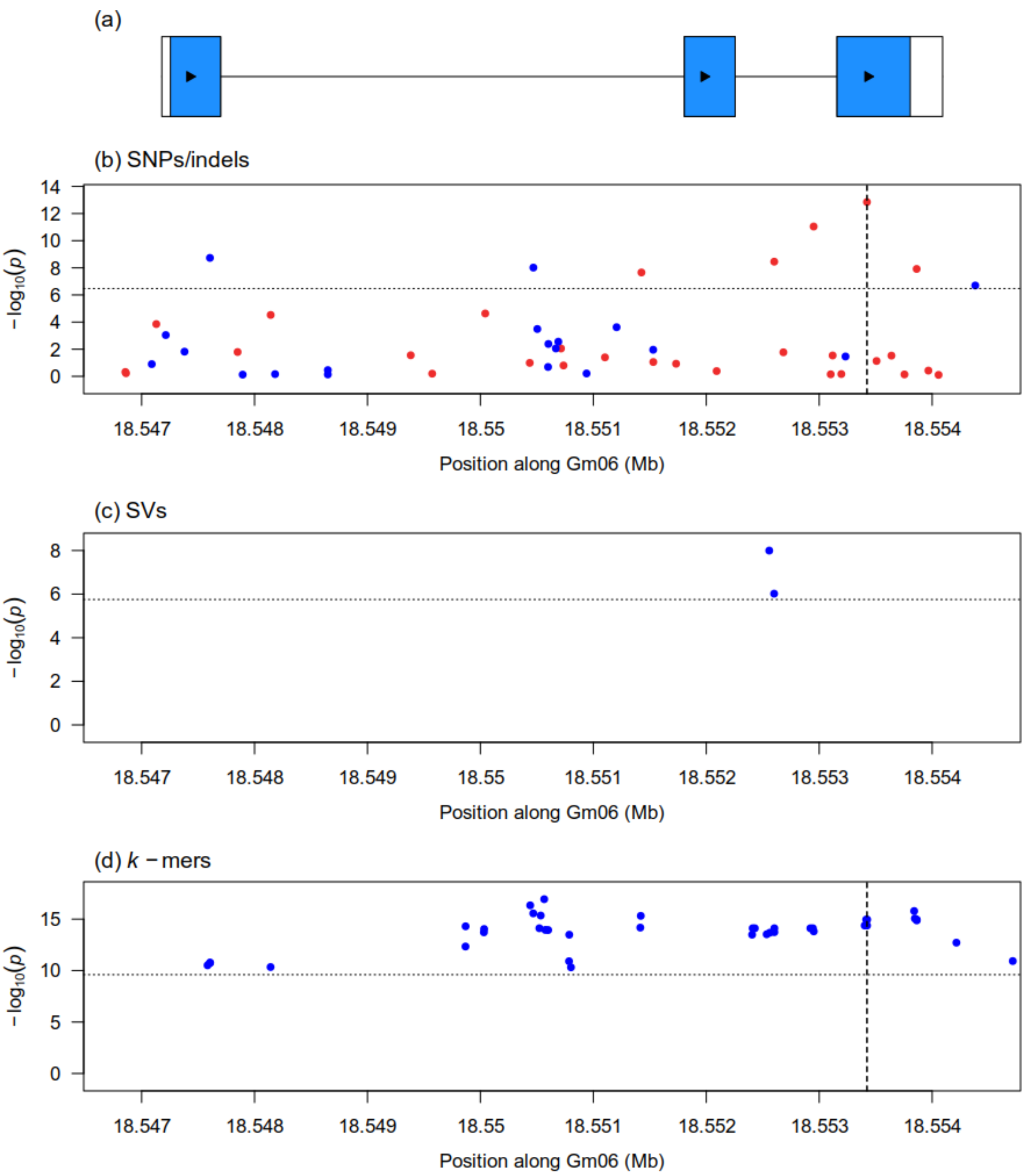


Figure S50: Zoomed-in Manhattan plots generated from the GWAS analysis of hilum color (first GWAS) at the Glyma.06g202300 gene associated with the $T$ locus using three genotype datasets: (b) Platypus (SNPs and indels), (c) Paragraph (SVs), (d) $k$-mers presence/absence. Vertical dotted lines in panels (b) and (d) indicate the location of the causal indel at this locus. Horizontal dotted lines indicate the 5% family-wise error-rate significance threshold determined from a randomization approach. In the case of SNPs/indels, blue points denote markers used in the original analysis, whereas red points denote markers that had originally been pruned but whose $p$-values were computed after signal discovery. Panel (a) shows gene models over the plotting interval. Exons are represented by rectangles whereas introns are represented by horizontal lines. Coding sequences are shown in blue and the direction of transcription is indicated by arrows.

(a)

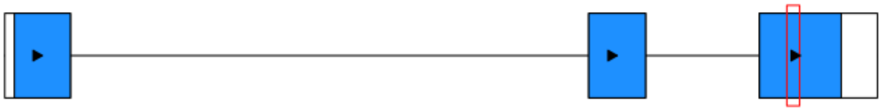

(b)

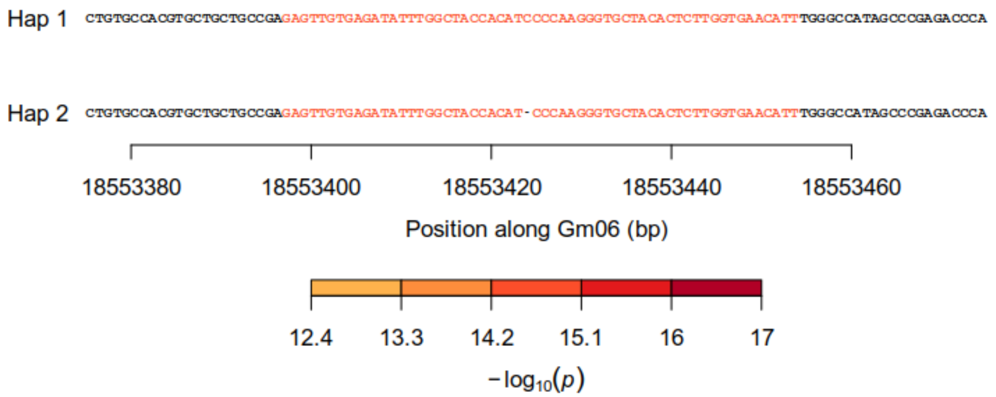


(c)

| | Hap 1 | Hap 2 |
|---|---|---|
| Black | 95 | 0 |
| Brown | 61 | 0 |
| Buff | 17 | 81 |
| Gray | 3 | 2 |
| Imperfect black | 0 | 20 |
| Reddish brown | 13 | 0 |
| Yellow | 3 | 22 |

Figure S51: Identification of a causal indel underlying significant $k$-mers at the Glyma.06g202300 gene associated with the $T$ locus for hilum color (first GWAS). (a) Gene model of Glyma.06g202300. Exons are represented by rectangles whereas introns are represented by horizontal lines. Coding sequences are shown in blue and the direction of transcription is indicated by arrows. The red rectangle identifies the region that is zoomed-in in panel (b). (b) Nucleotide sequences of haplotypes observed in at least five samples across the dataset. Individual nucleotides are colored according to the $-\log_{10}(p)$ of the most significant $k$-mer overlapping them. Dashes indicate gaps in haplotype sequence alignment whereas vertical lines indicate differences in sequence between two haplotypes. (c) Contingency table of the phenotypes and haplotypes observed in the dataset. Haplotypes correspond to those shown in panel (b).

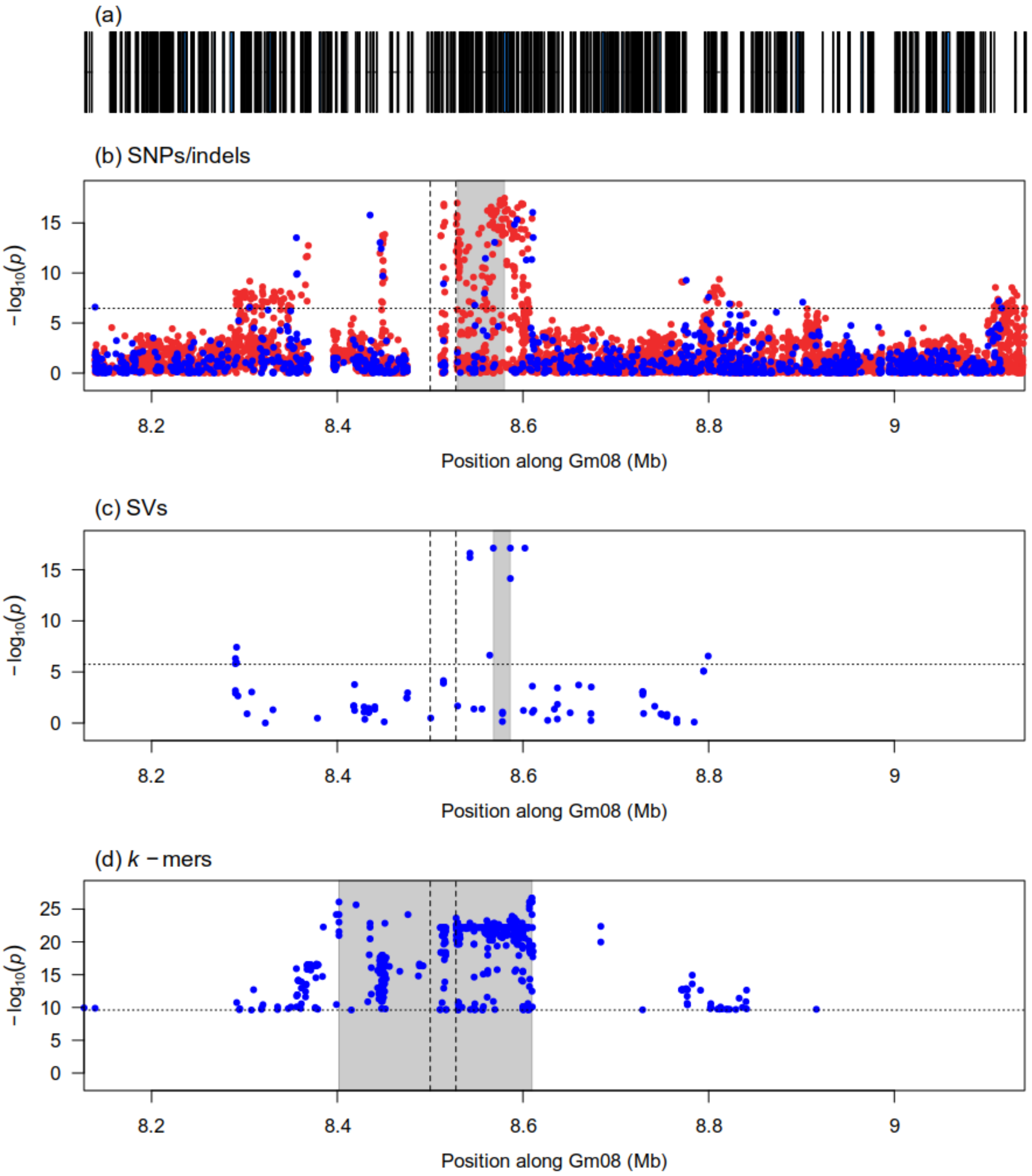


Figure S52: Zoomed-in Manhattan plots of signals detected by the GWAS analysis of hilum color (first GWAS) at the *I* locus using three genotype datasets: (b) Platypus (SNPs and indels), (c) Paragraph (SVs), (d) $k$-mers presence/absence. Panel (a) shows gene models over the genomic interval. Horizontal dotted lines indicate the 5% family-wise error-rate significance threshold determined from a randomization approach. Vertical dotted lines indicate the boundaries of the tandem duplication/inversion identified as the causal variant at this locus. Gray shaded rectangles indicate the region delimited by the top 5% (SVs) or top 1% (SNPs/indels and $k$-mers) associations in the signal region. In the case of SNPs/indels, blue points denote markers used in the original analysis, whereas red points denote markers that had originally been pruned but whose $p$-values were computed after signal discovery.

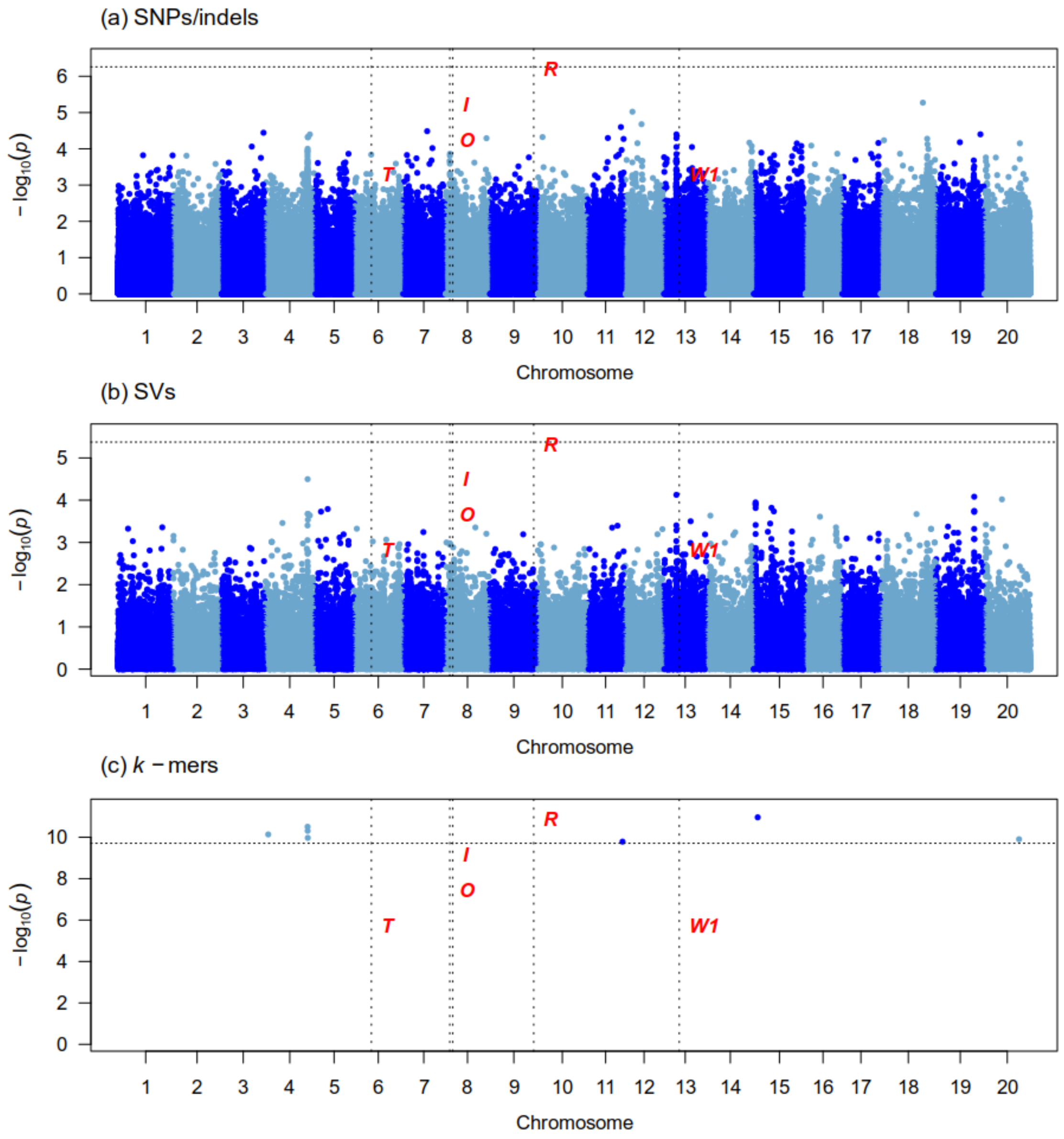


Figure S53: Manhattan plots generated from the GWAS analysis of hilum color (third GWAS) on 76 samples using three genotype datasets : (a) Platypus (SNPs and indels), (b) Paragraph (SVs), (c) $k$-mers presence/absence. The x-axis shows the position along the reference assembly version 4 of Williams82. Each point shows the $-\log_{10}(p)$ associated with a particular marker or $k$-mer. Horizontal dotted lines indicate the 5% family-wise error-rate significance threshold determined from a randomization approach. Vertical dotted lines indicate the position of signals associated with the trait. Documented loci are colored according to whether they were found (green) or not (red) by a particular approach. The "Gm" prefix has been left out of chromosome names for simplicity.

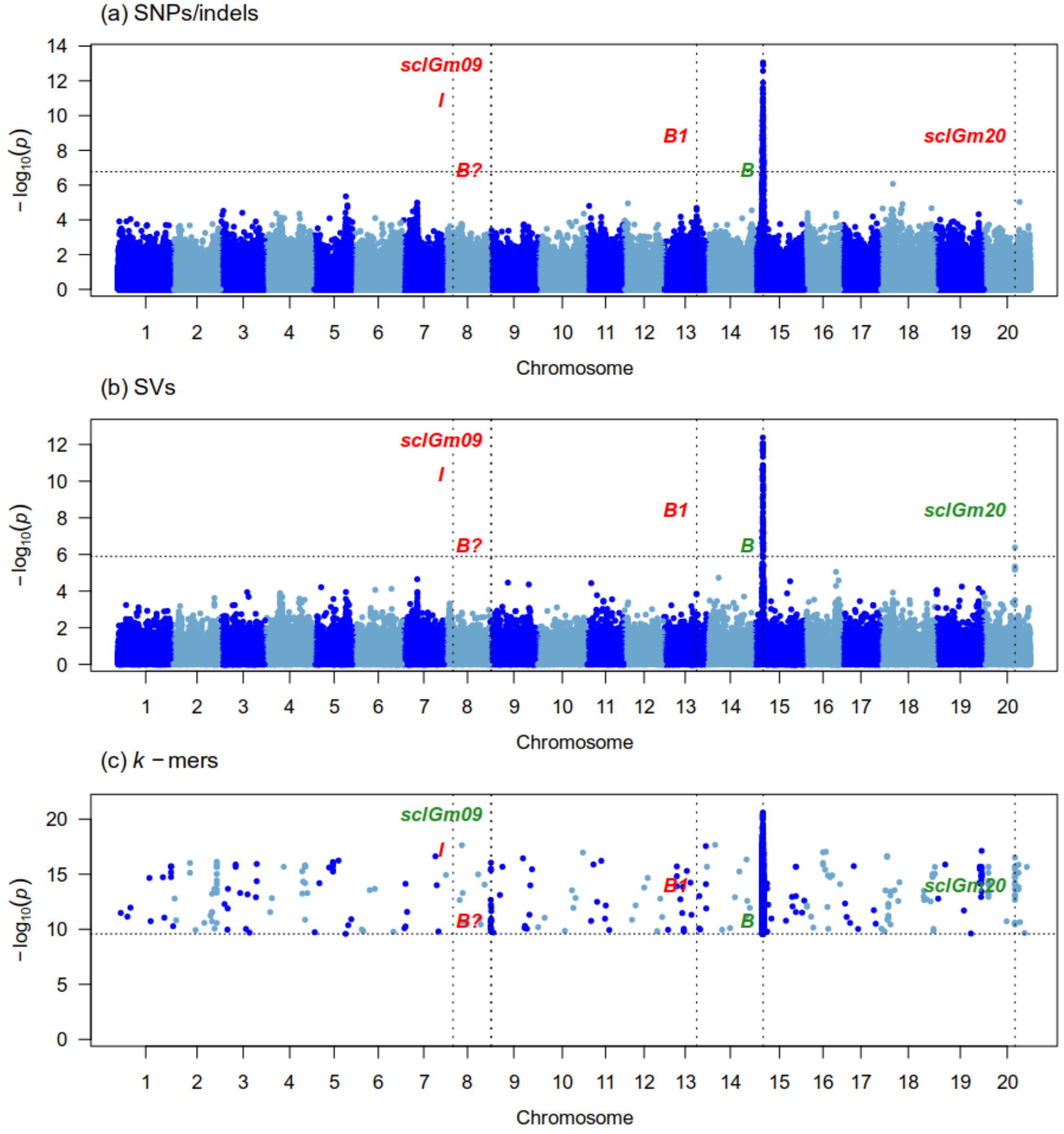


Figure S54: Manhattan plots generated from the GWAS analysis of seed coat luster (first GWAS) on 347 samples using three genotype datasets : (a) Platypus (SNPs and indels), (b) Paragraph (SVs), (c) $k$-mers presence/absence. The x-axis shows the position along the reference assembly version 4 of Williams82. Each point shows the $-\log_{10}(p)$ associated with a particular marker or $k$-mer. Horizontal dotted lines indicate the 5% family-wise error-rate significance threshold determined from a randomization approach. Vertical dotted lines indicate the position of signals associated with the trait. Documented loci are colored according to whether they were found (green) or not (red) by a particular approach. The "Gm" prefix has been left out of chromosome names for simplicity.

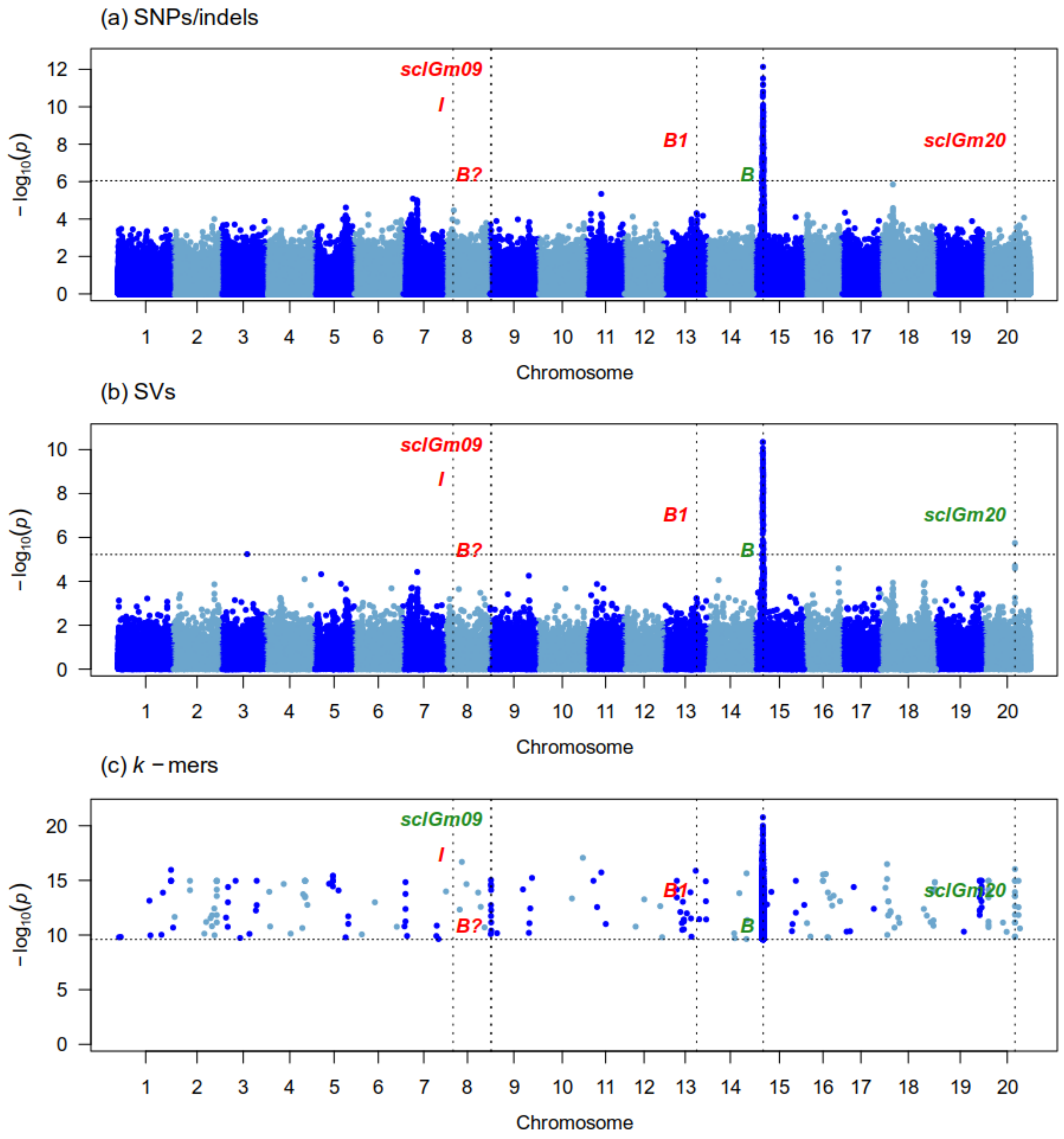


Figure S55: Manhattan plots generated from the GWAS analysis of seed coat luster (second GWAS) on 170 samples using three genotype datasets : (a) Platypus (SNPs and indels), (b) Paragraph (SVs), (c) $k$-mers presence/absence. The x-axis shows the position along the reference assembly version 4 of Williams82. Each point shows the $-\log_{10}(p)$ associated with a particular marker or $k$-mer. Horizontal dotted lines indicate the 5% family-wise error-rate significance threshold determined from a randomization approach. Vertical dotted lines indicate the position of signals associated with the trait. Documented loci are colored according to whether they were found (green) or not (red) by a particular approach. The "Gm" prefix has been left out of chromosome names for simplicity.

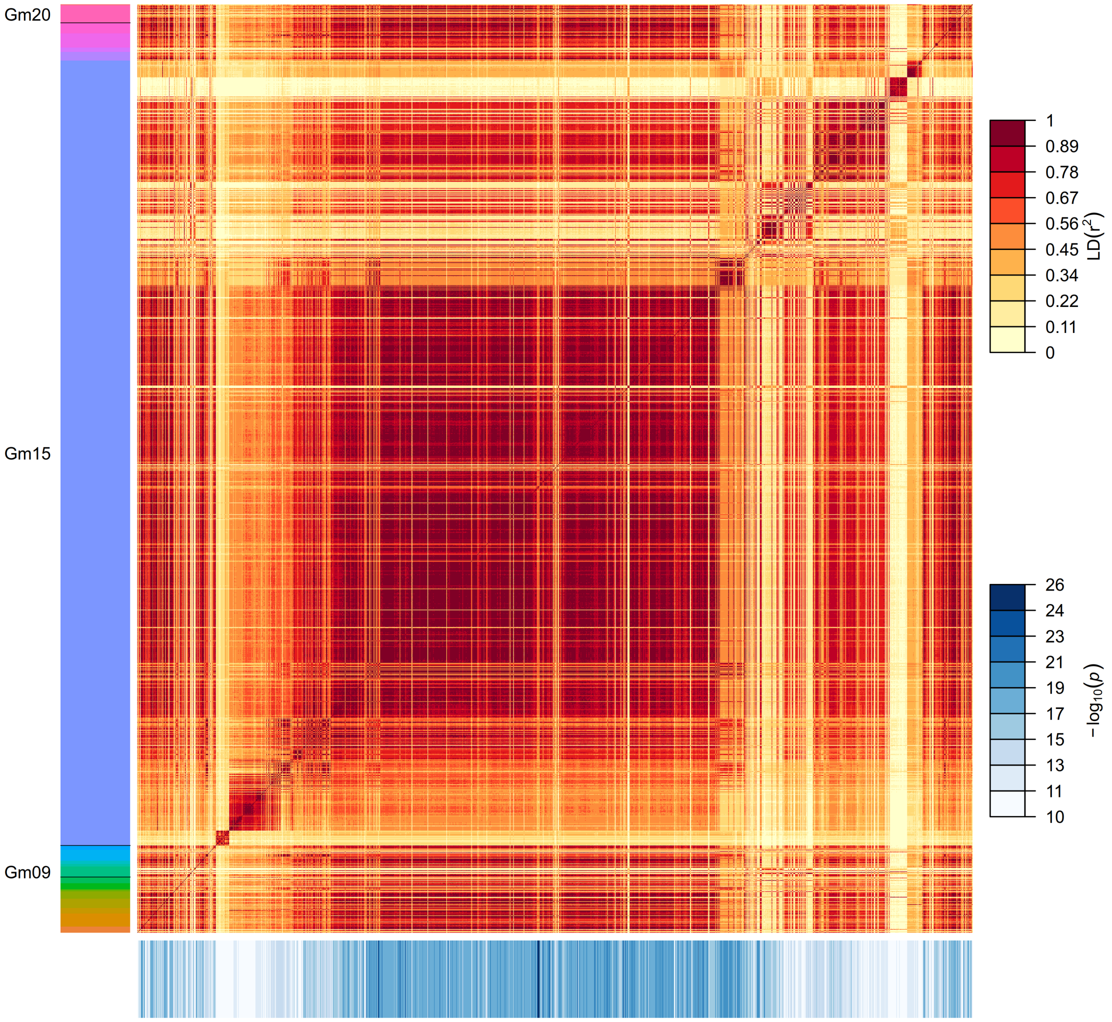


Figure S56: Pairwise LD among 1500 significant $k$-mers identified for seed coat luster (third GWAS). $k$-mers are sorted along the y-axis according to their putative position along the reference assembly version 4 of Williams82, as identified by “Gm” chromosome labels. Sequences that lack a “Gm” label (top of the y-axis) represent unanchored scaffolds. $k$-mers are represented in the same order along the x- and y-axis. The colored rectangles drawn below the x-axis represent the $-\log_{10}(p)$ of each $k$-mer.

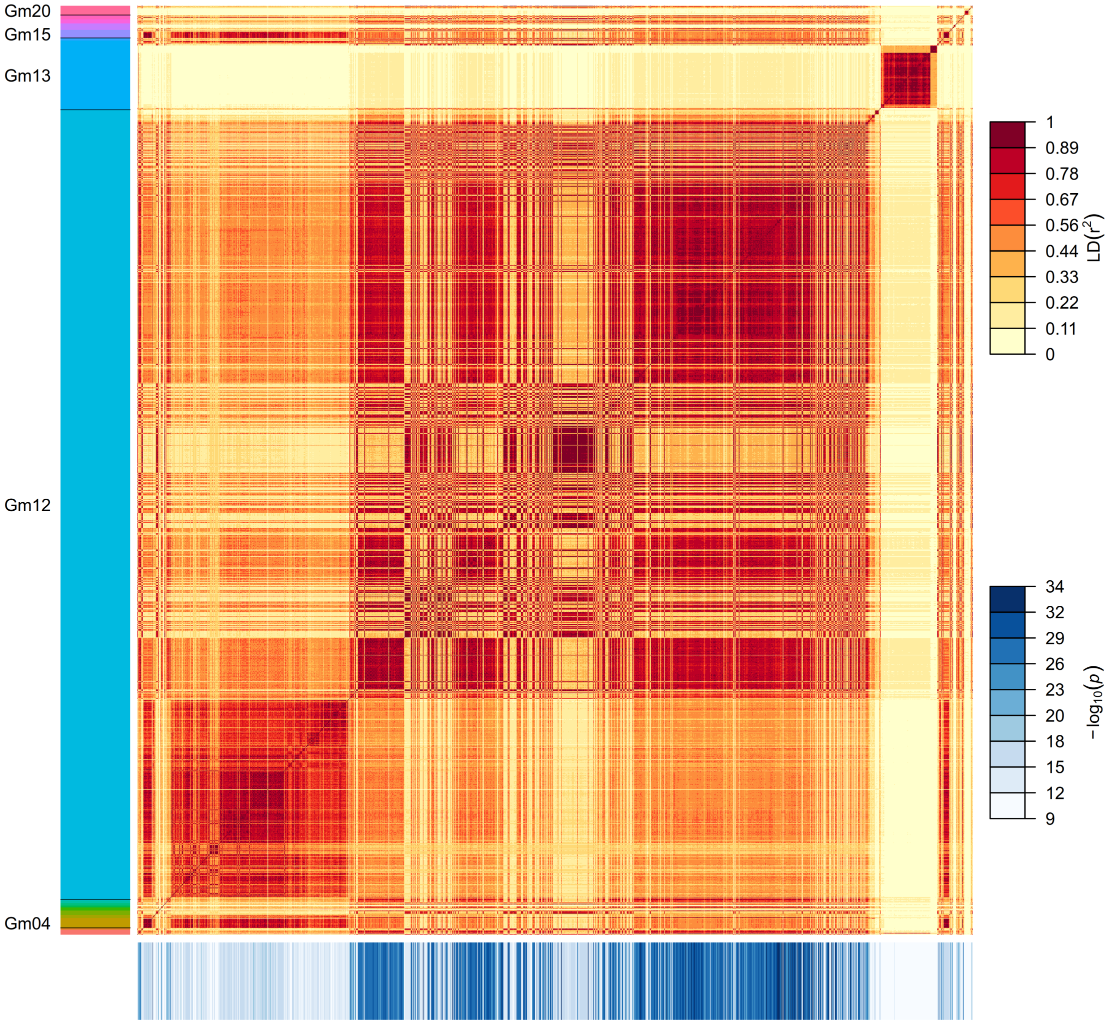


Figure S57: Pairwise LD among 1500 significant $k$-mers identified for pubescence form (first GWAS). $k$-mers are sorted along the y-axis according to their putative position along the reference assembly version 4 of Williams82, as identified by "Gm" chromosome labels. Sequences that lack a "Gm" label (top of the y-axis) represent unanchored scaffolds. $k$-mers are represented in the same order along the x- and y-axis. The colored rectangles drawn below the x-axis represent the $-\log_{10}(p)$ of each $k$-mer.

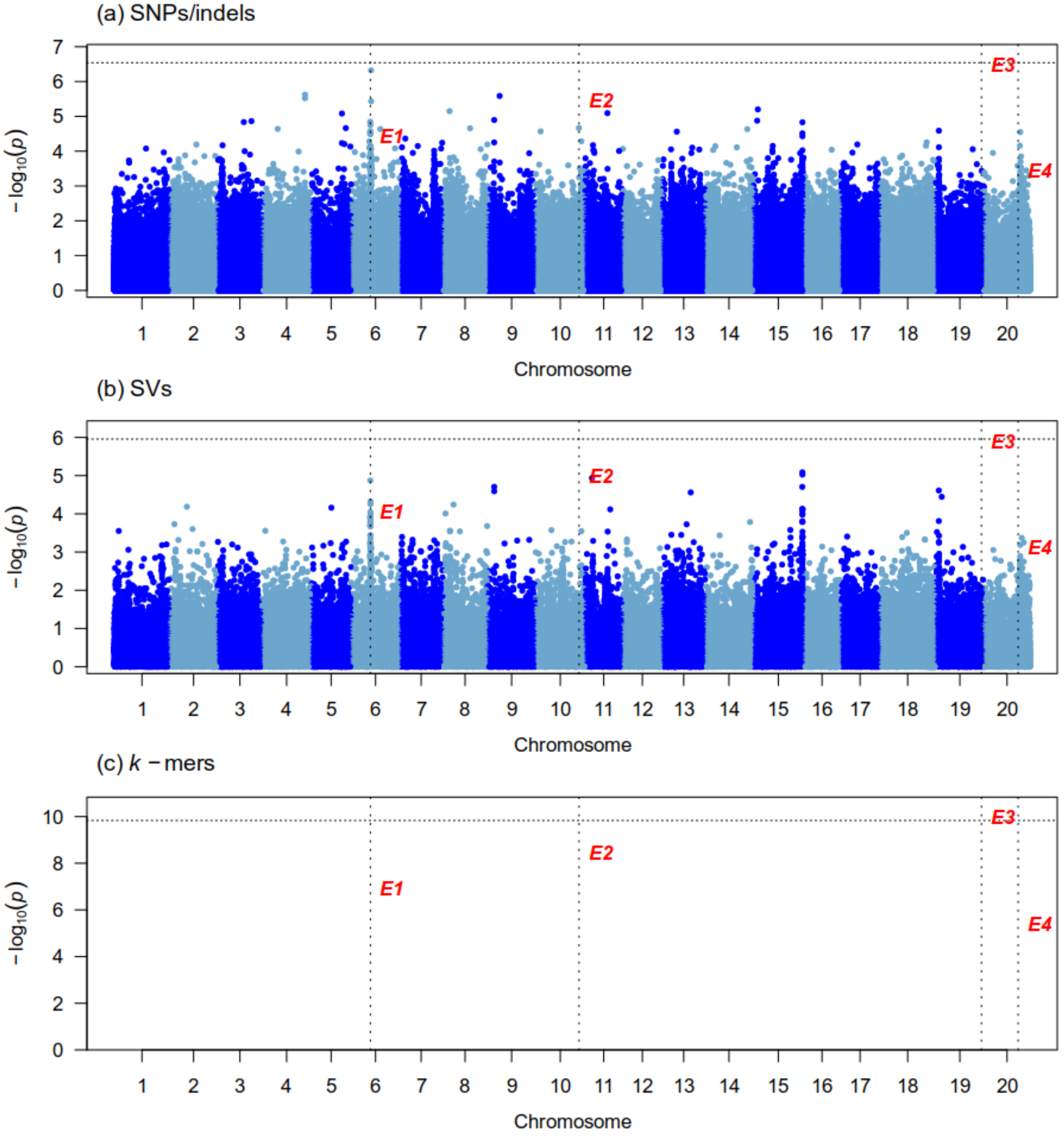


Figure S58: Manhattan plots generated from the GWAS analysis of maturity group on 348 samples using three genotype datasets : (a) Platypus (SNPs and indels), (b) Paragraph (SVs), (c) $k$-mers presence/absence. The x-axis shows the position along the reference assembly version 4 of Williams82. Each point shows the $-\log_{10}(p)$ associated with a particular marker or $k$-mer. Horizontal dotted lines indicate the 5% family-wise error-rate significance threshold determined from a randomization approach. Vertical dotted lines indicate the position of signals associated with the trait. Documented loci are colored according to whether they were found (green) or not (red) by a particular approach. The “Gm” prefix has been left out of chromosome names for simplicity.

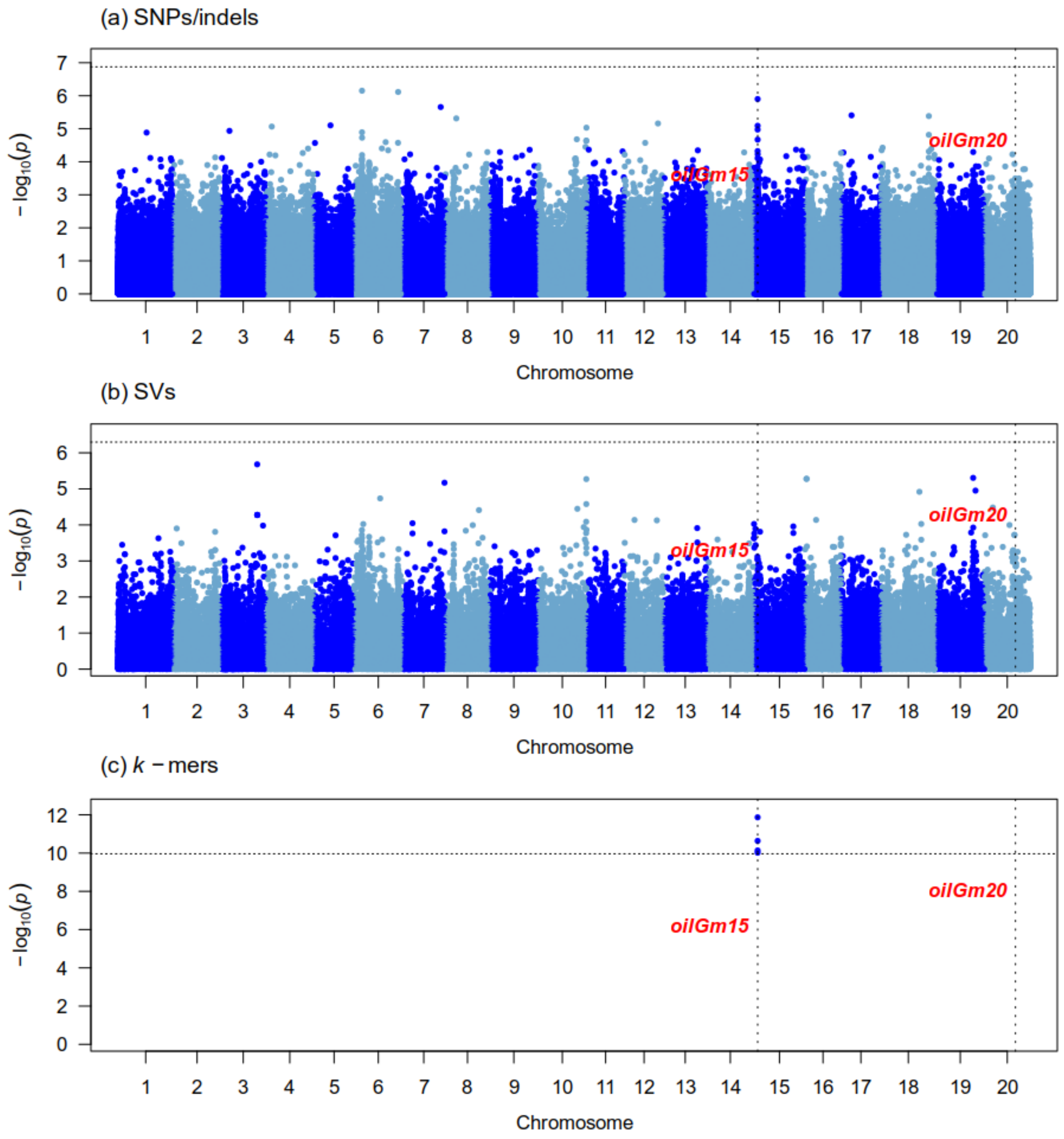


Figure S59: Manhattan plots generated from the GWAS analysis of seed oil content on 342 samples using three genotype datasets : (a) Platypus (SNPs and indels), (b) Paragraph (SVs), (c) $k$-mers presence/absence. The x-axis shows the position along the reference assembly version 4 of Williams82. Each point shows the $-\log_{10}(p)$ associated with a particular marker or $k$-mer. Horizontal dotted lines indicate the 5% family-wise error-rate significance threshold determined from a randomization approach. Vertical dotted lines indicate the position of signals associated with the trait. Documented loci are colored according to whether they were found (green) or not (red) by a particular approach. The "Gm" prefix has been left out of chromosome names for simplicity.

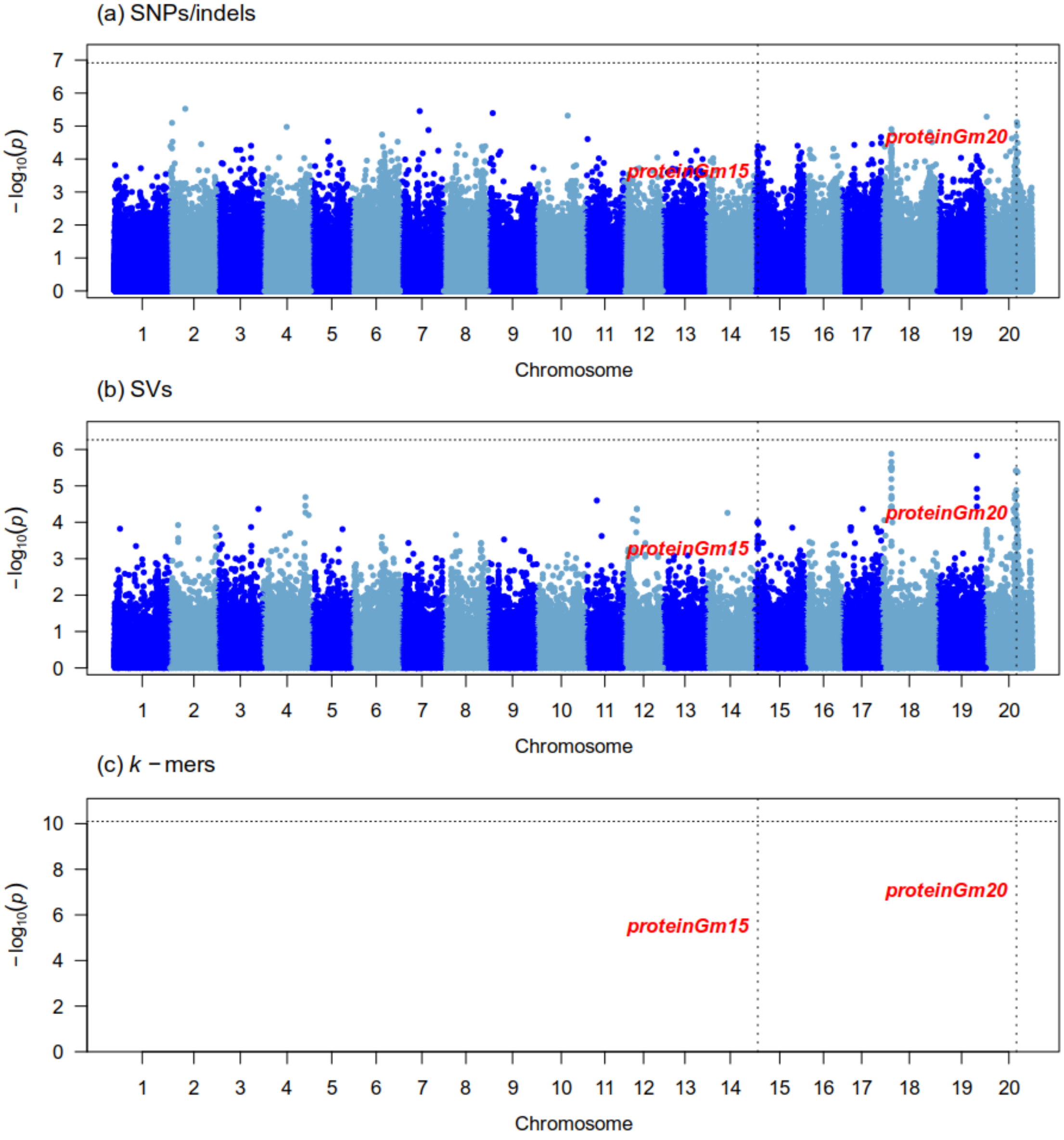


Figure S60: Manhattan plots generated from the GWAS analysis of seed protein content on 342 samples using three genotype datasets : (a) Platypus (SNPs and indels), (b) Paragraph (SVs), (c) $k$-mers presence/absence. The x-axis shows the position along the reference assembly version 4 of Williams82. Each point shows the $-\log_{10}(p)$ associated with a particular marker or $k$-mer. Horizontal dotted lines indicate the 5% family-wise error-rate significance threshold determined from a randomization approach. Vertical dotted lines indicate the position of signals associated with the trait. Documented loci are colored according to whether they were found (green) or not (red) by a particular approach. The "Gm" prefix has been left out of chromosome names for simplicity.
